# Mechanistic insights into CFTR function from molecular dynamics analysis of electrostatic interactions

**DOI:** 10.1101/2025.11.23.690003

**Authors:** Ahmad Elbahnsi, Jean-Paul Mornon, Isabelle Callebaut

## Abstract

The Cystic Fibrosis Transmembrane Conductance Regulator (CFTR) is an ATP-gated anion channel whose function is tightly linked to its conformational dynamics and is influenced by the composition of its membrane lipid environment. Despite high-resolution three-dimensional (3D) structures, the molecular determinants that stabilize specific CFTR conformations and enable ion conduction remain incompletely understood. Here, we performed all-atom molecular dynamics (MD) simulations of the human CFTR 3D structure in both the apo and VX-770 (ivacaftor)–bound states, embedded in a heterogeneous lipid bilayer, in order to systematically analyze electrostatic interactions, linking amino acids to each other as well as to anions and membrane lipids.

We identified 557 electrostatic interactions between charged and polar amino acid side chains, which we systematically mapped across the CFTR 3D structure. They are organized into specific regions, with a subset showing high frequency and conservation across simulations, suggesting a structural role in stabilizing CFTR architecture. In contrast, more transient electrostatic interactions were detected in dynamic regions potentially linked to conformational transitions or other functional roles. Irregularities in transmembrane (TM) helices often incorporate amino acids involved in electrostatic interactions. Many basic and polar residues involved in electrostatic interactions also engaged in anion coordination, underscoring their contribution to ion conduction. In addition, some showed selective interactions with cholesterol and phosphatidylserine, revealing spatially organized lipid binding, particularly at the level of the lasso and in the vicinity of the VX-770 binding site, which may mark regions important for allosteric communication. VX-770 binding preserved the global architecture of the electrostatic interaction networks but induced subtle shifts, acting on specific salt bridges. Regardless of whether VX-770 is present or not, a secondary portal displayed between TM10/TM12 emerged from these MD simulations, in addition to the main TM4/TM6 portal, whose morphology and diameter is controlled by a fluctuating salt bridge. Two exit routes also appeared for the exit of anions towards the extracellular milieu. Altogether, our integrative analysis highlights how dynamic electrostatic networks, together with ion and lipid interactions, support CFTR’s structural plasticity and functional modulation, offering molecular insights into potentiation mechanisms and into the specific evolution of CFTR in the ABC transporter superfamily.

**TOC Graphic:** 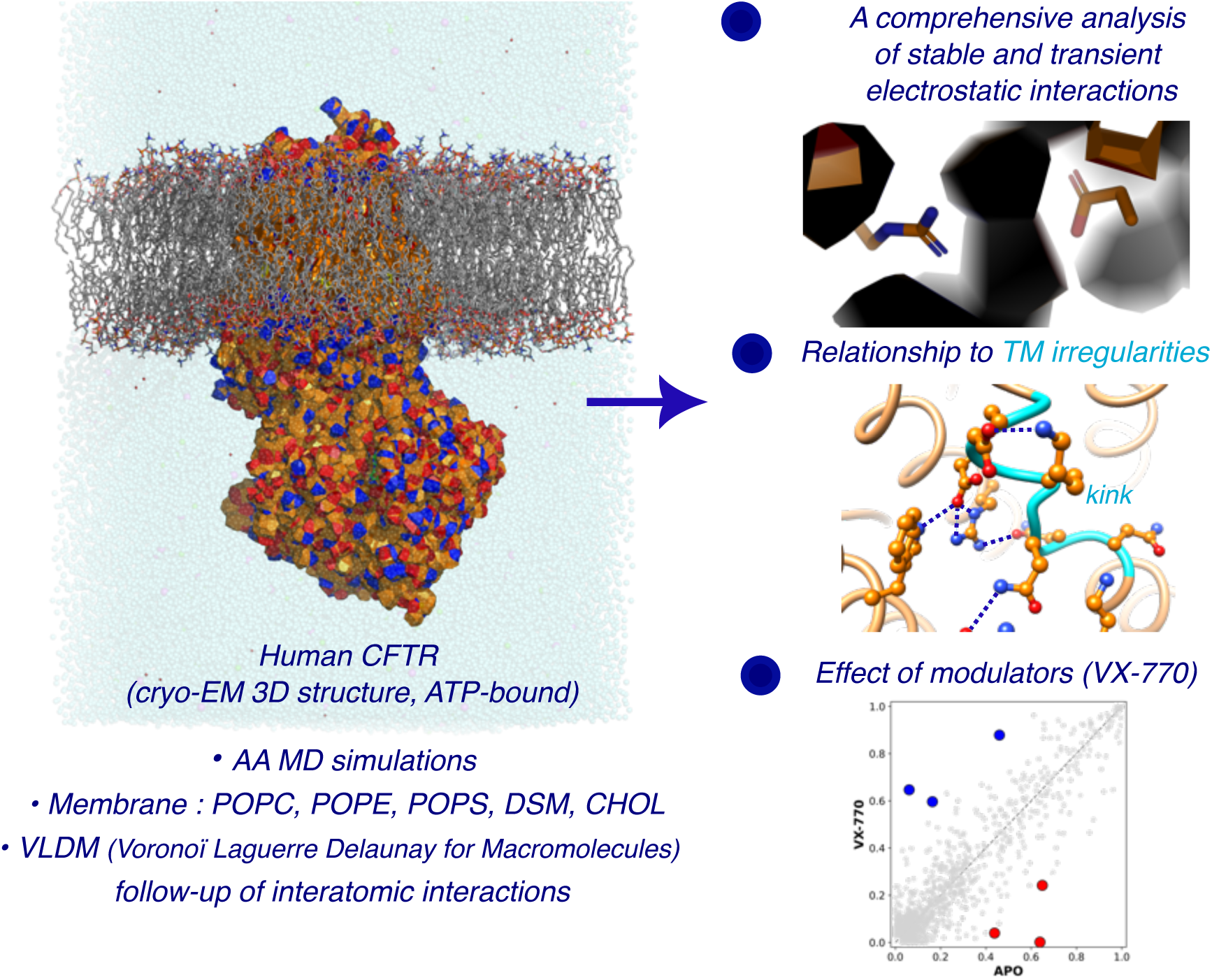

## Introduction

The cystic fibrosis transmembrane conductance regulator (CFTR) protein is a chloride/bicarbonate channel whose dysfunction is the root cause of cystic fibrosis (CF), a life-shortening genetic disease that causes inflammation and infections in the lungs and pancreatic insufficiency ^1^. As a member of the ATP-binding cassette (ABC) superfamily, CFTR has two transmembrane domains (TMDs) and two nucleotide-binding domains (NBDs) that assemble via multiple, dynamic interfaces ^2^. CFTR (ABCC7) is closely related to members of the ABCC family which, together with the ABCB family, form the type IV systems, according to the topology of the TMDs ^2–4^. The core of type IV TMDs consists of six transmembrane helices (TMs), which extend widely into the cytoplasm to form intracellular loops (ICLs) terminated by coupling helices (CHs) that interact with NBDs. In the fully assembled architecture of type IV systems, the fourth and fifth TMs of each TMD swap into the other TMD, so that the second CHs interact with the NBDs in the opposite half of the transporter.

Nine ABCC proteins (ABCC1-6 and ABCC10-12) form a group of multidrug-resistance-associated proteins (MRPs), involved in chemotherapy resistance by translocating anticancer drugs out of cells ^5^. The closest paralog of CFTR, ABCC4, transports numerous organic anions including chemotherapeutic agents, as well as various endogenous molecules, including cyclic nucleotides, folic acid, ecoisanoids, urate, prostaglandins, leukotrienes and conjugated steroids ^6^. Like other proteins of the ABCC family, CFTR possesses a N-terminal lasso motif (called linker L0 in “long” ABCC members, which possess an additional N-terminal membrane-spanning domain (TMD0)). This lasso/L0 is partly inserted in the membrane and embraces both TMDs ^7, 8^. The ABCC family also possesses long, disordered links between NBD1 and TMD2 (called the regulatory (R) domain in CFTR), the phosphorylation of which regulates function ^9^. CFTR has however evolved from this ABCC mold to exhibit regulated (ATP-gated) anion channel activity by acquiring unique structural features (reviewed in ^10^). The alternating access mechanism of ABCC transporters associated with an anionic substrate translocation pathway has thus evolved towards the construction of an anionic channel pore, in which the cytoplasmic-side gate is degraded and the open pore conformation stabilized ^11^. The specific structural features of human CFTR include an unusual conformation of the TM7-TM8 hairpin, in which unwinding (or discontinuity of the helical pattern) of a five amino acid long segment in TM8 leads to a displacement of the top of helix TM7 relative to its usual position on the extracellular side and gives rise to a pronounced asymmetry of the channel (^8^ and reviewed in ^12^). Another specific feature of CFTR is a “degenerated” intracellular gate, which is closed when the NBDs are in a dimerized state, but accompanied by an open lateral entrance (or portal) between TM4 and TM6 (TM4/TM6 portal), allowing anion flow from the cytoplasm to the inner vestibule. This characteristic is attributable to the pronounced divergence of TM6, harboring a kink at the level of arginine R352, at which TM6 can move away from TM4 to create this portal at the basis of the intracellular loops ^10^. Of note is that a symmetric, secondary TM10/TM12 pore has been identified by two independent studies using molecular dynamics (MD) simulations on a human CFTR three-dimensional (3D) structure model ^13^ and on the experimental 3D structure of zebrafish CFTR ^14^, without being demonstrated experimentally by cryo-electron microscopy (cryo-EM) studies conducted to date. Experimental data based on site-directed mutagenesis have indicated that the TM10/TM12 portal has a minor role when compared to that the TM4/TM6 portal ^15, 16^. The static cryo-EM 3D structures of the phosphorylated, ATP-bound CFTR ^17, 18^ have also failed to elucidate the mechanism of complete opening of the channel at the extracellular end. Indeed, even in complex with the potentiator VX-770 (Ivacaftor) ^17^, the first drug that improved channel activity ^19^, these structures do not have a conduction pore passing through the entire thickness of the membrane, as this one is closed near the channel selectivity filter or, in one case ^20^ , too narrow at this level to accommodate any other else than a fully dehydrated chloride ion.

Previous studies have enlightened the importance of a few electrostatic bonds (i.e. chemical bonds characterized by an electrostatic attraction) for stabilizing the open state of the CFTR channel. These involve R352_TM6_ mentioned above, forming a salt bridge with D993_TM9_, but also R347_TM6_ interacting with D924_TM8_ (at the N-terminal extremity of TM8) ^21–23^. The existence of these salt bridges was supported by cryo-EM 3D structures of human CFTR, which revealed at this depth in the lipid bilayer an additional salt bridge between R933_TM8_ (at the C-terminal extremity of the TM8 unwound segment) and E873_TM7_ ^18^. Another salt bridge was also identified within the four-helix bundle linking the four ICLs, at the level of ICL2 (E267_TM4_) and ICL4 (K1060_TM10_), which directly influence channel gating ^24, 25^. Finally, a hydrogen bond has also been highlighted between the side chain of R117 (ExtraCellular Loop ECL1) and the backbone carbonyl group of E1124_ECL4_, formed only in the open state of the channel ^26, 27^.

Our aim here was to comprehensively analyze the full networks and dynamics of electrostatic bonds (salt bridges and hydrogen bonds) between side chain atoms present in human CFTR, but also linking these side chains to ions and lipids. To that aim, we performed all-atom molecular dynamics (MD) simulations of the human CFTR 3D structure (phosphorylated, ATP-bound form), in the apo form and in the presence of the potentiator VX-770, embedded in a complex, asymmetric membrane bilayer. This analysis led to the identification of novel key partners, including some only revealed in our MD simulations (and not present in the cryo-EM 3D structure). These bonds may be critical either for the architecture of the channel or for regulating its activity, including via the potentiator VX-770, whose mechanisms of action remain incompletely understood. Finally, this study has enabled us to shed additional light on the specific evolution of CFTR with respect to other members of the ABCC family, probably linked to the optimization of the gating mechanism. Altogether, this fundamental study may provide novel insights into how the open state of the channel may be induced and stabilized and ultimately, may help the design of novel potentiators.

## Results

### Overview of the MD simulations

For this study, we performed eight independent all-atom molecular dynamics (MD) simulations of the human CFTR 3D structure (ATP-bound) embedded in a heterogeneous lipid bilayer composed of the phospholipids phosphatidylcholine (POPC), phosphatidylethanolamine (POPE) and phosphatidylserine (POPS), together with cholesterol (CHL1) and the sphingolipid DSM, using proportions similar to those proposed by Domicevica and colleagues ^28^ to study the behavior of the P-glycoprotein in a human brain epithelial cell model (see Methods). This composition is consistent with the mean composition of epithelial cells from a variety of biological sources (as reviewed in ^29^). This system has the advantage of offering a membrane that is closer to physiological ones, with an asymmetric distribution of lipids between leaflets (POPC and DSM mostly appearing in the outer leaflet and POPE and POPS being the dominant species of the inner leaflet), whereas molecular dynamics simulation studies carried out to date have only considered simplified models (POPC only, ^14, 30–35^). We kept the stabilizing mutation E1371Q (NBD2 Walker B motif) preventing ATP hydrolysis in the canonical ATP binding site, as we wish to specifically analyze the network related to channel opening. Such a condition was also adopted in another computational analysis of the structure and properties of the CFTR open state ^31, 32^. Four simulations were made in the absence of ligand (apo conditions, labeled Apo1, Apo2, Apo3, Apo4) and four in the presence of the potentiator VX-770 (Ivacaftor) (ligand-bound conditions, VX1, VX2, VX3 and VX4). Each simulation was run for either 500 ns or 1000ns, yielding a total of 6 μs of simulation time (**Table S1**). To further ensure the robustness of our results, we employed different thermostat–barostat combinations across the simulations (Nosé–Hoover / Parrinello–Rahman, V-rescale / C-rescale; **Table S1**, see Methods). This strategy allowed us to confirm that the observed behaviors were independent of the coupling scheme and thus not an artifact of a specific algorithm.

To evaluate the structural stability of the protein, we calculated the root-mean-square deviation (RMSD) of CFTR backbone atoms relative to the starting cryo-EM structure. As shown in **Figure S1**, all trajectories reached convergence within the first 100–200 ns, with RMSD values stabilizing between 2.5 and 3.5 Å, indicating stable overall protein conformations in both apo and ligand-bound conditions. In parallel, we examined membrane behavior to ensure bilayer stability across all simulations. The area per lipid (APL) and membrane thickness were computed for each replica, as shown in **Figure S2**. These metrics remained stable throughout the simulations, with an area per lipid of approximately 62 Å² and an average membrane thickness of 45 Å, confirming that all systems remained equilibrated and structurally consistent. Compared to a pure POPC bilayer (thickness ∼37 Å and with APL ∼65 Å² in ^34^), the mixed bilayer appeared thicker. This observation is consistent with findings from other ABC transporters, such as P-glycoprotein, where POPE-containing systems produce more ordered bilayers (smaller APL and greater thickness ^36^) and cholesterol-rich bilayers induce increased membrane thickness ^37, 38^.

These MD simulations enabled reliable analysis of residue-level electrostatic interactions, as well as contacts with anion and lipid headgroups in both the apo and ligand (VX-770)-bound conditions. These contacts were calculated using a rigorous, geometry-based definition of interatomic interactions, based on a tessellation method used by the VLDM (Voronoi Laguerre Delaunay for Macromolecules) program (see Methods). This geometrical approach, integrating all heavy atoms of the MD simulations including water, has the advantage of allowing for a comprehensive analysis of the entire system, without preconceptions and without imposing arbitrary distance cutoffs. Contacts are correlated to the presence of electrostatic bonds (salt bridges (SBs) and hydrogen bonds (HBs)), terms that will be used appropriately in cases supported by 3D structure inspection. From the analysis of the MD trajectories, we identified 798 electrostatic interactions between charged or polar side-chain atoms within CFTR’s 3D structure. We included in our analysis basic (R, K, H) and acidic residues (D, E), as well as S, T, N, Q, C, Y and W. Interactions of W with R (cation-π and π-π interactions) and with H were not considered in the statistics of electrostatic interactions, but illustrated when associated with networks of electrostatic interactions. To focus on robust and functionally relevant interactions, we considered only bonds that formed for at least 10% of the total simulation time across all apo simulation replicates, resulting in a final set of 557 electrostatic interactions. The frequencies of these interactions are reported for each of the eight MD simulations in **Figures S3 to S4** (amino acids pairs, presented with the first amino acids of the pairs following the order of the polypeptide chain progression), while frequencies of contacts of polar/charged amino acids with ions and lipids are listed in **Figures S5 to S6**. We also recorded electrostatic interactions linking charged or polar side-chain atoms and main-chain atoms (738 contacts with frequencies above 0.1 across all apo MD simulations). In the following chapters, we analyzed in detail the side-chain/side-chain networks by focusing on the most robust ones (frequency ≥ 0.25; 337 interactions), while only illustrating some notable features of the side-chain/main-chain networks.

### Spatial distribution of CFTR electrostatic interactions and functional impact

Electrostatic interactions between side chain atoms were categorized by their location into five structural regions of CFTR, based on the average Z-coordinate of the involved residues (see Methods): (i) the extracellular loops (ECLs), (ii) the transmembrane (TM) helices (TM helices/Membrane), (iii) the membrane-proximal elbow and lasso regions along with the intracellular loops (Elbow/Lasso/ICLs), (iv) the ICLs-nucleotide-binding domains (ICLs:NBDs) interface and (v) the NBDs. **Figure S7** illustrates the distribution of these contacts in each region, ordered by mean frequency, while **Figure 1** maps their spatial distribution on CFTR 3D structure where each residue is color-coded by its assigned structural region (red for ECLs, orange for TM helices/Membrane, yellow for elbow/lasso/ICL regions, green for ICLs:NBDs interface and blue for NBDs). We then classified electrostatic interactions by whether the basic amino acids (K,R,H) or S/T/N/Q/Y/W/C simultaneously engaged in anion coordination (cyan), lipid binding (yellow), both (purple), or neither (light gray) (**Figure S8**). Strikingly, over half of the frequent contacts in the ECLs, TM helices/Membrane, Elbow/Lasso/ICLs regions have dual functionality: many of the K/R/H/S/T/N/Q/Y/W/C residues forming electrostatic interactions with other residues also served as binding sites for chloride or bicarbonate ions, membrane lipids, or both. This suggests that several non-covalent bonds in CFTR are components of larger electrostatic networks that link the protein’s intramolecular contacts to its interactions with the membrane environment and anions.

**Figure 1:**
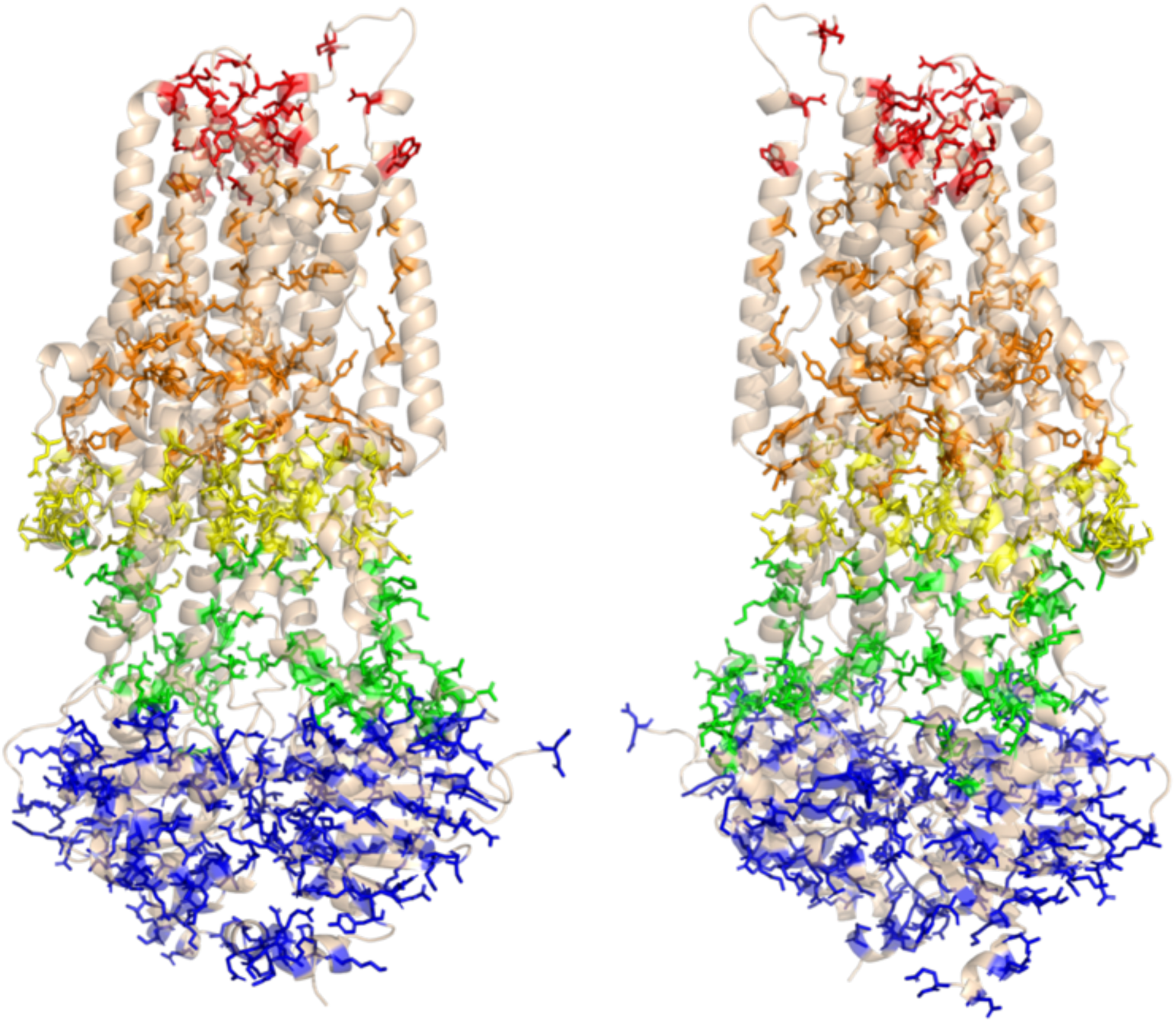
3D mapping of CFTR residues involved in side-chain/side-chain electrostatic interactions. Two views of the human CFTR 3D cryo-EM structure (pdb 6O2P, 180° apart along the z-axis, ribbon representation), show the spatial distribution of residues engaged in electrostatic interactions (side-chain/side-chain contacts, mean frequencies ≥ 0.25). Residues are represented as sticks and color-coded according to their location on the 3D structure: red for ECLs, orange for transmembrane helices / membrane regions, yellow for elbow/lasso/top ICLs regions, green for the bottom ICLs:NBDs interface and blue for the NBDs. A coordinate-based scheme along the Z-axis was used to classify residues into these five structural regions (see Methods).

We then analyzed in detail these networks, region by region, starting from the extracellular side toward the cytoplasm.

#### ECLs

ECLs can be divided into two distinct groups : (i) ECL1, ECL3, ECL4 and ECL6 (colored in blue, green, yellow and red, respectively, in **Figure 2A**), which form the core of the transmembrane assembly, (ii) the remaining two shorter ECLs (ECL2 and ECL5, grey in **Figure 2A**), which link the four three-helix units (TM1-2-3, TM4-5-6, TM7-8-9 and TM10-11-12) ^39^ and are located at the periphery of the transmembrane assembly. The vast majority of the intra-and inter-ECL interactions (side-chain/side-chain and side-chain/main-chain, **Figure S7**) are observed at the level of the core ECL1, ECL3, ECL4 and ECL6 and are transient in nature (mean frequencies below 0.6, details of the amino acids involved with frequencies ≥ 0.25 in **Figure 2A**).

**Figure 2:**
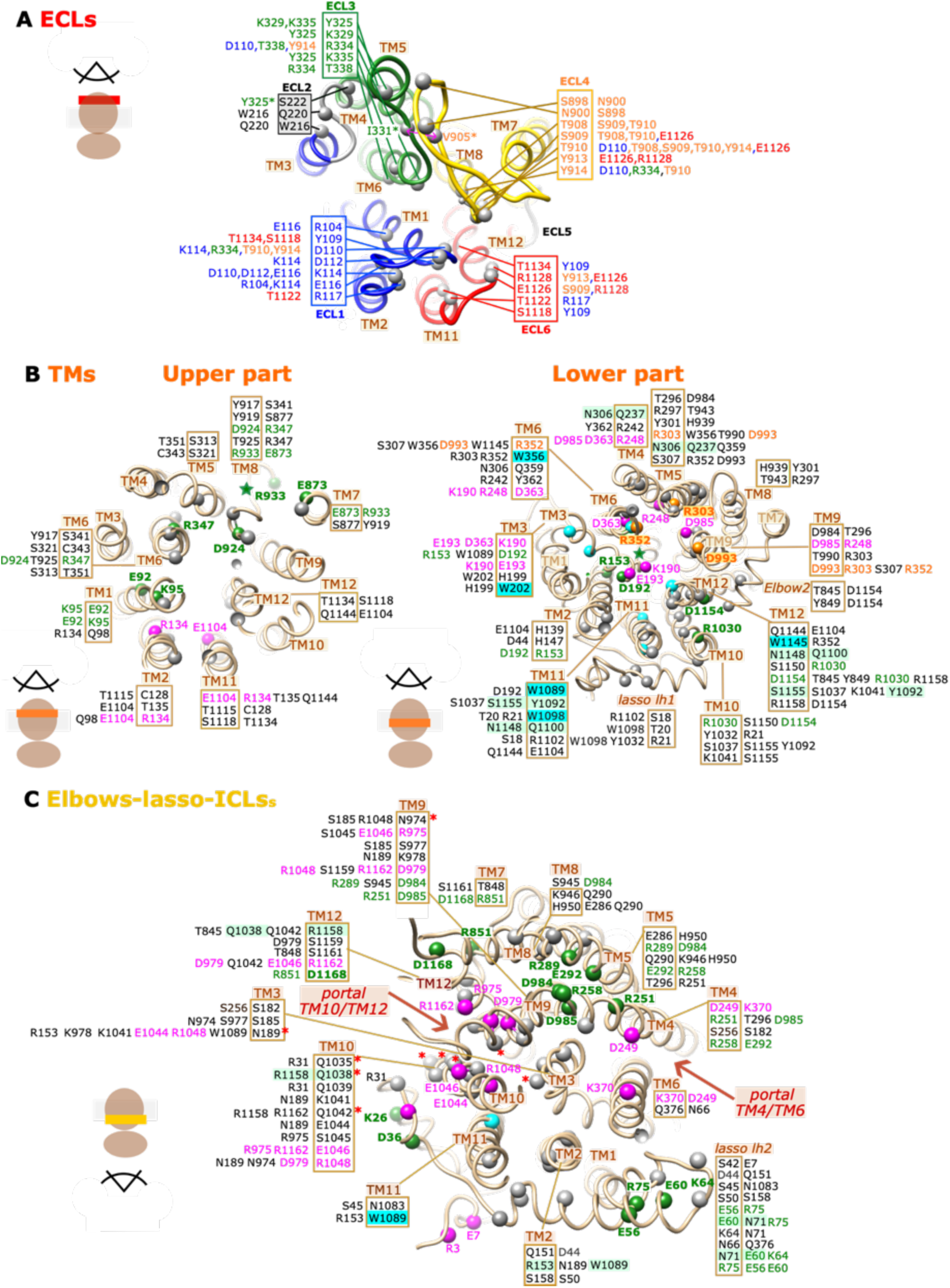
Electrostatic interactions in the transmembrane domains (TMDs) of CFTR. Amino acids involved in these interactions are visualized on the human CFTR 3D structure after 500 ns of MD simulation (apo condition, replica 1). **A) Within the ECLs** (region colored in red in Figure 1, view from the extracellular side). The charged/polar amino acids involved in electrostatic interactions (side-chain/side chain interactions, with frequencies ≥ 0.25), as listed in Figure S7, are depicted with balls (Cα atoms only) and colored according their belonging to ECL1 (blue), ECL3 (green), ECL4 (yellow) and ECL6 (red). Next to amino acids in the ECL boxes are their intra-and inter-ECL partners. ECL4 is particularly long and is not fully resolved in current cryo-EM structures; therefore, it was modeled in our simulations to ensure the structural continuity and completeness required for the accurate representation of CFTR dynamics in this extracellular region. Only one, highly stable HB is present between the S222^ECL2^ side chain and the Y325^ECL3^ main chain (asterisk) N atom. Main chain/main chain (asterisks) HBs were also observed between atoms of ECL3 (I331) and ECL4 (V905), leading to the establishment of a short, inter-ECL β-sheet. In addition to being involved in side-chain/side-chain interactions, R117 side chain N atoms were also observed to also establish contacts with main chain oxygen atoms of other amino acids from ECL1 (<u>Y109</u>^TM1^, <u>D110</u>^ECL1^, P111^ECL1^) and ECL6 (<u>I1119</u>^TM11^, L1120^TM11^, T1121^ECL6^, <u>T1122</u>^ECL6^, <u>G1123</u>^ECL6^, <u>E1124</u>^ECL6^, <u>G1125</u>^ECL6^, G1127^ECL6^) (frequencies above 0.25 are underlined). **B) Within the membrane regions of the transmembrane domains** (region colored in orange in Figure 1, view from the extracellular side), cut into two approximate layers (upper and lower parts). As in panel A, the charged/polar amino acids involved in electrostatic interactions (side-chain/side chain interactions, with frequencies ≥ 0.25), as listed in Figure S7, are depicted with balls (Cα atoms only). Highly stable salt bridges are shown in green, while less stable ones are in pink. The particular case of bonds linking D993^TM9^ to R303^TM5^ and R352^TM6^ is highlighted in orange. Tryptophan residues, which are mostly involved in cation-π interactions with arginine residues, are shown in cyan. Amino acids participating in stable HBs at the level of the swapped TM4-TM5 (carrying ICL2 CH) and TM10-TM11 (ICL4 CH) hairpins are shaded green. Next to amino acids included in TM boxes are their partners. **C) At the level of the Elbow-lasso-ICLs** (region colored in yellow in Figure 1, view from the intracellular side). The same colors as in panel B are used to highlight stable and transient salt bridges. Next to amino acids included in TM boxes are their partners.

In particular, these interactions form an intricate network at the center of the ECL assembly, reflecting the dynamic and solvent-exposed nature of the extracellular region. Participating to this network are D110_ECL1_ and E116_ECL1_, which have been shown in an earlier study to stabilize the open pore architecture ^26^ (details illustrated in **Figure 2A**). In particular, D110_ECL1_ interacts with amino acids from ECL3 and ECL4, among which R334_TM6_ and Y914_TM8_, which both play a key role in controlling anion flux ^26, 40^. Only one stable interaction (mean frequency above 0.6) is observed between the peripheral ECL2 and ECL3 (side chain/main chain interaction, **Figure S7** and **Figure 2A** *(main chain designated with an asterisk)*) and, in connection with this one, we noted the presence of an HB network between ECL3 and ECL4 main chain atoms, leading to a short *β*-sheet, which likely contributes to stabilize the ECL assembly at this level (**Figure 2A)**. No such features were detected for the symmetrical ECL5/ECL6/ECL1, in which ECL5 remains disconnected from ECL6. With regards to ECL1, we retrieved in our side-chain/main chain analysis the previously described HB made by R117_ECL1_ (an amino acid also known to stabilize the open pore ^26^) with the backbone oxygen atom of E1124_ECL6_ ^27^, but also observed high variability in the position of its side chain, being able to establish HBs with main chain atoms of several other amino acids from ECL1 and ECL6 (**Figure S7** and legend of **Figure 2A**). This dense network likely contributes to the stabilization of the ECL1 conformation.

Overall, the analysis of electrostatic interactions in the ECL region reveals an intricate and dynamic network in which several charged and polar amino acids, including D110_ECL1_, R117_ECL1_, R334_TM6_, Y914_TM8_, E1126_ECL6_ and R1128_ECL6_, play a central role (**Figure 2A**).

#### The transmembrane (TM helices/Membrane) region

Contrasting with ECLs, the membrane region contains five SBs which are highly stable (mean frequency > 0.8) across simulations (**Figure S7**; amino acids highlighted in green in **Figure 2B**). The high stability of these SBs suggests a structural role in maintaining the integrity of the CFTR transmembrane architecture. Three of them (R933_TM8_–E873_TM7_, R153_TM2_–D192_TM3_ and R347_TM6_–D924_TM8_) are located in the vicinity of TM irregularities, involving TM8 (unwound segment) and TM3 (green stars in in **Figure 2B**), suggesting that they may be critical for maintaining such particular conformations (further discussed below). Of note is that the R347_TM6_ and R933_TM8_ side-chains are also involved in multiple HBs with main chain atoms (**Figure S7)**, contributing to reinforce interactions in the unwound TM8 region. R347_TM6_–D924_TM8_, has been described in earlier studies as playing a key role in the channel function, as it is also the case of R352_TM6_-D993_TM9_ ^21, 22, 41^. However, this last SB, observed in the cryo-EM 3D structure ^18^, appears to be far less stable (mean frequency < 0.3), while D993_TM9_ is more frequently salt bridged with R303_TM5_ (mean frequency ∼0.7) (orange in **Figure 2B**). Several basic residues which are involved in transient SBs, such as R134_TM2,_ K190_TM3,_ R303_TM5_, R248_TM4_ and R352_TM6_, are positioned along the channel pore and were found to engage in anion contacts, supporting a potential relationship between reduced salt bridge stability and functional involvement in anion interactions through electrostatic interactions. In contrast, the stable SB R1030_TM10_-D1154_TM12_ (green in **Figure 2B**) is located at the periphery of the membrane assembly, more precisely at the top of the potential TM10/TM12 portal, and appears accessible to lipid headgroups. Notably, only one SB in this region, R1158_TM12_–D1154_TM12_, was found to have both anion and lipid contacts (**Figure S8**), although with very low frequency. Its strategic position in the upper part of the TM10/TM12 portal may reflect a dual role in membrane coupling and ion conduction. Beside SBs, stable HBs were found at the level of the swapped TM4-TM5 (carrying ICL2 CH) and TM10-TM11 (ICL4 CH) hairpins, suggesting key roles in their proper positioning and/or packing (shaded green in **Figure 2B**). Several tryptophan (W) residues are found in this lower part of the membrane region (cyan in **Figure 2B**), principally making cation-π interactions with arginine (R) residues involved in SBs (stars in **Figure S7**), thereby strengthening the bond networks. Also of note at this level, the stable contact between H199_TM3_ and W202_TM3_ (**Figure 2B**), both amino acids belonging to an hydrophobic pocket at the TMD1 surface, which interacts with the correctors VX-809 (Lumacaftor) and VX-661 (Tezacafor) ^42, 43^.

In conclusion, the membrane region, and especially its lower part, concentrates a large number of electrostatic interactions (SBs and HBs), forming dense networks and involving amino acids from all the TMs. Both stable interactions and more fluctuating ones are observed, the first ones contributing to the architecture of the pore and stabilization of its open conformation, while the second ones appear to play active roles in providing basic residues to contact ions.

#### The Elbow/Lasso/ICLs region

A combination of stable and transient interactions is also observed in this region, including the inner vestibule and lateral portals. Ten SBs exhibited high stability (mean frequency > 0.8) (**Figure S7** and **Figure 2C** (amino acids depicted in green)) and thus appear to play a structural role on both sides of the lateral portals. Some of them, within the lasso region and at the base of TMD2, are also involved in contacts with lipids (**Figure S8** and **Figure 2C**), suggesting a critical role in membrane insertion/anchoring, or with ions (**Figure S8** and **Figure 2C**), in continuity of the anion conduction network described before. Several less frequent electrostatic interactions (**Figure S7)** form a tightly interconnected network at the level of the inner vestibule and of the portals providing access to it (**Figure 2C**). Central to this network are several asparagine (N) and glutamine (Q) residues (**Figure S7** and red asterisks in **Figure 2C)**, whose amide groups may act simultaneously as hydrogen bond donors and acceptors. The network in the vicinity of the TM4/TM6 portal (a known access route for anions) integrates few amino acids, except for the transient K370_TM6_–D249_TM4_ SB, which do not exhibit contacts with lipids or anions (**Figure 2C** and **Figure S9A**). The relatively low stability of this particular SB raises the question of whether it plays a transient or conditional role in stabilizing the portal architecture, potentially responsive to conformational shifts or binding events. In contrast, the region of the TM10/TM12 portal (which has been described only at the computational level ^13, 14^, while site-directed mutagenesis indicated it plays a minor role as compared to the TM/TM6 portal ^15, 16^) is teeming with interactions, including the most stable HB linking the Q1038_TM10_ and R1158_TM12_ side chains (**Figure 2C** and **Figure S9A**). The residues involved form an extended network spanning TM9, TM10, TM12, and the ICLs, converging around a region leading to the intracellular vestibule and involved in anion coordination ^44, 45^. The recurrence of overlapping residues across multiple interactions within this cluster suggest a cooperative mechanism, where dynamic bond switching may modulate local electrostatics and facilitate anion conduction or structural flexibility. As for membrane regions, several HBs are observed linking side-chain to main-chain atoms in the vicinity of side-chain/side-chain bonds (**Figure S7**), therefore contributing to reinforce the observed networks.

#### The ICLs:NBDs interface

Here, SBs and HBs linking side chains appear to play primarily a structural role, as none was observed to participate in anion or lipid headgroup interactions (**Figure 3A, Figures S7** and **S8**). This suggests that their main role is to stabilize the coupling between the TMDs and the NBDs, in particular through amino acids belonging to the ICLs CHs. The most stable bonds (green in **Figures 3A** and **S9B**) are also here located at the periphery of the CFTR domain assembly and likely act as buttresses, helping to maintain the proper alignment and communication between ICLs and NBDs. The importance of R1066_CH4_ in this network is supported by the fact that mutations of this amino acid in patients with CF (R1066H and R1066C) are associated with a severe class II defect, which is rescued by ETI (Elexacaftor (VX-445), Tezacaftor (VX-661), Ivacaftor (VX-770)) therapy ^46^. Furthermore, always at the ICL4 level, there are multiple HBs between arginine side chain atoms (R1066_CH4_, R1070_TM11_) and main chain atoms, including a very stable one between R1070_TM11_ and F508_NBD1_ (**Figure S7**). This tight network of electrostatic interactions, including amino acids from ICL1, ICL4 and NBD1 but also from the TMD1-NBD1 linker, is located at the level of the non-canonical/degenerate ATP-binding site (accommodating ATP(1) in **Figure 3A**), at which prolonged ATP retention is observed ^47–49^. A tight network also exists at the ICL2:ICL3:NBD2 interface, including an amino acid of the TMD2-NBD2 linker and integrating several HBs established between side and main chain atoms (**Figure S7**).

**Figure 3:**
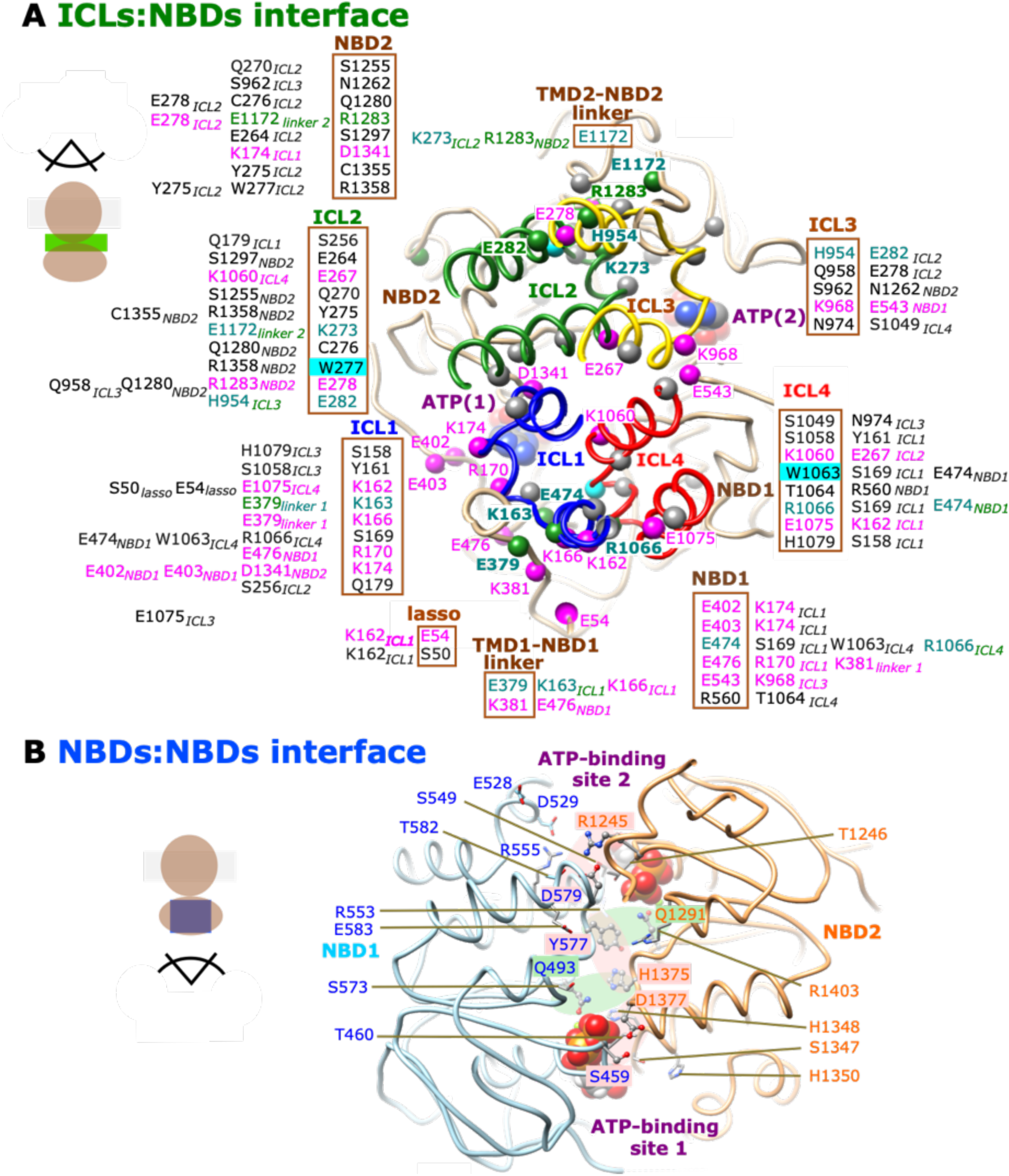
Electrostatic interactions at the ICLs:NBDs interface (A) and at the NBD1:NBD2 interface (B). These interactions, visualized on the human CFTR 3D structure after 500 ns of MD simulation (apo condition, replica 1), are extracted from regions colored in green (A, viewed from the extracellular side) and blue (B, view from the intracellular side) in Figure 1. **A)** The charged/polar amino acids involved in electrostatic interactions (side-chain/side chain interactions, with frequencies ≥ 0.25), as listed in **Figure S7**, are depicted with balls (Cα atoms only). Highly stable interactions between charged amino acids are shown in green and less stable contacts are in pink (labels shown on the 3D structure only for frequencies ≥ 0.5). Tryptophan (W) residues are shown in cyan. **B)** Electrostatic interactions linking NBD1 (blue) and NBD2 (orange). ATP (spheres) is bound at both ATP-binding sites (non-hydrolytic ATP-binding site 1 - bottom and hydrolytic ATP-binding site 2 - top).

Several additional electrostatic interactions exhibited intermediate stability (mean frequency between 0.5 and 0.8, labels in pink on the 3D structure in **Figure 3A**). These moderately stable interactions likely contribute to dynamic structural adjustments that support the conformational transitions required for CFTR function while maintaining sufficient anchoring of intracellular elements. Of note is the central K1060_TM10_–E267_TM4_ SB, at the very C-terminal ends of TM10 and TM4, which helps to maintain the ICL four-helix bundle assembly and was previously shown to be critical for the CFTR function ^24, 25^.

#### The NBDs

Numerous bonds were identified here, with a noticeably higher concentration within NBD1 compared to NBD2 (**Figure S7**). Interestingly, the majority of these bonds exhibited low stability, with approximately two-thirds showing mean frequencies below 0.5. This may reflect the inherent flexibility of the NBDs and their dynamic behavior during the ATP-binding and hydrolysis cycle. The lower stability of these bonds could also suggest a more transient regulatory or allosteric role rather than a purely structural function. This regional asymmetry in electrostatic interactions distribution and stability may be linked to the non-equivalence of NBD1 and NBD2 in CFTR’s transport mechanism ^50, 51^. NBD1 plays a critical role in signal transmission leading to CFTR gating through its interface with the TMDs, involving the coupling helices from ICL1 and ICL4 (*e.g.* ^52^). This suggests that electrostatic interactions within NBD1 may come in support to the allosteric pathways linking the ATP-binding sites ^53^ and phosphorylated regulatory insertion (RI) ^54^ to the coupling interfaces. On another hand, it is possible that electrostatic interactions can help to counteract NBD1 intrinsic instability, in particular conferred by the flexible regulatory insertion ^55, 56^. Finally, they also can help some steps of NBD1 co-translational folding, which is particularly complex, involving the sequential and coordinated compaction of its three sub-domains, and which is finely tuned by native codon usage allowing to limit aggregation ^57–59^.

Several bonds bridging NBD1 and NBD2 exhibited moderate stability (mean frequency > 0.5) (**Figure 3B** and **Figure S7**). The two first ones (S459_NBD1_–D1377_NBD2_ and R1245_NBD2_–D579_NBD1_) involve two homologous pairs of amino acids: on the one hand, the two aspartic acids of the D-loop and, on the other hand, two amino acids preceding the Walker A motif. These bonds lock both ends of the dimer interface (**Figure 3B**). Three other bonds involve two homologous amino acids (Y577_NBD1_ and H1375_NBD2_) located two residues upstream of the D-loop aspartic acids and which interact either together or with the two Q-loop glutamine residues of the opposite NBD (Q1291_NBD2_ and Q493_NBD1_) (**Figure 3B**). In ABC transporters, the position of these two homologous amino acids located in the center of the dimer interface are generally occupied by small amino acids (G, A, S), except for CFTR (Y577_NBD1_ and H1375_NBD2_) and for members of the ABCE (Y) and ABCF (H, N, M) families ^49^. A dozen other bonds also contribute to these interdomain interactions, distributed all along the NBD dimer interface (**Figure 3B**), among which the one comprising H1348_NBD2_ (the first histidine of the degenerate (LS<u>H</u>GH) ABC signature) forming an alternative bond with Q493_NBD1_ to that established with H1375_NBD2_. In conclusion, all these observed bonds distributed throughout the dimer interface may contribute to maintain its alignment or to facilitate proper conformational transitions during ATP hydrolysis. Notably, their moderate stability could reflect a balance between the need for dynamic dissociation (during NBD opening) and transient stabilization (during dimer formation), underscoring a potential mechanistic role in the gating cycle of CFTR. We emphasize here the partners of the D-loop motifs, which have been shown in several ABC transporters to play a central role in mediating allosteric communication between the two ATP-binding sites, thereby regulating NBD dimer dynamics and ATP hydrolysis. It is suggested to play asymmetric roles, with the consensus site D-loop being essential for ATP hydrolysis, while the degenerate site D-loop stabilizes the NBD conformation ^49, 60^. Strikingly, most of these inter-NBDs bonds were only captured by MD and were not formed in the cryo-EM structures of CFTR (**Figure S9C**), suggesting their transient or state-dependent nature and pointing to a subtle but possibly critical mechanism for aligning the NBD dimer during the gating cycle.

### Mapping of selective ion contact sites across CFTR – insights into channel openings

We then analyzed contacts of charged or polar side-chain atoms (R, K, H, S, T, N, Q, Y, W) with anions (chloride and bicarbonate) in each MD simulation replica in the apo conditions (**Figure 4 and Figures S10-S11**). The three most frequent contact sites with both chloride and bicarbonate ions are R303_TM5_, R248_TM4_ and K190_TM3_ (orange in **Figure 4A, Figure S10**). The initial attraction typically occurs at R248_TM4_, which likely serves as the primary entry point for anions through the portal displayed between TM4 and TM6 (TM4/TM6 portal), which is stably open in our MD simulations (**Figures 4B,4D**) and emerges as the most frequent and accessible anion entry pathway in our simulations. The ion may be then coordinated in the inner vestibule by R251_TM4_ and/or K190_TM3_, afterwards transferred to R303_TM5_ and R352_TM6_, positioned higher along the channel axis, and ultimately to K95_TM1_ (green in **Figure 4A**). Several other amino acids along this pathway are also involved in specific interactions with chloride ions (**Figure 4A**). Residues identified here at TM4/TM6 portal and in the central conduction pore are consistent with the results obtained from mutagenesis experiments ^15, 16, 45^. However, it should be noted that K370_TM4_, at the entrance of the TM4/TM6 portal, was not detected here in the group of anion binding residues, while it is involved in a transient SB with D249_TM4_ (**Figure 2C**). This suggests that this interaction plays an essential role in regulating the anion flux. Interestingly, the formation of this D249_TM4_-K370_TM6_ SB divides the TM4/TM6 portal into two distinct, smaller portals (**Figure 4D**), suggesting that breakage of the bond could be associated with a higher ion flux.

**Figure 4:**
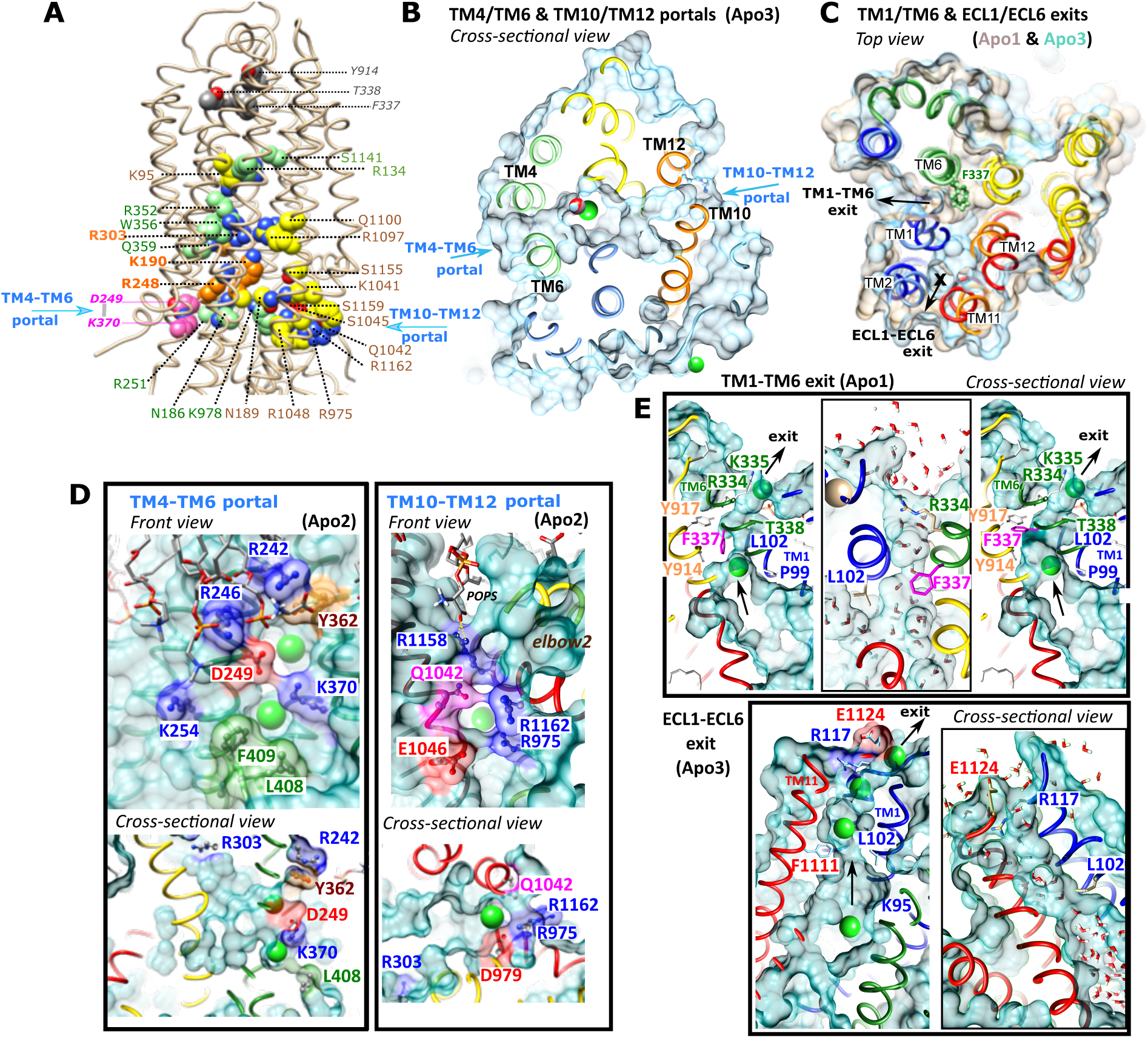
Electrostatic interactions with ions – intracellular portals and extracellular exits. **A) Electrostatic interactions with ions** are visualized on the human CFTR 3D structure after 500 ns of MD simulation (apo condition, replica 1). The carbon atoms of basic and polar side chains involved in electrostatic interactions with ions (chloride and bicarbonate) are represented in orange (most frequent ones) and green, whereas those of residues involved in specific interactions with chloride ions are shown in yellow. R370^TM6^ and D249^TM4^, which form a transient salt bridge at the entrance of the portal TM4/TM6, are shown in magenta. Amino acids of the dehydrated ion binding site (selectivity filter) identified by ^16^ in the upper part of the channel (T338 ^TM6^, F337^TM6^ and Y914^TM8^) are colored in gray. For clarity purpose, the lasso is not represented in this view. **B-E) Views of the CFTR portals and exits. (B-C)** General views of the portals **(B)** and exits **(C)**, shown on different replica of MD simulations (Apo states, superimposed last frames). The ECL1-ECL6 exit is only open in the Apo3 MD simulation, while closed in the Apo1 one (closure symbolized with a cross). **(D-E)** The solvent-accessible surfaces (last frame of MD simulations) are represented, illustrating the topologies of the portals (**D**, front and cross-sectional views) and the extracellular exits (**E**, cross-sectional views). Chloride ions (green) were positioned along the pore to visualize the conduction paths. For the extracellular exits (**E**), the boxed views show the water molecules within the conduction paths. For the exit TM1/TM6 (Apo1), the right-hand view illustrates the widening of the path at the selectivity filter following an alternative F337 rotamer.

Amino acids that specifically bind chloride ions (yellow on **Figure 4A**) are mostly found on the opposite side of the TM4/TM6 portal, at the level of the TM10/TM12 portal. The most frequent interaction occurs with K1041_TM10_ and R1048_TM10_, with involvement of several other residues at the entrance of the portal and deeper towards to intracellular vestibule (**Figure 4A**). Opening of this portal is repeatedly observed during our MD simulations (**Figures 4B,4D**). Remarkably, all these conduction-related residues were also involved in SB/HB.

The crucial question that currently remains related to the CFTR anion channel is that of its opening on its extracellular side (extracellular exits), a mechanism that has not yet been revealed by any 3D structure obtained by cryo-EM. An in-depth analysis of MD simulations in the absence of VX-770 shows two exit pathways occurring intermittently (**Figures 4C, 4E and Table S2**). The first extracellular exit, surrounded by TM1 and TM6 (TM1/TM6 exit), in the passage of which F337_TM6_ is found, is either not present, small or larger and conditioned by a F337_TM6_ rotamer. A similar route was already highlighted by Farkas et al, using metadynamics on the zebrafish CFTR 3D structure ^14^, as well as by a more recent study having conducted massively repeated simulations starting from chloride permeable conformations ^32^. The second exit observed here is displayed between ECL1 and ECL6 (ECL1/ECL6 exit, thus at the center of the 4-helix bundle formed by TM1-TM12 (inner side) and TM2-TM11 (outer side)), which is either closed, of reduced size and conditional (Apo4), or even large in the case of the Apo3 MD simulation. This exit is similar to the one called TM1-TM12 presented in the study mentioned above ^32^. In our Apo3 replica, after 1 µs of MD simulation, the interaction of R117_ECL1_ with a lipid led to closure of this conditional exit in its terminal part, while this closure can easily be lifted when R117_ECL1_ interacts with E1124_ECL6_. Consideration of ECL4 in its entirety (while this one is incomplete in the cryo-EM 3D structures) appears critical for the conformation of the whole set of ECLs and for the opening mechanism, in particular involving the interaction of R899_ECL4_ with lipids (see below). Of note is that R899_ECL4_ did not show any interaction with anions, although this residue is responsible for the CFTR activity as an extracellular chloride sensor ^61^. In this context, it would be interesting to explore further whether the transmission of the gating information by R899_ECL4_ could occur through an effect on its interaction with lipids.

### Mapping of selective lipid contact sites across CFTR

We next examined the lipid selectivity of charged or polar side-chain atoms (K, R, H, S, T, N, Q, Y, W) by specifically analyzing their interactions with cholesterol and phosphatidylserine (POPS), as these two lipids have been shown to have a direct influence on the intrinsic CFTR activity ^62–64^. Distinct spatial patterns of lipid binding were observed across the protein (**Figure 5 and Figure S11**). Cholesterol contacts were concentrated at specific sites (red/orange in **Figure 5**), especially in the vicinity (H939_TM8_, Y247_TM4_, R242_TM4_, Y362_TM6,_ R297_TM5_ and R298_TM5_ and within (Y304_TM5_) the VX-770 binding site ^17, 65^. Contacts with cholesterol were also observed for several amino acids within ECL4 and ECL6 (red in **Figure 5**-left**, Figure S11**). Additional frequent or occasional contacts were detected within (R21_lh1_) or in the vicinity (R1102_TM11_) of the lasso domain (orange in **Figure 5, Figure S11**). Interestingly, R1102_TM11_, is involved in the VX-445 (Elexacaftor)-binding site ^66^. Altogether, these observations suggest that cholesterol may contribute to the stabilization of the membrane insertion of the elbow and lasso domains, and to the regulation of the cytoplasmic-facing transmembrane helices involved in CFTR gating and/or allosteric regulation.

**Figure 5:**
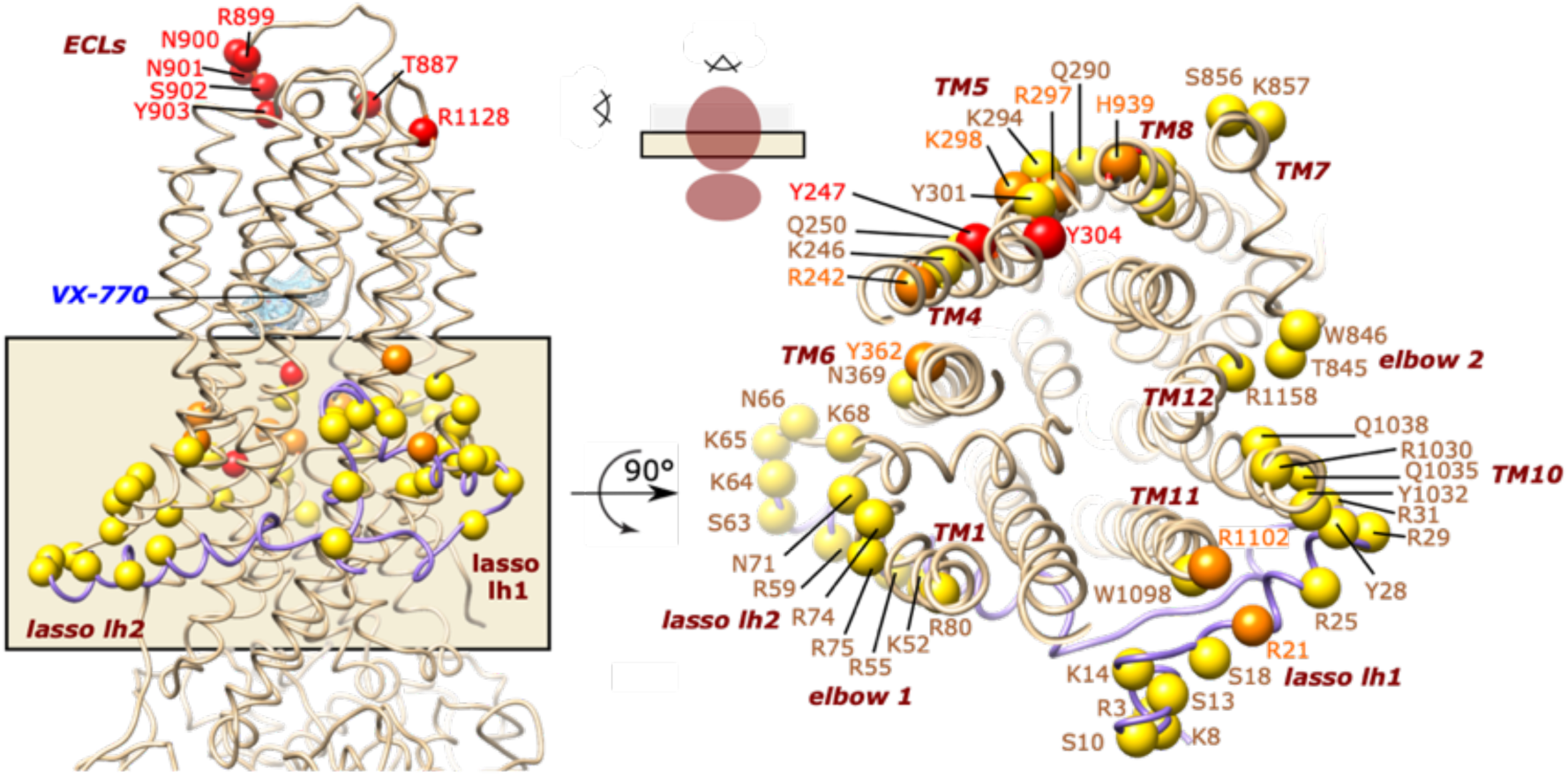
Electrostatic interactions with lipids. These interactions are visualized on the human CFTR 3D structure after 500 ns of MD simulation (apo condition, replica 1 - Two orthogonal views (at different scales) are provided, with the cross section seen from the transmembrane region). Cα atoms of basic and polar side chains involved in electrostatic interactions with POPS are represented in yellow, whereas specific interactions with cholesterol are shown in orange and red (depending on whether they overlap or not POPS binding sites, respectively).

In contrast to cholesterol, POPS-binding residues (yellow and orange in **Figure 5, Figure S11**) were more widely distributed and exhibited generally higher contact frequencies. Interestingly, several of the most significant POPS-binding regions overlapped with cholesterol-binding sites (orange in **Figure 5, Figure S11**). Prominent contacts were found within the lasso and proximal R1102_TM11_, as well as in TM5, TM4 and TM8, at the level of the headgroup region of the inner membrane leaflet, proximal to the VX-770 binding site. The elbow_1_ region and the elbow_1_:lasso interface displayed strong contacts with POPS (yellow in **Figure 5, Figure S11**). The elbow_2_ region and proximal TM10 and TM12 amino acids also contacted POPS (yellow in **Figure 5, Figure S11**), despite not being involved in cholesterol interactions, pointing to distinct lipid specificity in this region. Overall, the distribution of POPS interactions appeared asymmetric, concentrated mainly on the NBD1-facing side of the CFTR structure, driven by substantial involvement of the lasso and elbow1 regions. This asymmetric lipid binding pattern may play a role in anchoring or orienting CFTR in the membrane and modulating region-specific conformational dynamics.

In conclusion, lipid interactions across CFTR were not random but instead revealed organized patches of lipid selectivity, with certain residues and structural domains preferentially recognizing specific lipid types. These findings underscore the relevance of CFTR’s lipid-facing basic and polar residues in mediating membrane stabilization, potentially influencing conformational dynamics and coupling with ion transport.

### Coupling TM helix irregularities to non-covalent interactions

We next examined whether the identified electrostatic interactions (SBs and HBs) correlate with local secondary structure features, focusing on irregularities within the TM helices and their behavior during MD simulations. **Table S2** lists the amino acids which exhibited transient or persistent deviations from an ideal helical geometry, defined as maintaining less than 80% α-helix occupancy throughout the MD simulations (α-helix occupancy is indicated for each MD simulation). These regions (blue ribbons in **Figure 6**) include charged and polar residues, whose side chains are all involved in electrostatic contacts (see the interaction partners in **Table S2**), accounting for approximately 16.1% of all detected interactions and 19.4% of those located in the TM helices. Several of the affected residues also participated in contacts with anions, as specified in **Table S2**. Remarkably, these residues cluster into three distinct regions (encircled in **Figure 6**) (i) the end of TM helices belonging to the ECLs, (ii) the region around the TM8 unwound segment and (iii) a region located at the level of the internal vestibule and portals, with a particularly pronounced deformation of TM3 and TM9 (inset in **Figure 6**). Thus, residues located in helix irregularities often coincided with electrostatic contacts. In other words, when a charged amino acid side chain reaches out to form a SB, an alpha-helix can locally unwind or kink to accommodate that interaction.

**Figure 6:**
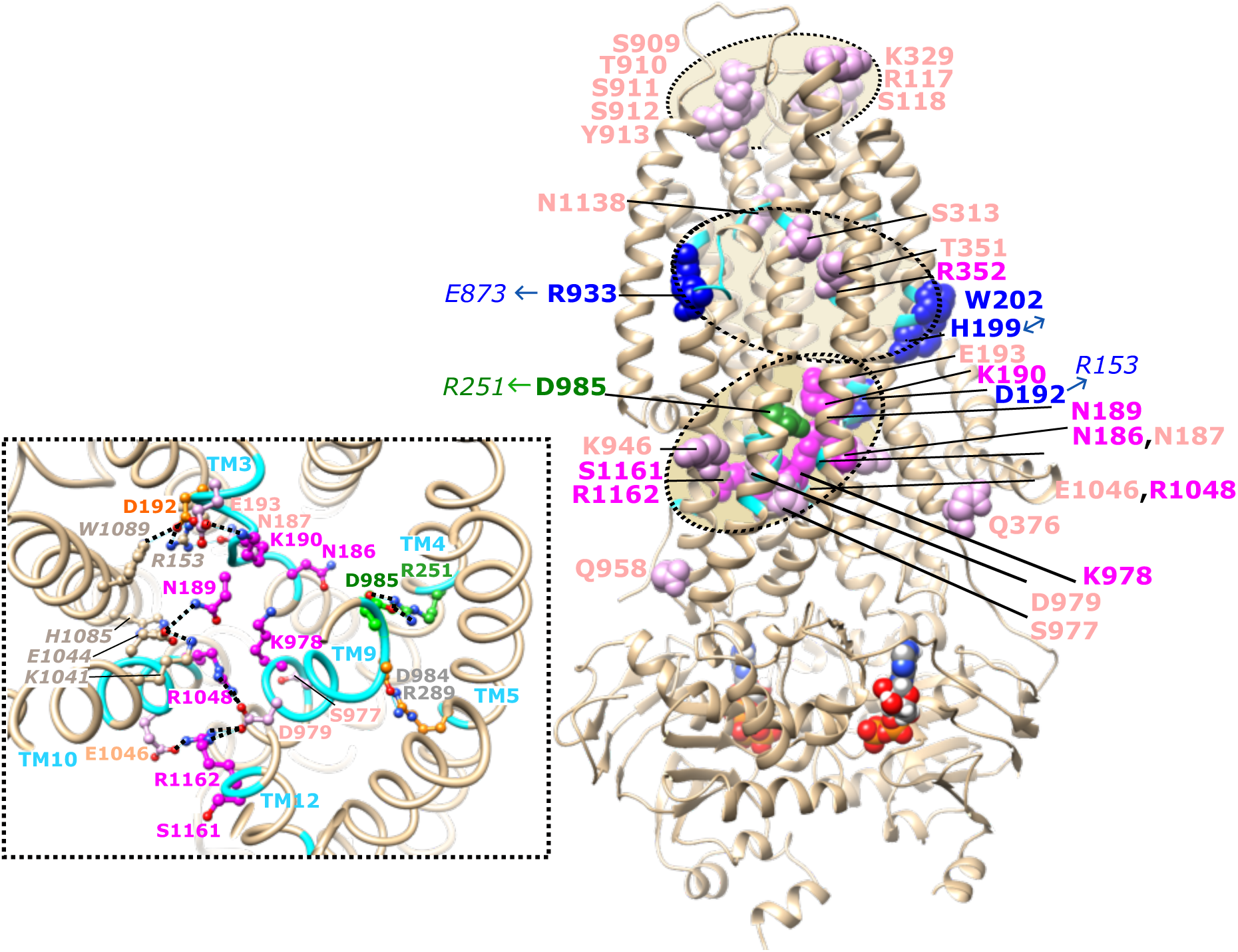
Charged and polar side chains involved in TM α-helix irregularities. The Cα atom positions of charged and polar residues included in α-helix irregularities, presented in **Table S2**, were mapped on the human CFTR 3D structure after 500 ns of MD simulation (apo condition, replica 1). They are colored according to their participation in electrostatic interactions that are stable (with (green) or without (blue) contact with anions) or transient (with (magenta) or without (pink) contact with anions). The partners of amino acids involved in stable interactions are shown in italics at the tips of single-headed arrows, whereas the double-headed arrow indicates the stable interaction between H199 and W202. The polar/charged residues included in irregularities cluster into three distinct regions (tan shaded), with that corresponding to the inner vestibule depicted in more details in the inset (amino acids involved in the interaction network but outside irregularities are depicted in grey).

A notable observation is that electrostatic interactions associated with anion contacts and integrating such local irregularities (green and magenta in **Figure 6**) were almost always less stable (mean frequency < 0.8, magenta in **Figure 6**), with the exception of the highly stable R251_TM4_–D985_TM5_ pair (0.98), which engages in anion binding (green in **Figure 6**). Conversely, several SBs that do not participate in any anion interactions exhibited very high stability, often a frequency above 0.98 - notably R153_TM2_–D192_TM3_ and R933_TM9_–E873_TM7_ — suggesting a purely structural role. Interestingly, HBs involving S or N displayed frequencies below 0.7, supporting the idea that they contribute to more transient or flexible interactions. Taken together, these observations suggest a trade-off between stability and functional adaptability, on a framework where the helices adapt by deviating locally from the alpha-helical pattern: highly stable electrostatic interactions may reinforce CFTR architecture, while less stable, dynamically switching interactions may support functional tasks like ion coordination or membrane coupling. Overall, integrating the secondary structure analysis with our interaction data highlights some residues as multi-faceted hubs where structure and function intersect: they are loci of helix deformability, intra-protein anchoring, and external ion/lipid engagement, all of which may be critical for CFTR’s mechanistic flexibility.

### Impact of VX-770 binding on electrosta tic interactions

#### The VX-770-binding site

The cryo-EM 3D structure of human CFTR in complex with the potentiator Ivacaftor (VX-770) has highlighted a binding site at the protein-lipid interface, docked into a cleft formed by transmembrane helices TM4, TM5 and TM8, and involving the unwound segment of TM8 ^17, 65^. The binding pocket mainly involves aromatic and aliphatic hydrophobic residues. Our MD simulations highlighted the participation of lipids in the stabilization of the drug into the binding site. Most of the key interactions observed in the cryo-EM 3D structure were conserved during all the MD simulations (**Figures 7A (**highlighted in green) and **7B** (green stars)). Lower-frequency or conditional contacts (detected in some simulations) included several other amino acids of TM4, TM5, TM7 and TM8 (**Figures 7B**). In one of the MD simulations, VX-770 slightly shifts towards TM7, establishing novel contacts with E873_TM7_ (HB with VX-770 phenolic hydroxyl), which forms a strong SB with R933_TM8_. This R933_TM8_–E873_TM7_ SB appears thus to serve a dual function: stabilizing the architecture of the binding pocket (including the unwound segment of TM8) and directly contributing to ligand binding and stabilization. Our MD simulations also revealed a noticeable effect of VX-770 on the structural integrity of TM8. In all the trajectories in the presence of VX-770, the region spanning G930_TM8_ to R933_TM8_ tends to a more stable helical conformation, whereas in all Apo simulations, this segment displayed clear irregularities and a tendency to unwind (**Table S4**). This observation suggests that VX-770 not only anchors within the TM8 cleft but also contributes to stabilize the local secondary structure, thereby potentially supporting its role in CFTR gating modulation.

**Figure 7:**
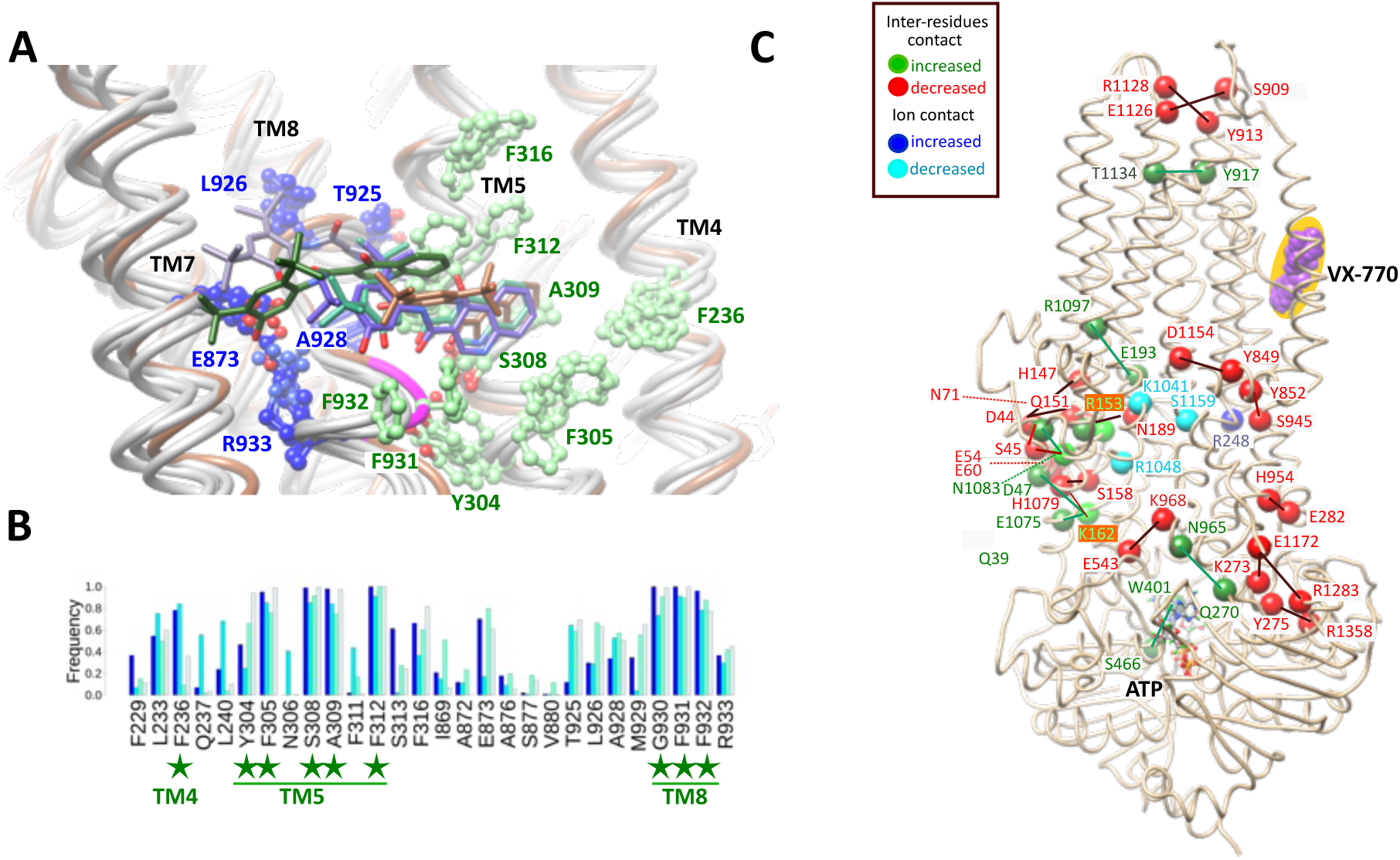
VX-770 binding and effect on the electrostatic interactions. **A.** View of the VX-770 binding site and the main contacts that the drug establishes with amino acids from TM4, TM5, TM7 and TM8, depicted in atomic details. The 3D structures at the end of the ligand-bound MD simulations were superimposed to the experimental 3D structure of human CFTR, in complex with VX-770 (pdb 6OP2, colored in brown). Conserved contacts relative to the cryo-EM 3D structure are shown in green, while novel contacts (VX1 MD simulation) are depicted in dark blue (VX-770 is shown in purple). The 930-933 segment, which is in helical conformation only in the VX-bound conditions, is colored in magenta. **B.** Barplot displaying the frequency of contacts between CFTR amino acids and VX-770 during the four ligand-bound MD replicates: VX1 (blue), VX2 (cyan), VX3 (aquamarine) and VX4 (azure). Each bar corresponds to a residue that formed a contact with VX-770, and the y-axis indicates the fraction of simulation time the contact was maintained. **C.** View of the CFTR 3D structure (pdb 6O2P), on which are reported amino acids (Cα atoms only) involved in electrostatic interactions and whose contact frequencies (%) differ significantly (≥ 25% for side-chain/side-chain interactions, ≥ 20% for interactions of side chains with ions) between the apo and VX770-bound conditions, as reported in **Table S4**. The pairs involved in side-chain/side-chain interactions are linked with bars. Some amino acids shown in light green participate in different bonds, which are respectively increased and decreased in the VX-770-bound conditions.

#### Modulation of inter-residue electrostatic interactions by VX-770

We next compared the prevalence of each electrostatic interaction between the Apo and VX-770–bound conditions to detect potential drug-induced shifts. **Figure S12A** plots the mean contact frequency of each electrostatic interaction between amino acid side chains in the absence (x-axis) versus presence (y-axis) of VX-770. Most data points lie along the diagonal, indicating that the majority of electrostatic interactions are maintained at comparable frequencies across both conditions. This overall conservation suggests that VX-770 binding does not globally alter CFTR’s interaction network, consistent with its mode of action as a potentiator that enhances channel function without inducing large-scale structural rearrangements.

However, several notable exceptions were identified, reported in **Table S3**, with outliers with contact frequency differences greater than 25% labeled in **Figure S12A** and highlighted in green/red on the 3D structure shown in **Figure 7C**. Bonds with significantly increased and decreased frequencies (green and red, respectively in **Figure 7C**) are concentrated into specific areas relative to the membrane plane, at the level of (i) the ECLs, ii) the TMD:lasso interface including ICL1 and ICL4, iii) the ICL2-ICL3:NBD2 interface. Interestingly, in the extracellular region, frequency increase of T1134_ECL6_–Y917_ECL1_ bond (+36%) coincided with loss of the R1128_ECL6_–Y913_ECL1_ (-34%) and E1126 _ECL6_-S909_ECL4_ interaction (-29%), potentially facilitating opening of the outer vestibule to support ion permeation.

Altogether, these observations support a model in which VX-770 enhances CFTR gating by allosterically modulating dynamic interactions across gating-relevant regions.

#### Changes in anion and lipid contacts upon VX-770 binding

We also examined whether VX-770 binding influences the interaction of CFTR’s charged and polar residues with anions and lipids (**Table S3, Figure S12B-C**). Only one residue exhibited a substantial increase (15%) in anion contact in the presence of VX-770, notably (R248_TM4_, dark blue in **Figure 7C, Table S3, Figure S12B**). This residue is located just behind the R249_TM4_-K370_TM6_ SB at the entry of the TM4/TM6 portal. Conversely, three last residues in the vicinity of to the TM10/TM12 portal (K1041_TM10_, R1048_TM10_, and S1159_TM12_) show a slight decrease. This suggests that VX-770 binding may have an effect, albeit modest, on the anion attraction at the portals.

A parallel comparison for lipid headgroup contacts (analyzed by lipid type) is shown in **Figure S12C** and **Table S3**. Most residues exhibited minimal differences, suggesting that CFTR’s overall membrane association remains largely unchanged upon VX-770 binding. However, some residues at the protein–lipid interface displayed notable positive shifts. Contacts with POPS globally decreased (strong effect for R21_lh1_/ R25_lh1_ / R1102_TM11_ at the lasso:TMD2 interface) in favor of an increase with other lipids, in particular POPE (strong increase of contacts with R21_lh1_ / R25_lh1_ / R29_lh1_ at the lasso/TMD2 interface and R74elbow1 / R75_elbow1_) and POPC (strong increase of contacts with R80_TM1_ / R242_TM4_ / R1102_TM11_) (**Figure S12C, Table S3)**. These shifts suggest that VX-770 may subtly remodel lipid engagement at specific sites, in particular at the level of the lasso motif and elbow helices, consistent with the modification of the network of interactions between amino acids described above.

#### Effect of VX-770 on the portals/extracellular exits

The two portals and the two exit pathways discussed above do not appear to be particularly favored in the presence of the potentiator (**Table S4**). The analysis of water flux and ion occupancy within the membrane slice (centered on the selectivity filter region) did not reveal a statistically significant effect of VX-770 in the conditions of its observed position, as the differences between apo and VX trajectories remained within error bars. The TM1-TM6 exit, in the passage of which F337_TM6_ is found, is either non present, small or larger and conditioned by a F337_TM6_ rotamer, while the ECL1-ECL6 exit is not observed (**Table S4**). Notably, some trajectories displayed stable and stronger contacts not only with E873_TM7_ and R933_TM8_, which together form the anchoring salt bridge of the VX-770 binding pocket, but also with surrounding residues of the unwound TM8 segment, including T925_TM8_, L926_TM8_, A928_TM8_, and M929_TM8_. This suggests that enhanced stabilization of the TM8 region by VX-770 may facilitate ion capture, thereby representing an early premise of channel opening.

## Discussion

In this comprehensive study, we have performed microsecond-long, all-atom unbiased molecular dynamics (MD) simulations of the human CFTR 3D structure embedded in a mixed and asymmetric lipid bilayer to investigate how electrostatic interactions involving amino acid side chains — specifically salt bridges and hydrogen bonds — shape the protein’s architecture, dynamics and functions. By comparing apo and VX-770–bound states, we provide mechanistic insights into how electrostatic networks (protein-protein, protein-ion and protein-lipid interactions) and transmembrane helix irregularities may contribute to CFTR gating and potentiation. Our findings illustrate several key features that cannot be captured by static cryo-EM 3D structures alone, including the dynamics of bond switching, the influence of the lipid environment, and the coupling between helix deformability and specific stable or transient interactions.

We identified 557 electrostatic interactions between amino acid side chains across CFTR domains, which we classified by frequency and location on the 3D structure. We focused our analysis on the most frequent ones (337 with a frequency ≥ 0.25) and distinguished stable from transient interactions. Many highly stable bonds allow specific interactions between TM helices that are particularly strong in the low dielectric environment of the membrane ^67^, and enable the anchoring of domains in relation to one another—particularly at the level of the elbow, lasso, and ICLs:NBDs interfaces. These stable, “architectural” salt bridges are consistent with the “connecting” principle proposed for immune receptors ^67, 68^ where such interactions constrain conformational flexibility and maintain global fold integrity ^69^. Several arginine residues involved in stable SBs, such as R347_TM6_ and R933_TM9_, establish additional hydrogen bonds with main chain atoms (for R347_TM6_ with F310_TM5_ and F311_TM5_ and for R933_TM9_ with C866_TM7_ and F870_TM7_). This allows adding structural rigidity, a property already observed in other systems ^70^. In contrast, dynamic or less stable electrostatic interactions are concentrated in regions where functional transitions related to gating may occur, such as the TM4/6 and TM10/12 portals. The data presented here indicated openings at both portals, with a transient salt bridge (R249_TM4_-K370_TM6_) that may modulate access to the main TM4/TM6 portal, whose role in anion flux has been demonstrated experimentally ^15^. While experimental data currently available attributed only a minor role to the TM10/12 portal ^15^, anion binding was also observed at this level, with a possible modulation by lipids. On another hand, the “functional bonds” (as opposed to the “architectural” term used above) fluctuate during simulations and often involve basic residues that also contact chloride or bicarbonate ions. This mirrors mechanisms described in other transporters and ion channels, where salt bridge switching has been shown to underlie conformational transitions or channel activation (*e.g.* references ^71–74^). It is important to note that several amino acids, notably N189_TM3_, N974_TM9_, D979_TM9_, D984_TM9_ and R1158_TM12_ (**Table S5**), have been identified here as central hubs, exhibiting a large number (≥ 5) of electrostatic interactions.

A striking result of our analysis is the tight correlation between electrostatic interactions and transmembrane helix irregularities—regions where α-helical geometry is locally disrupted. These include π-helix bulges, 3₁₀-helix constrictions, or unwound segments, which are increasingly recognized as evolutionarily conserved “weak spots” enabling conformational changes due to a larger plasticity ^75^. In CFTR, we show that most residues located in irregularities are directly involved in electrostatic interactions, supporting a model where local electrostatic interactions act on helix flexibility to modulate gating functions. TM irregularities associated with electrostatic interactions cluster into three distinct regions, namely the ECLs (ends of the TM helices flanking the loops), the intracellular vestibule/portals region and the unwound TM8 segment. The inherent flexibility associated with non-stable electrostatic interactions in the intracellular vestibule/portals region is consistent with its role in anion coordination and permeation. Interestingly, this region also includes several gain-of-function variants affecting basic residues (e.g. K978_TM9_ ^76^, R1030_TM10_ and R1158_TM12_ ^77^), supporting the fact that the electrostatic interactions in which they are involved play a key role in potentiation mechanisms. The central, unwound segment of TM8 contributes to CFTR’s structural asymmetry and was proposed to support gating-specific rearrangements ^17, 78^. Remarkably, this unwound segment has a very stable conformation all along our MD simulations, as it was also observed in previous MD studies ^30, 31^. This region is flanked by two salt bridges, which also proved to be particularly stable in our MD simulations: R347_TM6_–D924_TM8_ and R933_TM8_–E873_TM7_. Comparison of the experimental 3D structures of ATP-free and ATP-bound CFTR ^79^, as well of CFTR in the presence of the inhibitor CFTR_inh_-172 (bound at the same level than VX-770 but within the pore ^80, 81^) indicated that the extracellular portions of TM8 (ending at D924_TM8_), TM12 and TM1 are flexible and mobile, representing key actors in the CFTR gating apparatus. D924_TM8_ may thus act as a pivot around which conformational transitions occur at the extracellular side, while maintaining a stable base on the unwound segment pillar. Interestingly, we find that the R933_TM8_–E873_TM7_ bridge is consistently present (100 % frequency) in both apo and VX-770-bound MD simulations, suggesting a critical role in stabilizing the TM8 unwinding and maintaining the architecture of this region, thereby contributing to the integrity of the VX-770 binding pocket. Moreover, we also highlighted in our MD simulations different positions of the extracellular portions of TM8, coupled with movements of TM12 and TM1, and with the appearance of clear exits towards the extracellular milieu (TM1-6 and ECL1-6 exits, **Figures 4B/D**), suggesting a transition towards an open channel. Notably, in a work that predates the resolution of CFTR cryo-EM structures, the R347_TM6_–D924_TM8_ interaction was shown to be essential for open-state stabilization ^41^, along with another salt bridge, R352_TM6_-D993_TM9_. While the latter is visible in cryo-EM 3D structures, it appears less stable in our simulations. Instead, D993_TM9_ preferentially pairs with R303_TM5_, leaving R352_TM6_ more available for anion coordination. Intriguingly, R352_TM6_ is also located within a helix irregularity (a π-helix segment), reinforcing the notion that such residues act as regulatory switches whose availability and functional role depend on dynamic bond engagement.

Although VX-770 did not globally alter the CFTR salt bridge landscape, it did have an effect on specific networks, while it also stabilized the TM8 α-helical structure following its unwound segment (amino acids 930-933). Interestingly, VX-770 has an effect on inter-residue contacts at the level of the TMDs:lasso and ICLs:NBDs interfaces, suggesting long-range allosteric pathways that may modulate communications with the cytoplasmic domains. Effects are also observed on the lipid interaction network, especially at the level of the lasso motif and TM11, indicating a role for peripheral elements near the membrane in stabilizing gating-relevant conformations. Among the involved amino acids are S18_lasso_, R21_lasso_, W1098_TM11_ and R1102_TM11_, which are known to directly contact VX-445 (Elexacaftor) ^66^, supporting a potential convergence of VX-770 and VX-445 potentiation mechanisms in this region. In addition, some contact frequencies are modified in the drug-bound state at the level of ECLs, potentially contributing to opening of the outer vestibule and promoting ion efflux. The likely important role of ECLs in the gating mechanism was already pointed out in the work of Simon and Csanady ^27^. In our analysis, their role is all the more obvious here, as the long ECL4 is considered in its entirety. Despite this and although some first insights have been gained on the dynamics of lateral portals and extracellular exits, the studies presented here only lead to transient openings of the channel, which undoubtedly requires additional factors to be taken into account. N-glycosylation of ECL4 could be a critical factor, in addition to its role in CFTR folding and trafficking ^82^.

Compared to ABCC transporters and its closest paralog ABCC4 ^23, 83^, CFTR has evolved through a number of pathways to adopt a regulated channel behavior (reviewed in ^10^). Notably, Infield et al. ^10^ demonstrated that the CFTR chloride channel evolved by utilizing residues that were already functionally important in the transport of organic anionic substrates in ABCC transporters (i.e. K95_TM1_, R248_TM4_, R303_TM5_, K370_TM6_, R1030_TM10_, K1041_TM10_, R1048_TM10_), and repurposing them for the new function of conducting inorganic anions through the channel pore. This is consistent with the overall conservation of pore-lining residues of several TMs ^10^, with the notable exception of TM6, a highly discriminatory feature of CFTR that may account for neofunctionalization ^84, 85^. The highly divergent TM6 is directly involved in the formation of the lateral TM4/6 portal at the level of the intracellular loops, allowing anion to flow from the cytoplasm to the internal vestibule. TM6 also contains two arginine residues (R347_TM6_ and R352_TM6_), which are involved in electrostatic interactions with surrounding helices (with D924_TM8_ and D993_TM9_, respectively), although as we have shown here, R352_TM6_ is more frequently left free in favor of the interaction of D993_TM9_ with R303_TM5_. These salt bridges are included in a key network specific to CFTR at the level of the unwound segment of TM8 (another specific feature of CFTR already discussed above), also including the stable R933_TM8_-E873_TM7_ salt bridge. Remarkably, only one of the two partners in each of these electrostatic interactions is conserved in ABCC4 (R347_TM6_, D993_TM9_ and R933_TM8_), while the second ones are V, L and Q, respectively (**Table S5**). The emergence of these electrostatic interactions in CFTR has been proposed to stabilize the open conformation of the pore, while we evidence here that R352_TM6_ likely plays a direct, key role in the regulation of anion conduction. In line with this observation, it was proposed that the positive charge of R352_TM6_ contributes to an electrostatic potential in the channel that forms a barrier to cation permeation ^86^, while reduced short-circuit current is observed with the rare R352Q mutation ^35^. Overall, beyond these critical SBs, we show here that CFTR has developed a specific network of electrostatic interactions, involving residues which are only partially conserved in ABCC4 (**Table S5**), suggesting the acquisition of novel functional features from an ancestral network linked to organic anion binding. Thus, it is likely that the larger and more complex network of electrostatic interactions of CFTR contributes, on the one hand, to stabilize the open conformation of the channel and, on the other hand, to regulate its activity. With this specific feature of CFTR clearly highlighted, it is also interesting to note the conservation, between ABCC4 and CFTR, of the extensive interaction networks around N189_TM3_ and N974_TM9_ at the level of the intracellular vestibule (**Table S5**), highlighting their likely key role in the architecture of a degraded intracellular gate ^11^.

Beyond interactions linking amino acids together, our study highlights the importance of CFTR’s coupling to ions and to its membrane environment, revealing among other amino acids which are involved in multiple types of interaction (**Figure S13**). Limited information is available about this environment and its influence on CFTR function and regulation. A few works have shown that CFTR is located in specialized membrane, detergent-resistant domains at the cell surface, assembled from clustered lipids - principally cholesterol and sphingomyelin ^87–89^. Moreover, a few studies have provided evidence of a direct influence of lipids on the intrinsic CFTR activity (phosphadylserine ^64^, cholesterol ^62, 63^). Here we showed that specific basic and polar residues engage in frequent contacts with cholesterol and phosphatidylserine (POPS), forming distinct lipid-binding patches. Notably, the lasso domain and elbow regions exhibited high POPS occupancy, consistent with the established role of these CFTR regions in membrane insertion ^2^. A TM4–TM5 region, located near the VX-770 binding site, also displayed organized cholesterol and POPS interactions, particularly involving R242_TM4_, R246_TM4_, and R297_TM5_, suggesting a membrane-mediated pathway for allosteric communication between the drug-binding cleft and the transmembrane core. Several of these residues do not participate in salt bridges or ion contacts, reinforcing their specialized role in lipid coupling. Besides direct binding coupled to allosteric communications, lipids can impose mechanical forces that affect the protein function, especially by modification of the membrane fluidity (for a review see ^90^). One may also wonder about the role played by the lasso, a segment largely inserted into lipids and interacting strongly with their polar heads, on the properties of the membrane, in particular its curvature. By considering an asymmetric membrane with a five-component system, our study thus takes a step forward in understanding the complex relationships between CFTR and the membrane, compared to computational studies that have examined the protein in a single-component system ^14, 30–35^. Nevertheless, it remains true that this system is still fairly simple, not yet incorporating, for example, sphingolipids and phosphatidylinositol lipids, which play a central role in the organization and polarization of epithelial cells ^91^.

Overall, this comprehensive mapping of the CFTR interaction network provides valuable insight into the mechanisms of channel activation. Relatively few amino acids of this network are subject to missense mutations (CF-causing (CF) or with varying clinical significance (VCC), **Table S5**). They include, in the TMDs, amino acids from the ECLs (D110H/E, E116K, R117H/C/L/G/P, R334W/L/Q, S1118F), key amino acids of the TMD central core (E92K, Q98R, H139R, S341P, R347P/H and D924N (both amino acids interacting together), R352Q/W, D1152H) and at the level of the inner vestibule/portals (D192G, R258G, S977F, D979V, H1085P, S1159F/P). Also affected are residues linking TMD2 to lasso helix 1 (Y1032C, W1098R), belonging to the VX-445 (Elexacaftor)-binding site ^66^, as well as a set of amino acids linking the lasso and elbow helix of TMD1 (E56K, E60K, R74W), a region that has been shown to be essential for folding and stabilization by the correctors VX-809/VX-661 ^42, 43, 92^. Finally, several mutations are also found at the ICLs:NBDs and NBD1:NBD2 interface (**Table S5**). Most of the variants for which information is available in the refined classification proposed by Veit and colleagues ^93^, which accounts for complex phenotypes of CFTR major cellular defects, correspond to pure or combined class II mutations (**Table S5**), indicating that the corresponding amino acids play critical roles in the CFTR structural integrity. Two class III mutations (D110E and R347H) correspond to conservative substitutions, whereas non-conservative ones affecting the same amino acids (D100H and R237P) are also associated with class II defects. The class III mutation R352Q stands out in this list because, as indicated above, it affects an amino acid that forms a transient salt bridge with D993, but the latter amino acid is more frequently linked to R303, leaving R352 available to interact with anions. On another hand, several of the amino acids subject to missense mutations are strictly conserved in the ABCC4 sequence (**Table S5**), further supporting that they play critical roles in the ABC architectural template. In conclusion, a detailed analysis of the CFTR variants, including those affecting a same amino acid but with different effects (*e.g.* S1159F and S1159P ^94^), in light of the networks described in this study could lead to a better understanding of the multiple mechanisms underlying their dysfunctions. For instance, the class II-III S945L variant in TM8, which has been reported to affect folding due to disruption of an H-bond with Y852_elbow2_ (^33^, **Figure S4-C**) may also affect, through its transient interaction with D984_TM9_ (**Figure S4-A**), the central interaction network at the level of the inner vestibule/lateral portals, thereby affecting structure and/or function. Our data also show a significant effect of VX-770 on the Y852_elbow2_-S945_TM8_ H-bond (**Figure 7C, Figures S4-C, S12-A, Table S3**), which may account for the allosteric mechanisms linked either to potentiation or to the destabilization of the CFTR protein ^95, 96^. Our study could thus open up interesting avenues for the rational design of targeted-pharmacological approaches to fully restore channel function for these mutations and, more generally, to stabilize the open state regardless of the mutation considered.

This work also illustrates the power of MD simulations to uncover structural and dynamic features that are invisible to static experimental techniques. We capture not only the stability of specific electrostatic interactions but also their switching behavior, interactions with lipids and ions, and their coupling to helix plasticity - all of which are essential for mechanistic understanding. Nonetheless, some limitations remain. Forcefield inaccuracies may affect the fine energetics of anion coordination and conduction. Importantly, the additive force fields used here are not designed to simulate ATP hydrolysis or to accurately capture large-scale conformational changes (such as the transitions toward the inward-facing state starting from the ATP-bound cryo-EM structure), or events such as helix winding and unwinding, which are likely critical for CFTR gating. Moreover, while our simulations show clear openings of the channel, we only observed transient conduction states. Future studies incorporating enhanced sampling methods, longer simulation timescales, or polarizable force fields could help address some of these limitations, as could the consideration of new parameters that have not yet been taken into account. A pragmatic low-cost alternative to explicit polarization is the use of electronic continuum correction (ECC), including uniform charge scaling in non-polarizable force fields ^97^. This approach can mitigate ion overbinding and improve transport properties when used consistently with an ECC-compatible water model, though care is needed to avoid overscaling artifacts. On the other hand, inclusion of phosphoinositides like PIP2 (phosphatidylinositol-4,5-bisphosphate) in the membrane bilayer, which have been implicated in CFTR modulation of gating ^98, 99^, may help complete the picture of membrane-driven modulation. Finally, as pointed before, proper N-linked glycosylation of ECL4 may also participate in the modulation of CFTR gating, while an arginine (R899) in the direct vicinity of one of the glycosylation sites is directly involved in the CFTR activity as an extracellular chloride sensor ^61^. Glycosylation could thus be necessary to stabilize the open state of CFTR, so explicitly modeling glycan chains in future simulations would be informative. One final point that should not be overlooked in the context of electrostatic interactions is the R domain, which simulations do not take into account but which establishes multiple phosphorylation-dependent intramolecular bonds ^100^.

## Materials and methods

### System preparation and molecular dynamics simulations

All molecular dynamics (MD) simulations were performed starting from a high-resolution cryo-EM structure of human CFTR in the ATP-bound, phosphorylated (E1371Q) conformation (outward occluded conformation) (PDB ID 6O2P), solved at ∼3.3 Å resolution in presence of the potentiator VX-770 (ivacaftor) ^17^. The ten missing amino acids of ECL4 (from Q890 to R899), located upstream the C-terminal extended structure in contact with ECL3, were added manually, and the structure was subsequently submitted to energy minimization and molecular dynamics simulation (see below). This procedure allowed to obtain a starting conformation of the ECL4 loop, in which D891 and R899 form two salt bridges (2.86 Å and 3.81 Å), the latter being also linked to the G893 carbonyl group through an H bond (2.79 Å). All titratable residues were assigned their standard protonation states at pH 7.4. The protein was embedded in a heterogeneous and asymmetric phospholipid bilayer composed of POPC (1-palmitoyl-2- oleoylphosphatidylcholine), POPE (1-palmitoyl-2-oleoylphosphatidylethanolamine), CHL1 (cholesterol), DSM (sphingolipid) and POPS (1-palmitoyl-2-oleoylphosphatidylserine) – at an approximate ratio of 3:3:2:1:1 as proposed by ^28^. The protein was positioned within the membrane bilayer using the PPM (Positioning of Proteins in Membranes) server ^101, 102^. The system was assembled using the CHARMM-GUI membrane builder workflow ^103, 104^. Physiological ionic strength was achieved by adding 150 mM NaCl. In addition, a small and variable amount of sodium bicarbonate (NaHCO₃) was included to represent the presence of bicarbonate anions in the solution, given CFTR’s known bicarbonate conductance (**Table S1**).

The system was parameterized with the CHARMM36m force field for the protein, lipids and ions, and TIP3P water molecules were used ^105^. The VX-770 parameters were calculated using the CHARMM General Force Field (CGenFF) ^106^ in CHARMM-GUI ligand modeler module ^107^. Energy minimization (5000 steepest descent steps) was first performed to remove bad contacts, followed by a six-stage equilibration protocol. The first three equilibration steps were performed under NVT conditions, followed by the last three steps under NPT conditions, all at a temperature of 310 K. Position restraints on the protein heavy atoms (and VX-770, in the bound cases) and the lipid heads were applied initially and slowly released over the 6 stages (∼1.9 ns), allowing the membrane and protein to relax and the solvent to equilibrate. Production runs were then carried out in the NPT ensemble at 310 K, 1 bar with periodic boundary conditions.

Simulations performed with GROMACS (versions 2020.3 and 2023.2) ^108^ employed Nose–Hoover thermostat and Parrinello–Rahman barostat ^109, 110^ or, alternatively, the V-rescale and C-rescale algorithms ^110^, with LINCS constraints on bonds to hydrogen ^111^. The use of different thermostat–barostat combinations allowed us to confirm that our results are consistent across coupling schemes, which differ mainly in numerical stability, relaxation timescales and damping but are equivalent in their ability to sample the NPT ensemble. A 2 fs integration time step was used in all cases, and long-range electrostatic interactions were computed with the Particle Mesh Ewald method ^112^. Eight independent MD trajectories were generated in total: four for the apo CFTR (Apo1, Apo2, Apo3, Apo4) and four for the VX-770–bound state (VX1, VX2, VX3, VX4). Simulations were run either for 500 or 1000 ns, yielding an aggregate simulation time of 6 µs for analysis (**Table S1**).

### Contacts analysis

Electrostatic contacts involving amino acids, anions, lipid headgroups, and VX-770 were calculated using the VLDM (Voronoi Laguerre Delaunay for Macromolecules) program ^113, 114^. VLDM relies on a tessellation method—that is, a partition of space into a collection of polyhedra that fill space without overlaps or gaps. Delaunay tessellation and its Laguerre dual were performed using a set of heavy-atom Cartesian coordinates and weights based on the van der Waals radii of the atoms, as defined by the CHARMM36m force field. Two atoms are considered to be in contact if they share a common polygonal face within the tessellation, allowing for a rigorous, geometry-based definition of interatomic interactions. The contact interface between molecular groups is quantified as the sum of the polygonal surface areas shared between their respective atoms.

For the analysis of the contacts, we focused on amino acids typically engaged in electrostatic interactions—namely arginine (R), lysine (K), glutamic acid (E) and aspartic acid (D), and we also included histidine (H), serine (S) and threonine (T), as well as asparagine (N), glutamine (Q), tyrosine (Y), tryptophane (W) and cysteine (C). Salt bridges (SBs) and hydrogen bonds (HBs), were defined as side chain-side chain electrostatic interactions between nitrogen or oxygen atoms of these residues. In addition, we also quantified side chain (N/O atoms of R, K, H, S, T, D, E, N, Q, Y, W, C) to main chain (backbone N/O atoms) hydrogen bonds, considering only residue pairs separated by more than five amino acids along the sequence. Anion contacts were identified when basic residues (R, K, H) and S/T/N/Q/Y/W shared a tessellation face with non-carbon heavy atoms from bicarbonate (HCO₃⁻) or with chloride (Cl⁻) ions. Lipid interactions were similarly defined as contacts between the same residues and non-carbon heavy atoms from the headgroups of lipid molecules. VX-770 contacts were calculated by considering all heavy atoms of both the ligand and all the protein residues.

For each simulation frame, VLDM generated contact data, which were aggregated over time using in-house R scripts to compute the frequency of each contact, expressed as the percentage of frames in which the contact occurred. These scripts also enabled the combination of contact frequency data across all simulation replicas, the export of the combined data as CSV files, and the generation of comparative bar plots.

The resulting tables were further analyzed using in-house Python scripts. First, contact frequencies were averaged across all simulation replicas. To focus on relevant contacts, only those with a mean frequency equal to or greater than 10% were retained. Next, the distribution of averaged electrostatic interactions across the regions in the CFTR 3D structure was determined (see next subsections), and overlaps between residues forming electrostatic interactions and those involved in anion or lipid contacts, as well as with those involved in alpha-helix irregularities, were calculated. This integrative analysis enabled a detailed annotation of the electrostatic interactions.

### CFTR region classification by z-coordinate

To analyze results by protein region, a coordinate-based scheme was applied to assign each residue (and each electrostatic interaction) to a structural/functional domain of CFTR along the Z-axis. The Z-coordinate of each residue’s Cα atom involved in an electrostatic interaction was extracted from a CFTR PDB structure and used to determine its regional assignment. Five regions were delineated based on natural breaks in the CFTR structure: (1) NBDs: residues with Z < 30 Å; (2) ICLs:NBD interface: residues with Z ≈ 30–53 Å, covering the ICLs and adjacent regions of the NBDs in contact with the TMDs; (3) Elbow/Lasso/ICLs region: Z ≈ 53–65 Å, including the elbow helices, lasso motif at the N-terminus, and cytoplasmic ends of transmembrane helices near the inner leaflet; (4) TM helices/Membrane: residues within the lipid bilayer, with Z ≈ 65–105 Å, corresponding to the membrane-spanning core; and (5) ECLs: residues with Z > 105 Å, located on the extracellular side, including loop segments connecting TM helices. Using this scheme, each electrostatic interaction was assigned to one of the defined CFTR functional regions.

### Secondary structure and TM helix irregularity analysis

To probe secondary structure dynamics, we used the DSSP algorithm ^115^ via *cpptraj* ^116^ to assign the secondary structure of each residue within CFTR’s transmembrane helices (TMs 1–12) throughout the trajectories. For each simulation, we calculated helical occupancy—the fraction of frames in which a residue was assigned as α-helix (as opposed to turn, coil, 3₁₀-helix, etc.). Residues with consistently high occupancy (≈100%) were considered part of stable helical regions, while those with occupancy below 80% (showing variability) were flagged as helix irregularity sites.

**Table S2** summarizes these results, listing all R, K, H, D, E, S, T, N, Q, Y, W, C residues within TM helices that exhibited <80% helical occupancy in any simulation. For each, the table reports helical occupancy across all simulation replicas (Apo1–4, VX1–4) and indicates whether the residue participated in electrostatic interactions and/or made notable contacts with anions. This integrative analysis allowed us to correlate the helix irregular sites with the different types of interactions.

### Data processing and visualization

Trajectory analyses were performed using a combination of VMD Tcl scripts and in-house Python scripts. In the Python workflow, MDAnalysis was used to manage trajectory input/output ^117, 118^. Additional analyses, such as averaging contact frequencies across replicates and identifying significant differences between conditions, were carried out using NumPy and pandas in Python. To compare apo and VX-770 conditions, the apo trajectories were treated as independent samples, as were the VX-770 trajectories. Mean values were computed, and Δfrequency > 0.3 were considered notable given the sampling length.

Bar plots, scatter plots, and overlap analyses (based on residue list intersections) were generated using Matplotlib, with consistent color schemes applied to represent regions and contact types. Three-dimensional molecular visualizations were created with PyMOL ^119^ or Chimera ^120^. Residue mapping was based on coordinates from the initial cryo-EM structure (PDB ID: 6O2P). PML files were generated to color residues according to their structural region or their involvement in specific types of interactions.

## Acknowledgments

This work was performed using HPC resources from GENCI-[CINES] (Grants 2017-A0020707206, 2018-A0040707206, 2019-A0060707206 and 2020-A0080707206). It has been supported by institutional grants from CNRS and INSERM, and benefited from financial supports of the French Association Vaincre la Mucoviscidose.

## Data and Software availability

MD simulations data (pdb files and trajectories / topology and parameters files / GROMACS entry files), analysis data (contact frequencies, secondary structures) and scripts for analyzing them are provided on Zenodo at https://doi.org/10.5281/zenodo.18922210. The jupyter-lab notebook (image generation from tsv files) is also provided. The VLDM program and the scripts used to generate contact frequencies must be requested from the program author (Christophe Oguey).

## Supporting Information

Additional material, including methods, comprehensive sets of data and additional analyses.

## Abbreviations

3D: three-dimensional
CHL1: cholesterol
CF: Cystic Fibrosis
CFTR: Cystic Fibrosis Transmembrane Conductance Regulator
cryo-EM: cryo-electron microscopy
HB: Hydrogen Bond
SB: Salt Bridge
ECL: ExtraCellular Loop
TMD: TransMembrane Domain
ICL: IntraCellular Loop
TM: TransMembrane
NBD: Nucleotide Binding Domain
MD: Molecular Dynamics
DSM: sphingolipid
POPC: 1-palmitoyl-2-oleoylphosphatidylcholine)
POPE: 1-palmitoyl-2-oleoylphosphatidylethanolamine)
POPS: 1-palmitoyl-2-oleoylphosphatidylserine

