## Supplementary data for "Mechanistic insights into CFTR function from molecular dynamics analysis of electrostatic interactions"

### Brief description / List of the contents

#### Supplementary Figures

**Figure S1.** RMSD of human CFTR 3D structures during MD simulations under apo and VX-770-bound conditions.

**Figure S2.** Membrane structural properties during MD simulations of the human CFTR 3D structure under apo and VX-770-bound conditions.

**Figure S3:** Contacts between basic/acidic side-chains along MD simulations of the human CFTR 3D structure in apo and VX-770-bound conditions.

**Figure S4:** Other contacts along MD simulations of the human CFTR 3D structure in apo and VX-770-bound conditions.

**Figure S5.** Charged/polar residues–anion interactions ($Cl^-$ and $HCO_3^-$) along MD simulations of the human CFTR 3D structure in apo and VX-770-bound conditions.

**Figure S6.** Charged/polar residues–lipid headgroup contacts along MD simulations of the human CFTR 3D structure in apo and VX-770-bound conditions.

**Figure S7.** Distribution of contacts between amino acids in the CFTR 3D structure – MD simulations in apo systems (4 replica).

**Figure S8.** Mapping of side-chain/side-chain contacts also involved in anion and/or lipid interactions in the CFTR 3D structure– MD simulations in apo systems (4 replica).

**Figure S9.** Detailed views of contacts in the different regions of the human CFTR 3D structure) – MD simulations in apo systems.

**Figure S10.** Residues involved in anion interactions and their overlap with inter-residue and lipid contacts – MD simulations in apo systems (4 replica).

**Figure S11.** Most frequent residues involved in lipid headgroups interactions and their overlap with inter-residue and anion contacts – MD simulations in apo systems (4 replica).

**Figure S12**. Comparison of the frequencies of contacts between the Apo and VX-770-bound conditions.

**Figure S13:** Residues involved in multiple interaction types.

#### Supplementary Tables

**Table S1.** MD Simulations of human CFTR.

**Table S2.** Amino acids involved in TM α-helix irregularities and involvement of charged/polar side chains in side-chain/side-chain contacts.

**Table S3.** Per-residue changes in amino acids/anion/lipid contacts in MD simulations in the presence of VX-770.

**Table S4.** Portals and exits observed at the end of the MD simulations.

**Table S5.** Amino acid couples with side chain/side chain contacts and homologous positions in human ABCC4.

#### Supplementary Figures

**Figure S1. RMSD of human CFTR 3D structures during MD simulations under apo and VX-770-bound conditions.**
The plot displays the time evolution of root-mean-square deviation (RMSD) for eight independent molecular dynamics simulations: four apo replicates (Apo1 (dark red), Apo2 (red), Apo3 (orange), Apo4 (purple)) and four VX-770-bound replicates (VX1 (blue), VX2 (cyan), VX3 (aquamarine) and VX4 (azure)). RMSD was calculated relative to the initial cryo-EM structure after alignment on the Cα atoms.

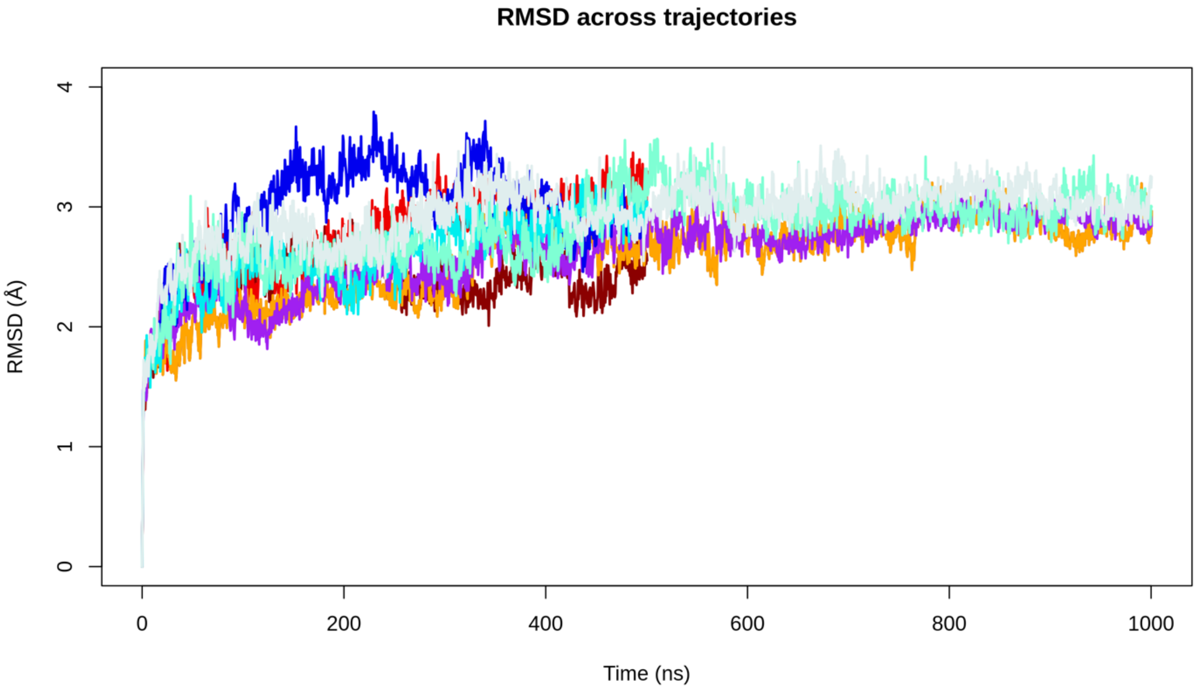

**Figure S2. Membrane structural properties during MD simulations of the human CFTR 3D structure under apo and VX-770-bound conditions.**
Top: Area per lipid (Å²) calculated over time for the eight simulation replicates (Apo1–Apo4, VX1–VX4). Bottom: Membrane thickness (Å) evaluated between the phosphate headgroup planes of the upper and lower leaflets. Each trace corresponds to a separate MD replica: Apo1 (dark red), Apo2 (red), Apo3 (orange), Apo4 (purple), VX1 (blue), VX2 (cyan), VX3 (aquamarine) and VX4 (azure).

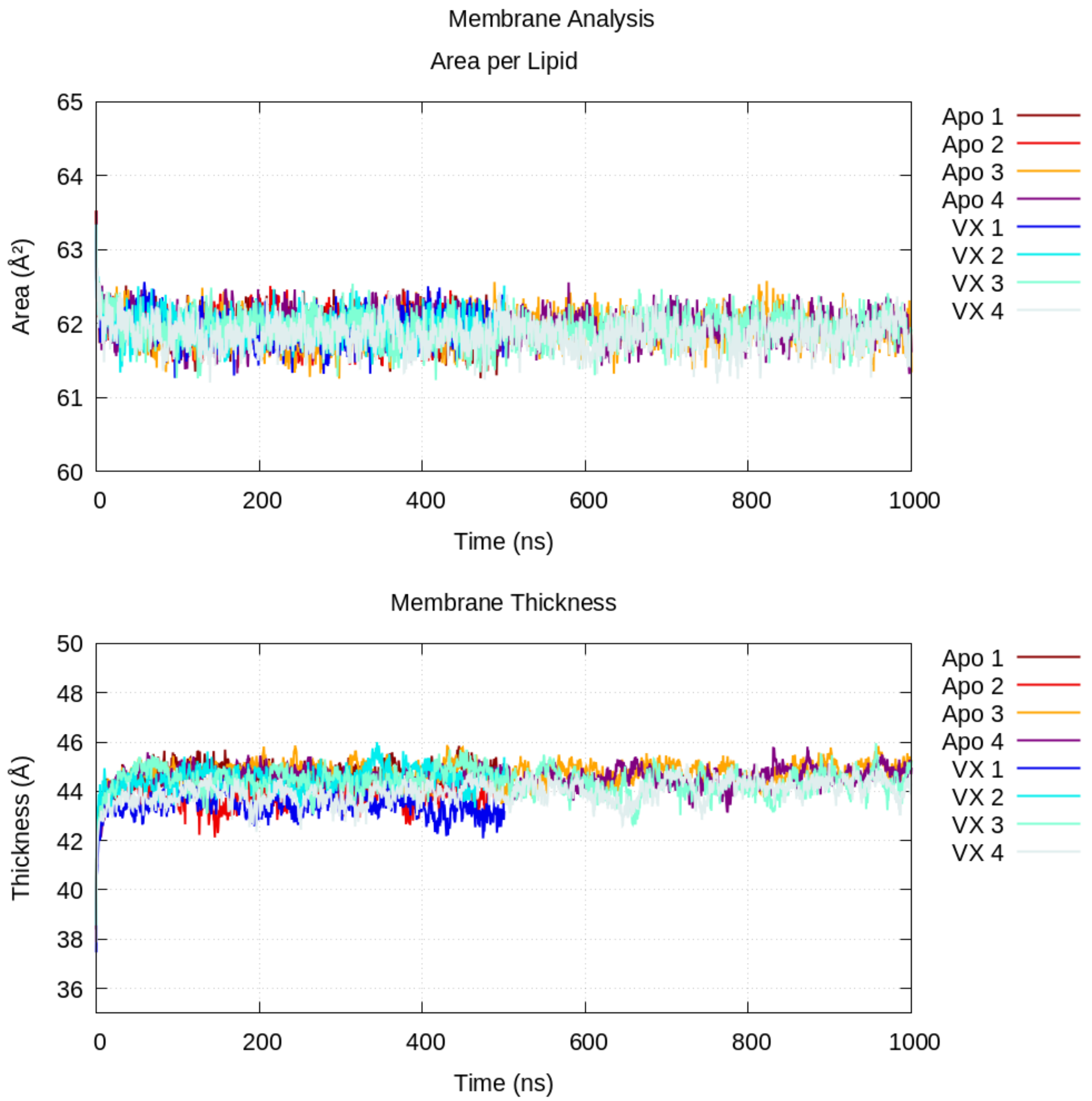

**Figure S3: Contacts between basic/acidic side-chains along MD simulations of the human CFTR 3D structure in apo and VX-770-bound conditions.**
Violin plots represent the distribution of contact frequencies (and their means) observed during for the 4 MD simulation in the Apo (orange) and VX-770-bound (green) conditions. Contact are ordered according to the numbering in the sequence of the first residue of the pair. This representation highlights the persistence and variability of contacts between ligand-free and ligand-bound conditions.

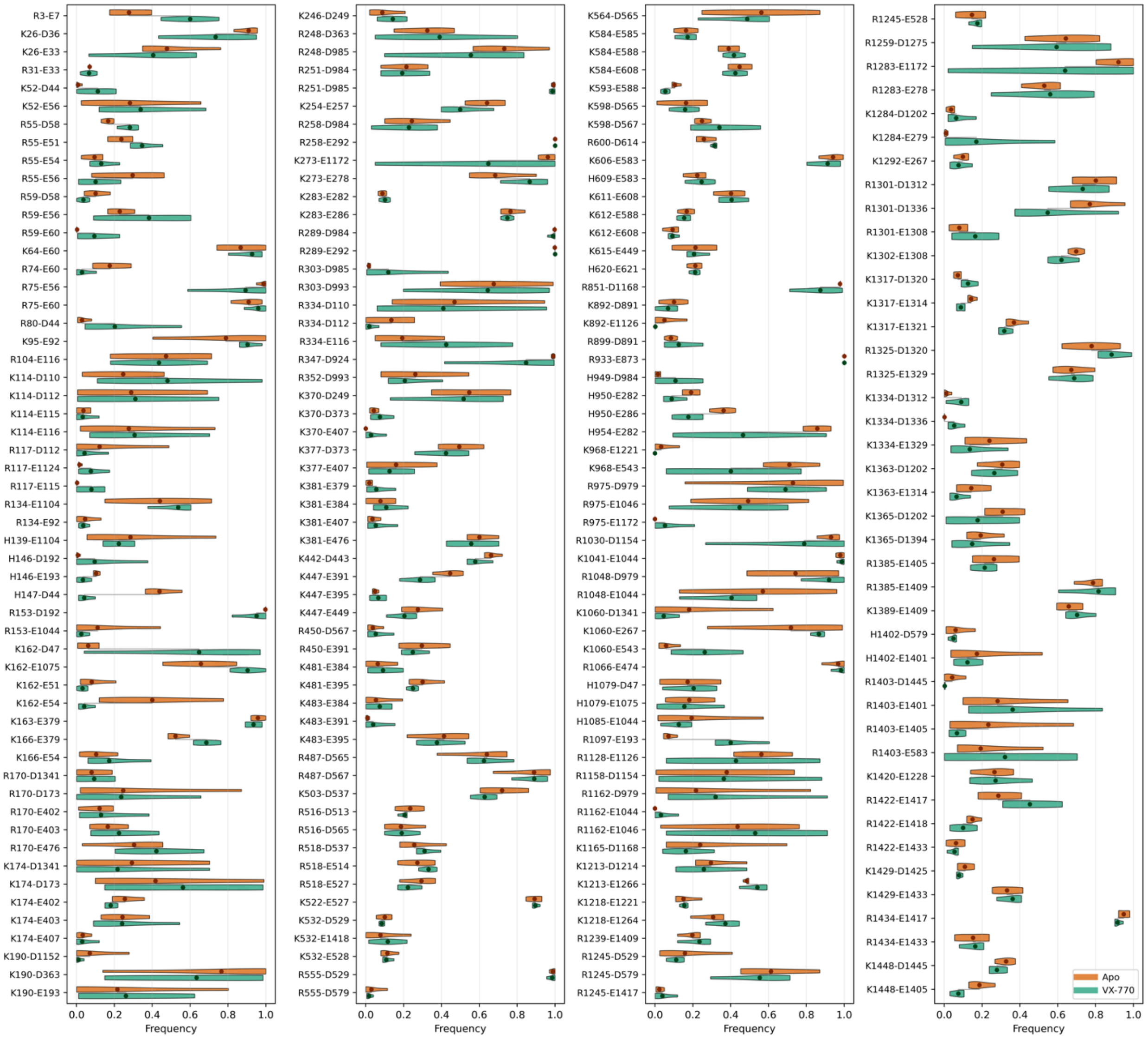

**Figure S4: Other contacts along MD simulations of the human CFTR 3D structure in apo and VX-770-bound conditions.** Violin plots represent the distribution of contact frequencies (and their means) observed during for the 4 MD simulation in the Apo (orange) and VX-770-bound (green) conditions, outside of the pairs shown in Figure S3. contacts are ordered according to the numbering in the sequence of the first residue of the pair. This representation highlights the persistence and variability of contacts between ligand-free and ligand-bound conditions. The three panels report the interactions of **(A)** S/T with either R/K/H or D/E, **(B)** N/Q/W/Y/C with R/K/H/D/E, (**C)** N/Q/W/Y/C with N/Q/W/Y/C/S/T.
Black stars indicated cation-π interactions between tryptophan (W) and arginine (R) residues, whereas the red star highlights the contact between H199 and W202.

**A) Interaction of S/T with either R/K/H, D/E or S/T**

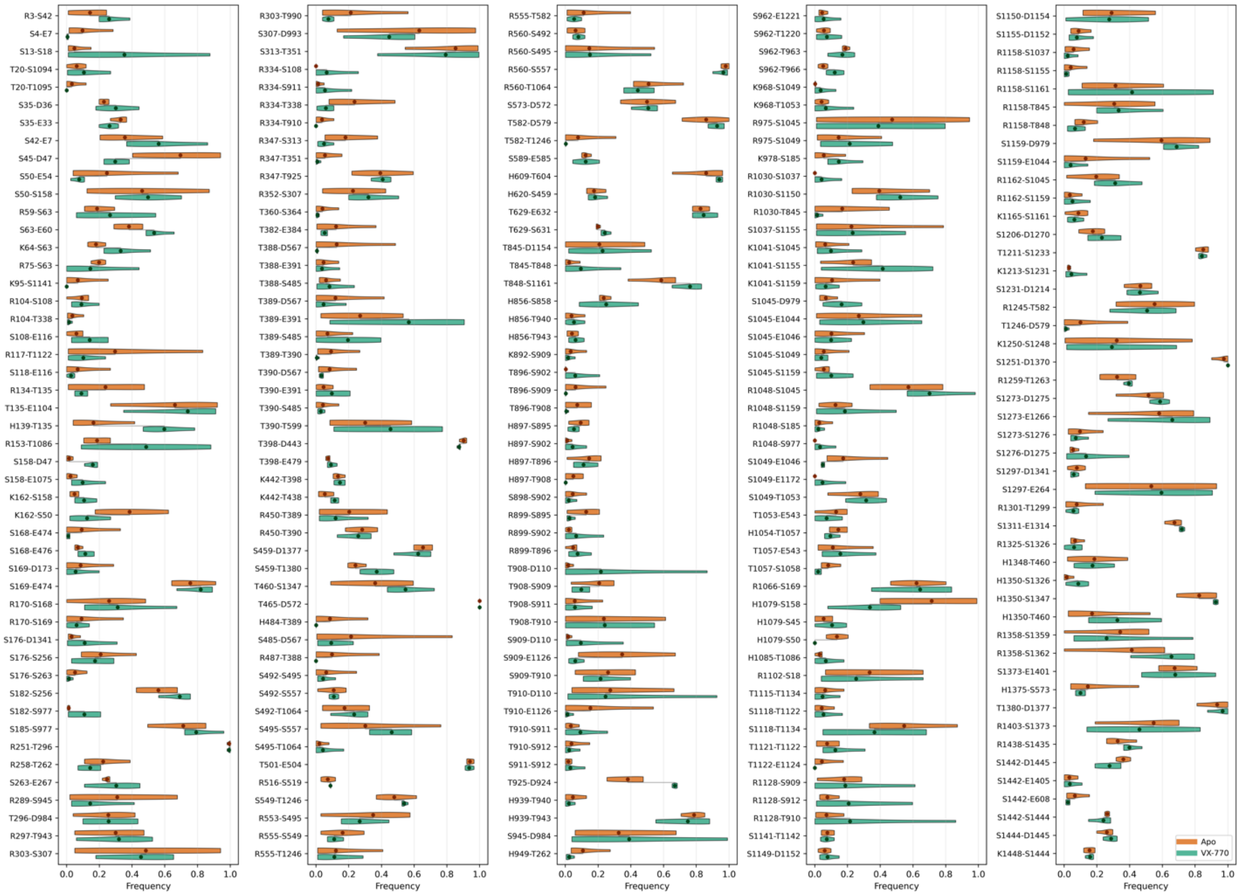

#### B) Interaction of N/Q/W/Y/C with R/K/H/D/E

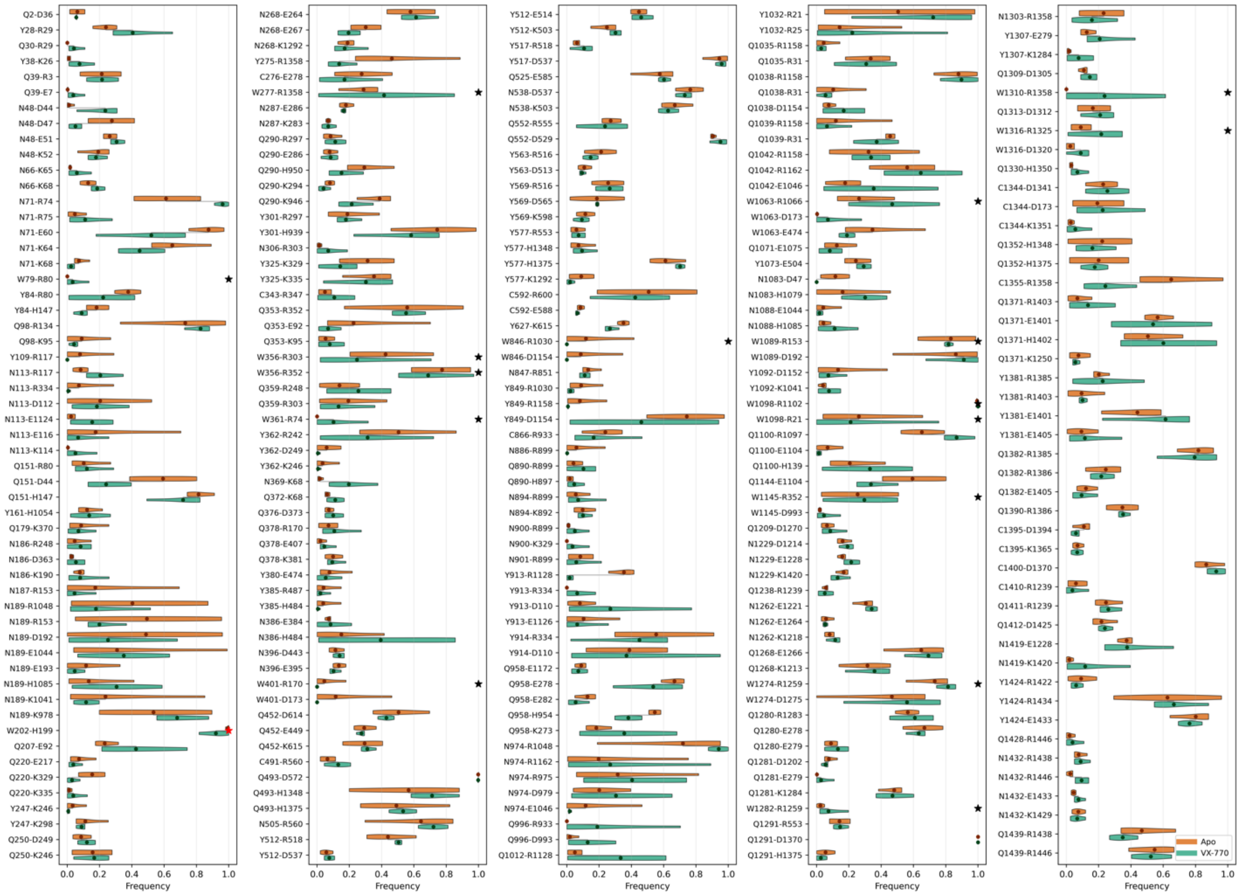

#### C) Interaction of N/Q/W/Y/C with N/Q/W/Y/C/S/T.

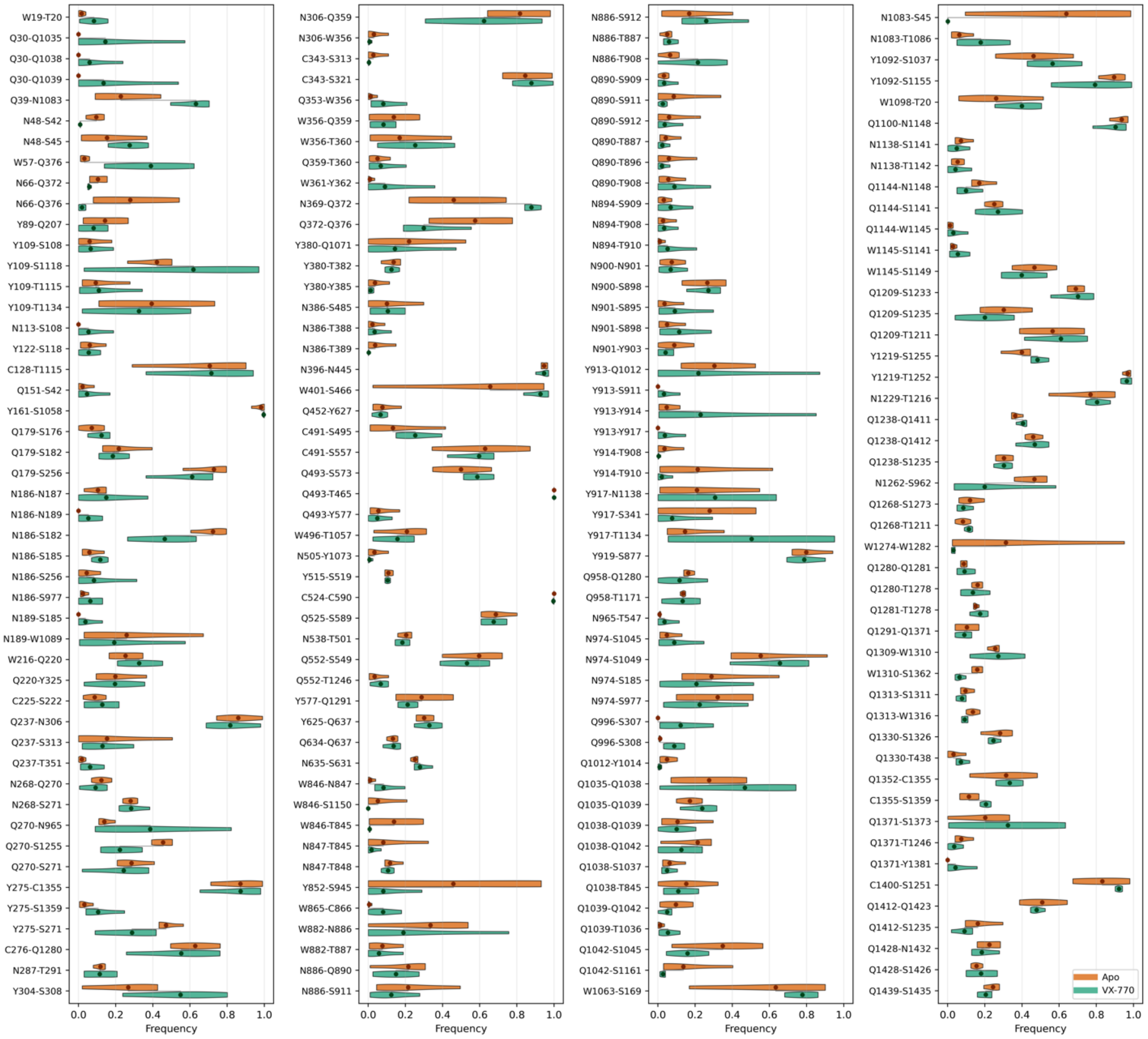

**Figure S5. Charged/polar residues–anion interactions ($Cl^-$ and $HCO_3^-$) along MD simulations of the human CFTR 3D structure in apo and VX-770-bound conditions.** Violin plots represent the distribution of contact frequencies (and their means) observed during for the 4 MD simulation in the Apo (orange) and VX-770-bound (green) conditions. For each residue–anion combinations is reported the fraction of simulation time during which the interaction was present. The two panels report the interactions involving **(A)** R/K/H/S/T, **(B)** N/Q/Y/W/C.

#### A) Interaction with R/K/H/S/T

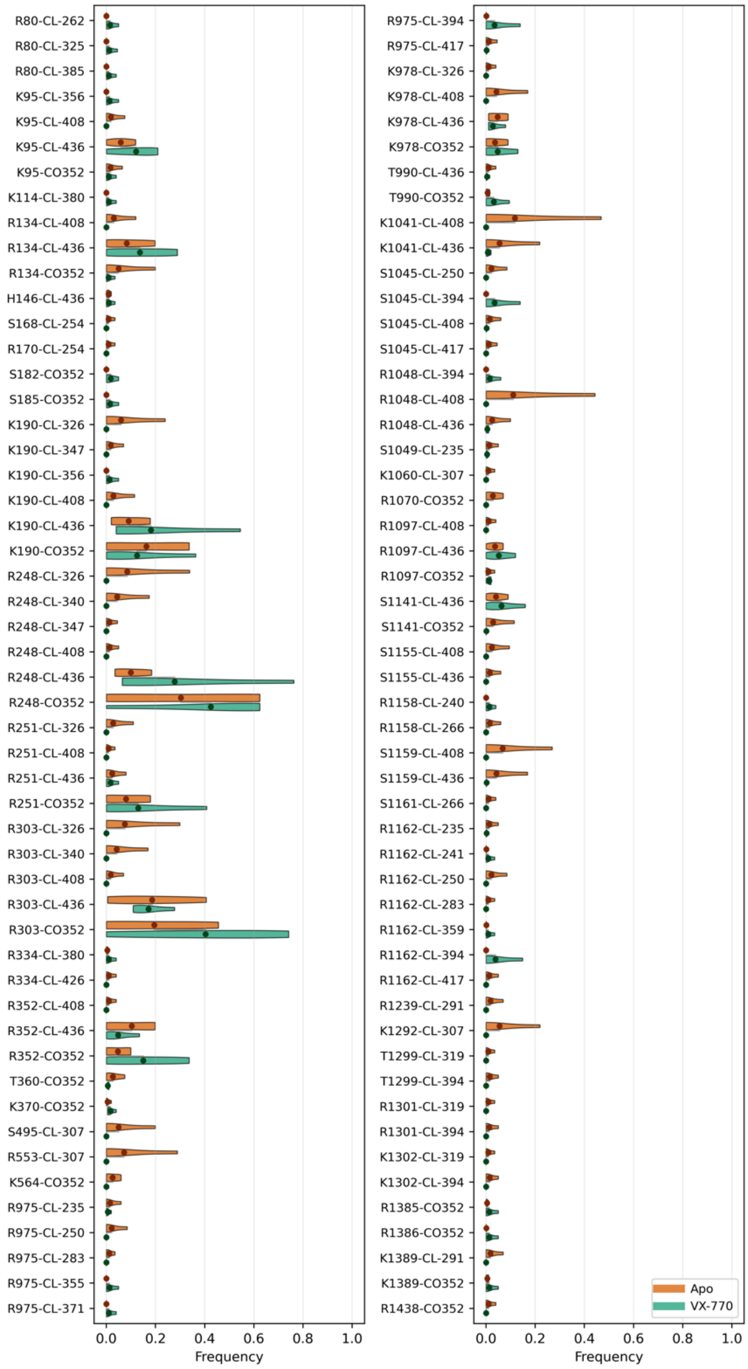

#### B) Interaction with N/Q/Y/W/C

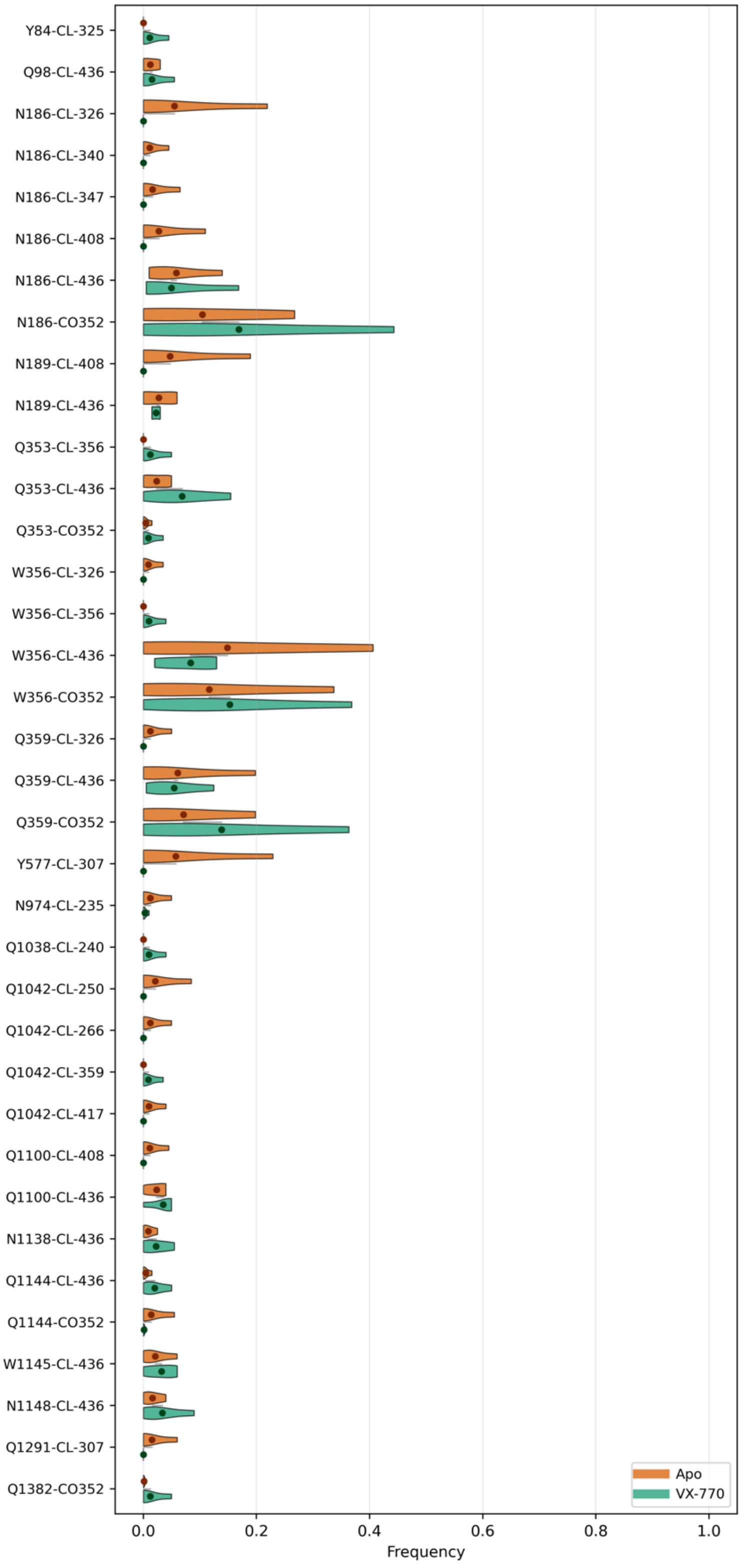

**Figure S6. Charged/polar residues–lipid headgroup contacts along MD simulations of the human CFTR 3D structure in apo and VX-770-bound conditions.** Violin plots represent the distribution of contact frequencies (and their means) observed during for the 4 MD simulation in the Apo (orange) and VX-770-bound (green) conditions. For each residue–lipid pair is reported the fraction of simulation time during which the interaction was present. The two panels report the interactions involving **(A)** R/K/H/S/T, **(B)** N/Q/Y/W/C. POC, POS, POE stand for POPC, POPS and POPE, respectively.

**(A) Interaction with R/K/H/S/T (1/2)**

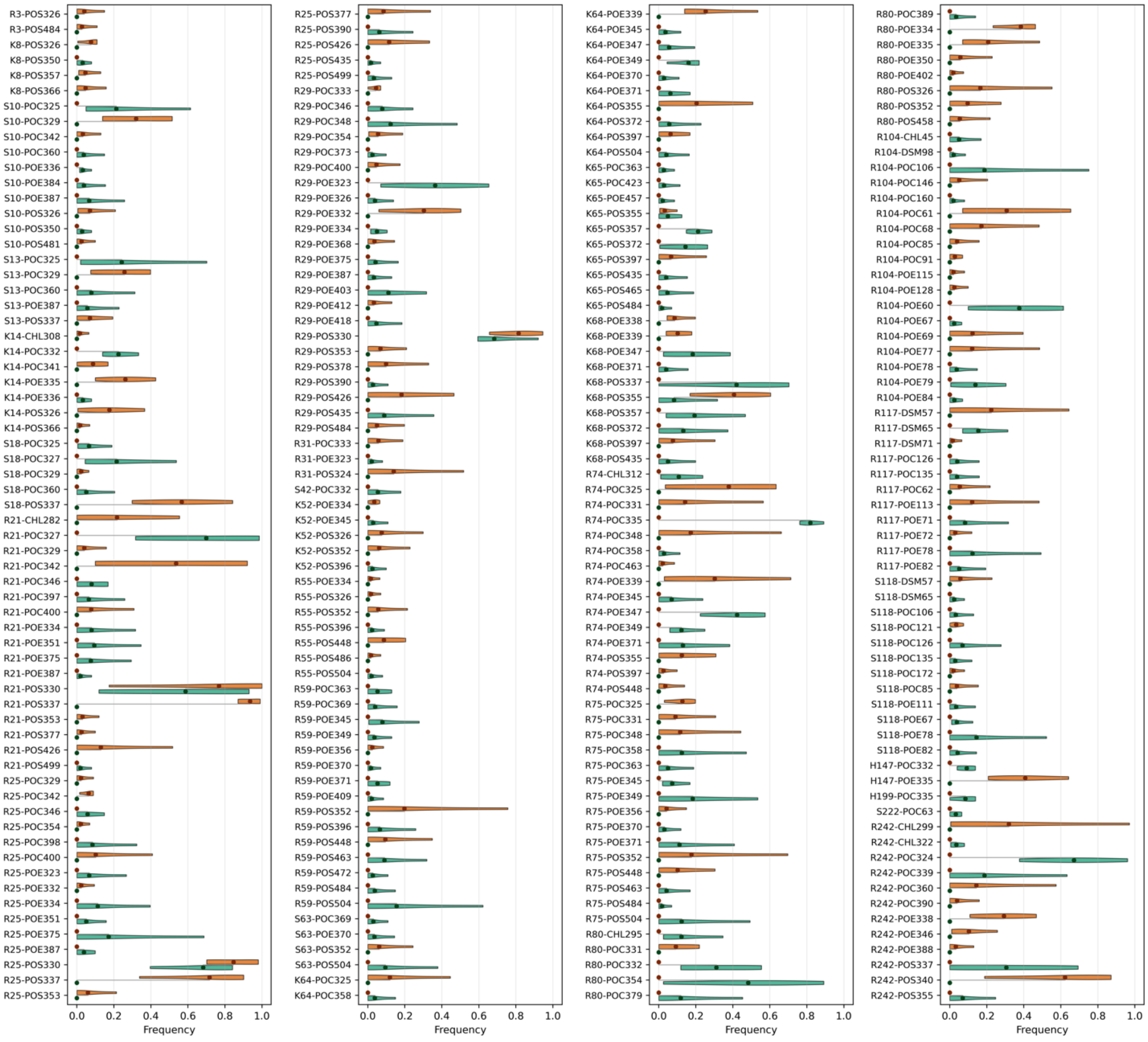

#### Interaction with R/K/H/S/T (2/2)

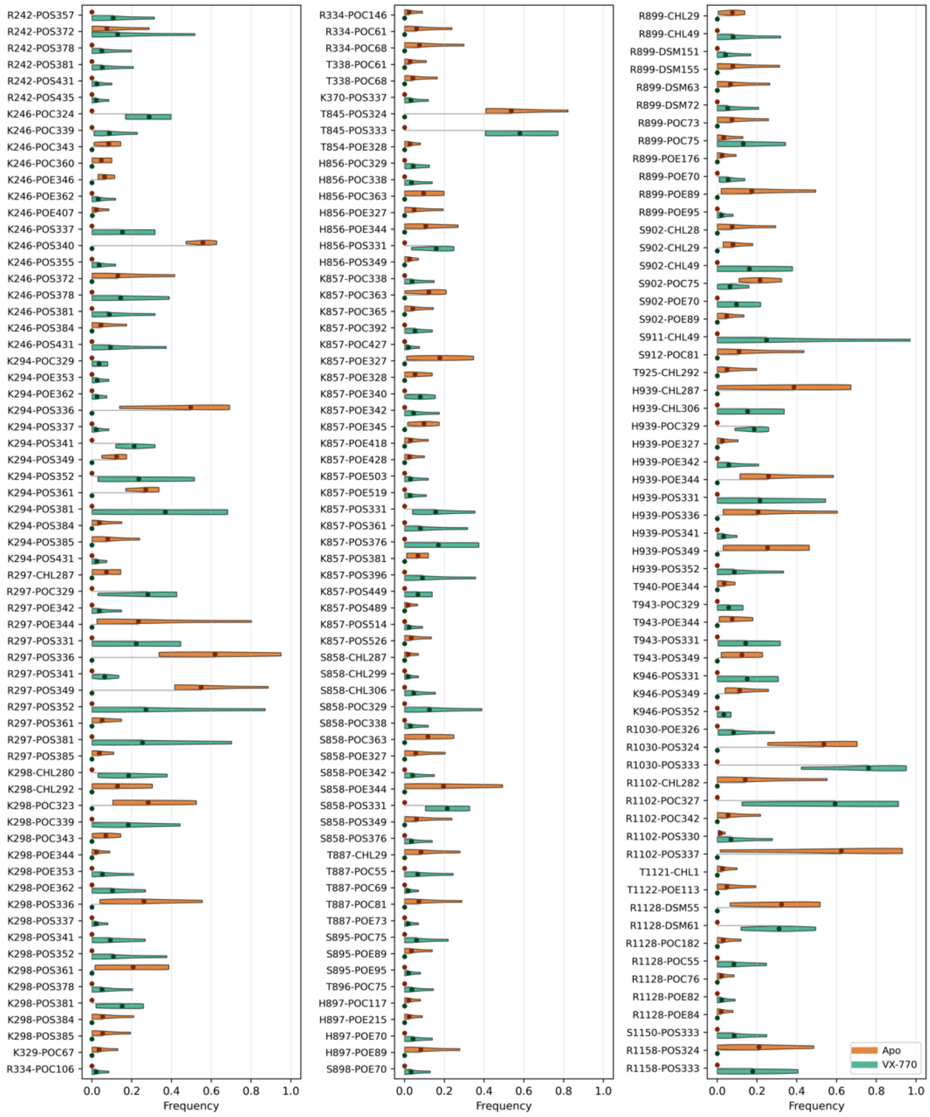

C) Interaction with N/Q/Y/W/C

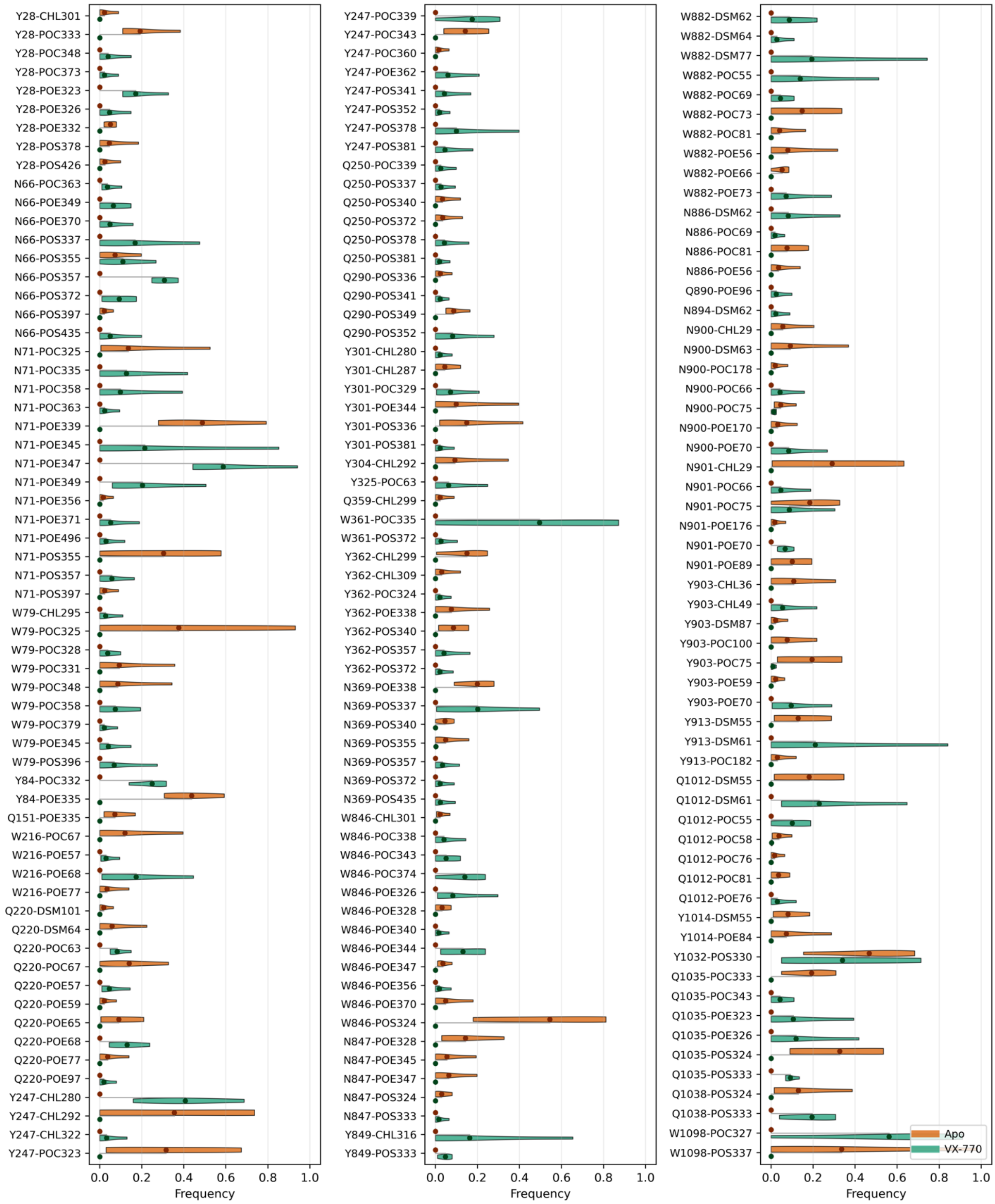

**Figure S7. Distribution of contacts between amino acids in the CFTR 3D structure – MD simulations in apo systems (4 replica).** Violin plots showing the distribution of contact frequencies in the 4 replica (Apo conditions) for each amino acid pairs, presented in the descending order of frequency means (bars). Only contacts with a mean frequency ≥ 0.10 across all systems are shown. These interactions were assigned to five regions – **(A)** ECLs, **(B)** TM helices, **(C)** Elbow/Lasso/ICLs, **(D)** ICLs/NBDs and **(E)** NBDs - according to the average Z-coordinate of the involved residues Cα atoms. For each region, the four plots refer, from left to right, to contacts between side chain charged/polar atoms from R/K/H/D/E/S/T (1) and from N/Q/Y/W/C with other charged or polar atoms (2), and contacts between side-chain charged/polar atoms (amino acids at left) from R/K/H/D/E/S/T (1) and from N/Q/Y/W/C (2) and main chain atoms (amino acids at right). Black stars indicated cation-∏ interactions between tryptophan (W) and arginine (R) residues, whereas the red star highlights the contact between H199 and W202.

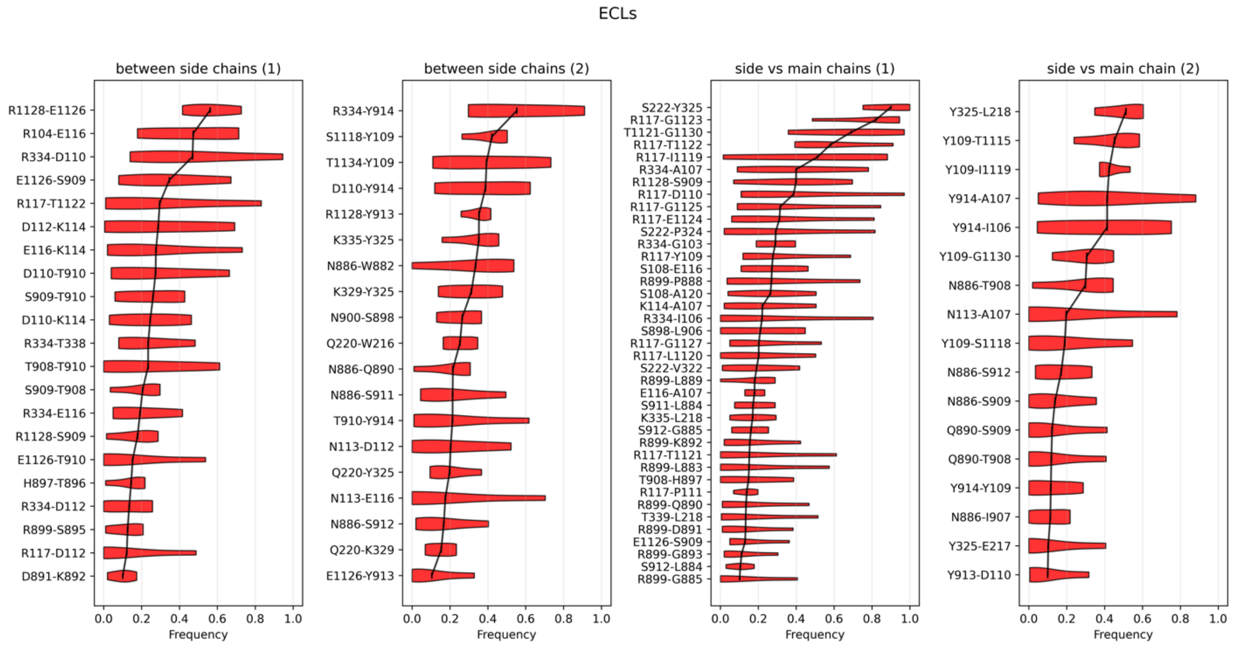

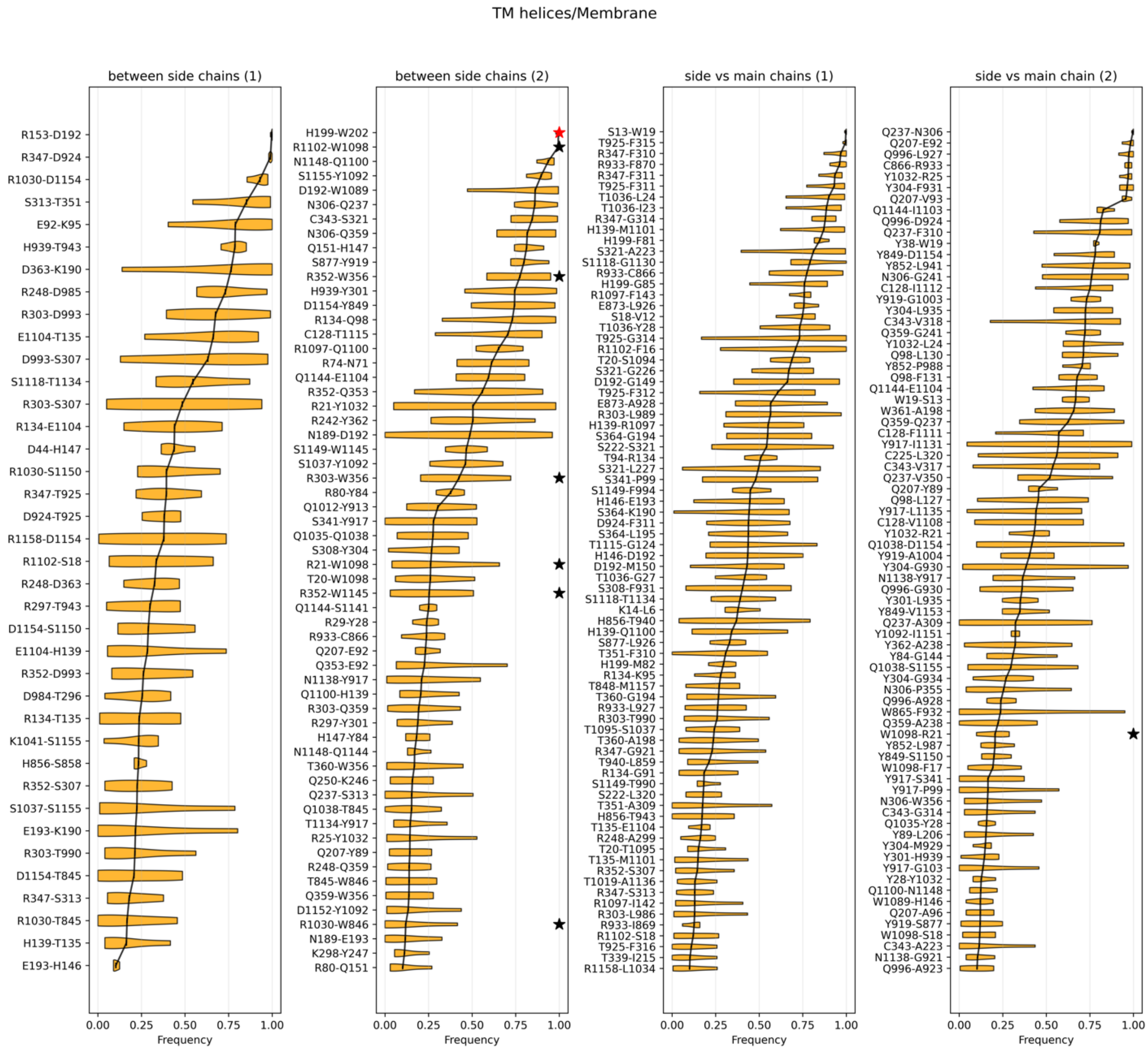
TM helices/Membrane
between side chains (1)
R153-D192
R347-D924
R1030-D1154
S313-T351
E92-K95
H939-T943
D363-K190
R248-D985
R303-D993
E1104-T135
D993-S307
S1118-T1134
R303-S307
R134-E1104
D44-H147
R1030-S1150
R347-T925
D924-T925
R1158-D1154
R1102-S18
R248-D363
R297-T943
D1154-S1150
E1104-H139
R352-D993
D984-T296
R134-T135
K1041-S1155
H856-S858
R352-S307
S1037-S1155
E193-K190
R303-T990
D1154-T845
R347-S313
R1030-T845
H139-T135
E193-H146
between side chains (2)
H199-W202
R1102-W1098
N1148-Q1100
S1155-Y1092
D192-W1089
N306-Q237
C343-S321
N306-Q359
Q151-H147
S877-Y919
R352-W356
H939-Y301
D1154-Y849
R134-Q98
C128-T1115
R1097-Q1100
R74-N71
Q1144-E1104
R352-Q353
R21-Y1032
R242-Y362
N189-D192
S1149-W1145
S1037-Y1092
R303-W356
R80-Y84
Q1012-Y913
S341-Y917
Q1035-Q1038
S308-Y304
R21-W1098
T20-W1098
R352-W1145
Q1144-S1141
R29-Y28
R933-C866
Q207-E92
Q353-E92
N1138-Y917
Q1100-H139
R303-Q359
R297-Y301
H147-Y84
N1148-Q1144
T360-W356
Q250-K246
Q237-S313
Q1038-T845
T1134-Y917
R25-Y1032
Q207-Y89
R248-Q359
T845-W846
Q359-W356
D1152-Y1092
R1030-W846
N189-E193
K298-Y247
R80-Q151
side vs main chains (1)
S13-W19
T925-F315
R347-F310
R933-F870
R347-F311
T925-F311
T1036-L24
T1036-I23
R347-G314
H139-M1101
H199-F81
S321-A223
S1118-G1130
R933-C866
H199-G85
R1097-F143
E873-L926
S18-V12
T1036-Y28
T925-G314
R1102-F16
T20-S1094
S321-G226
D192-G149
T925-F312
E873-A928
R303-L989
H139-R1097
S364-G194
S222-S321
T94-R134
S321-L227
S341-P99
S1149-F994
H146-E193
S364-K190
D924-F311
S364-L195
T1115-G124
H146-D192
D192-M150
T1036-G27
S308-F931
S1118-T1134
K14-L6
H856-T940
H139-Q1100
S877-L926
T351-F310
H199-M82
R134-K95
T848-M1157
T360-G194
R933-L927
R303-T990
T1095-S1037
T360-A198
R347-G921
T940-L859
R134-G91
S1149-T990
S222-L320
T351-A309
H856-T943
T135-E1104
R248-A299
T20-T1095
T135-M1101
R352-S307
T1019-A1136
R347-S313
R1097-I142
R303-L986
R933-I869
R1102-S18
T925-F316
T339-I215
R1158-L1034
side vs main chain (2)
Q237-N306
Q207-E92
Q996-L927
C866-R933
Y1032-R25
Y304-F931
Q207-V93
Q1144-I1103
Q996-D924
Q237-F310
Y38-W19
Y849-D1154
Y852-L941
N306-G241
C128-I1112
Y919-G1003
Y304-L935
C343-V318
Q359-G241
Y1032-L24
Q98-L130
Y852-P988
Q98-F131
Q1144-E1104
W19-S13
W361-A198
Q359-Q237
C128-F1111
Y917-I1131
C225-L320
C343-V317
Q237-V350
Q207-Y89
Q98-L127
Y917-L1135
C128-V1108
Y1032-R21
Q1038-D1154
Y919-A1004
Y304-G930
N1138-Y917
Q996-G930
Y301-L935
Y849-V1153
Q237-A309
Y1092-I1151
Y362-A238
Y84-G144
Q1038-S1155
Y304-G934
N306-P355
Q996-A928
W865-F932
Q359-A238
W1098-R21
Y852-L987
Y849-S1150
W1098-F17
Y917-S341
Y917-P99
N306-W356
C343-G314
Q1035-Y28
Y89-L206
Y304-M929
Y301-H939
Y917-G103
Y28-Y1032
Q1100-N1148
W1089-H146
Q207-A96
Y919-S877
W1098-S18
C343-A223
N1138-G921
Q996-A923
0.00
0.25
0.50
0.75
1.00
Frequency

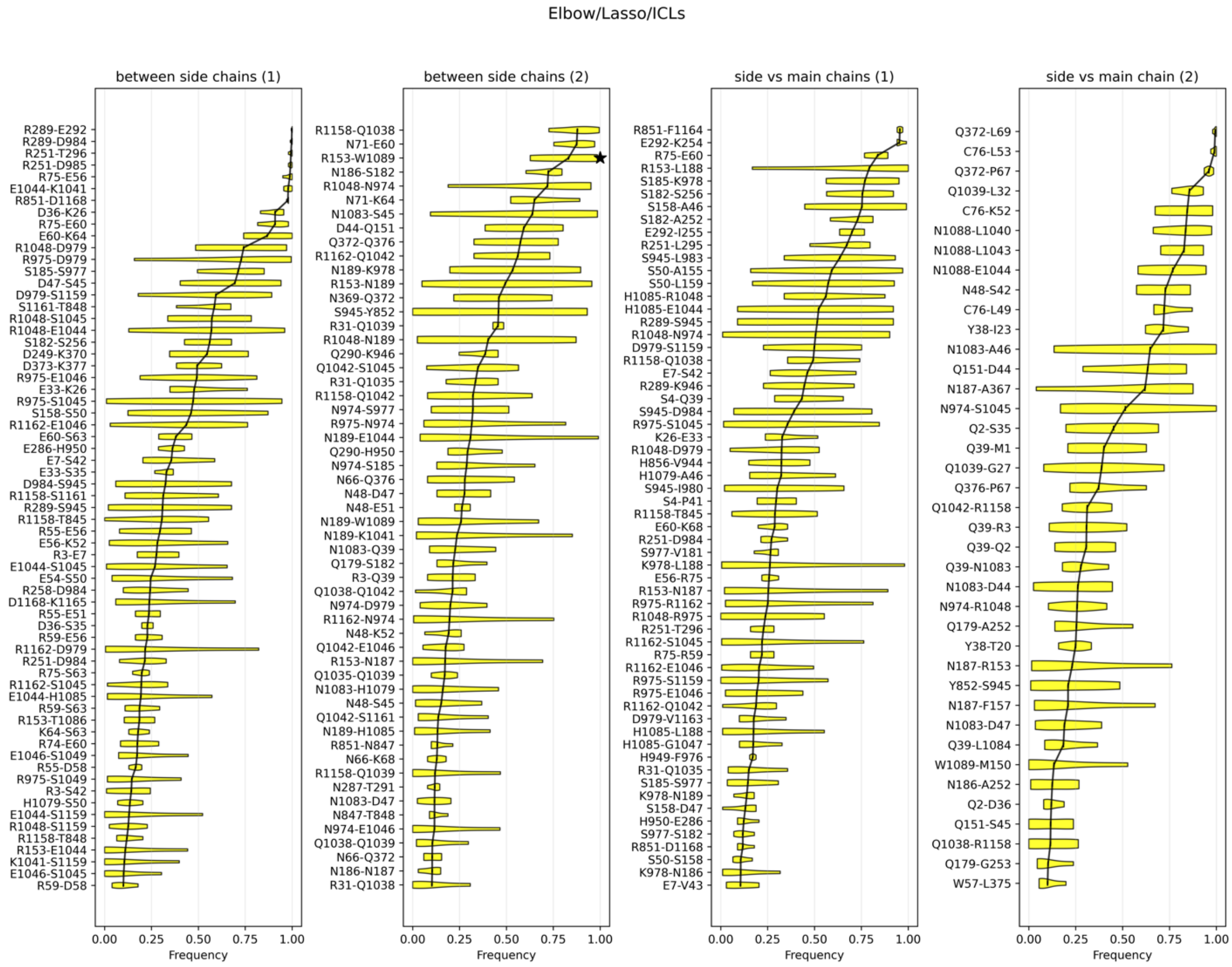
Elbow/Lasso/ICLs
between side chains (1)
R289-E292
R289-D984
R251-T296
R251-D985
R75-E56
E1044-K1041
R851-D1168
D36-K26
R75-E60
E60-K64
R1048-D979
R975-D979
S185-S977
D47-S45
D979-S1159
S1161-T848
R1048-S1045
R1048-E1044
S182-S256
D249-K370
D373-K377
R975-E1046
E33-K26
R975-S1045
S158-S50
R1162-E1046
E60-S63
E286-H950
E7-S42
E33-S35
D984-S945
R1158-S1161
R289-S945
R1158-T845
R55-E56
E56-K52
R3-E7
E1044-S1045
E54-S50
R258-D984
D1168-K1165
R55-E51
D36-S35
R59-E56
R1162-D979
R251-D984
R75-S63
R1162-S1045
E1044-H1085
R59-S63
R153-T1086
K64-S63
R74-E60
E1046-S1049
R55-D58
R975-S1049
R3-S42
H1079-S50
E1044-S1159
R1048-S1159
R1158-T848
R153-E1044
K1041-S1159
E1046-S1045
R59-D58
between side chains (2)
R1158-Q1038
N71-E60
R153-W1089
N186-S182
R1048-N974
N71-K64
N1083-S45
D44-Q151
Q372-Q376
R1162-Q1042
N189-K978
R153-N189
N369-Q372
S945-Y852
R31-Q1039
R1048-N189
Q290-K946
Q1042-S1045
R31-Q1035
R1158-Q1042
N974-S977
R975-N974
N189-E1044
Q290-H950
N974-S185
N66-Q376
N48-D47
N48-E51
N189-W1089
N189-K1041
N1083-Q39
Q179-S182
R3-Q39
Q1038-Q1042
N974-D979
R1162-N974
N48-K52
Q1042-E1046
R153-N187
Q1035-Q1039
N1083-H1079
N48-S45
Q1042-S1161
N189-H1085
R851-N847
N66-K68
R1158-Q1039
N287-T291
N1083-D47
N847-T848
N974-E1046
Q1038-Q1039
N66-Q372
N186-N187
R31-Q1038
side vs main chains (1)
R851-F1164
E292-K254
R75-E60
R153-L188
S185-K978
S182-S256
S158-A46
S182-A252
E292-I255
R251-L295
S945-L983
S50-A155
S50-L159
H1085-R1048
H1085-E1044
R289-S945
R1048-N974
D979-S1159
R1158-Q1038
E7-S42
R289-K946
S4-Q39
S945-D984
R975-S1045
K26-E33
R1048-D979
H856-V944
H1079-A46
S945-I980
S4-P41
R1158-T845
E60-K68
R251-D984
S977-V181
K978-L188
E56-R75
R153-N187
R975-R1162
R1048-R975
R251-T296
R1162-S1045
R75-R59
R1162-E1046
R975-S1159
R975-E1046
R1162-Q1042
D979-V1163
H1085-L188
H1085-G1047
H949-F976
R31-Q1035
S185-S977
K978-N189
S158-D47
H950-E286
S977-S182
R851-D1168
S50-S158
K978-N186
E7-V43
side vs main chain (2)
Q372-L69
C76-L53
Q372-P67
Q1039-L32
C76-K52
N1088-L1040
N1088-L1043
N1088-E1044
N48-S42
C76-L49
Y38-I23
N1083-A46
Q151-D44
N187-A367
N974-S1045
Q2-S35
Q39-M1
Q1039-G27
Q376-P67
Q1042-R1158
Q39-R3
Q39-Q2
Q39-N1083
N1083-D44
N974-R1048
Q179-A252
Y38-T20
N187-R153
Y852-S945
N187-F157
N1083-D47
Q39-L1084
W1089-M150
N186-A252
Q2-D36
Q151-S45
Q1038-R1158
Q179-G253
W57-L375
0.00
0.25
0.50
0.75
1.00
Frequency

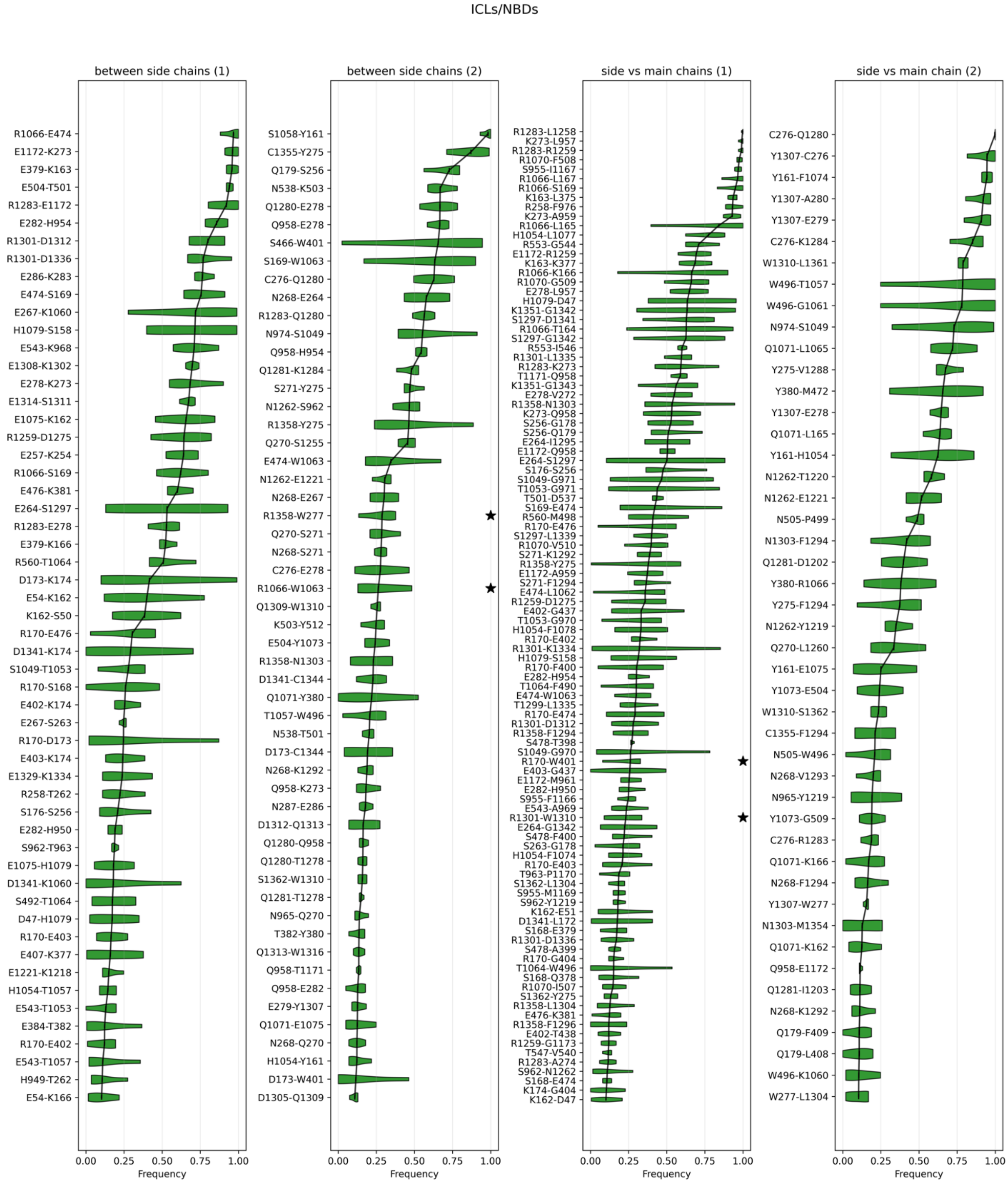
ICLs/NBDs
between side chains (1)
between side chains (2)
side vs main chains (1)
side vs main chain (2)
Frequency

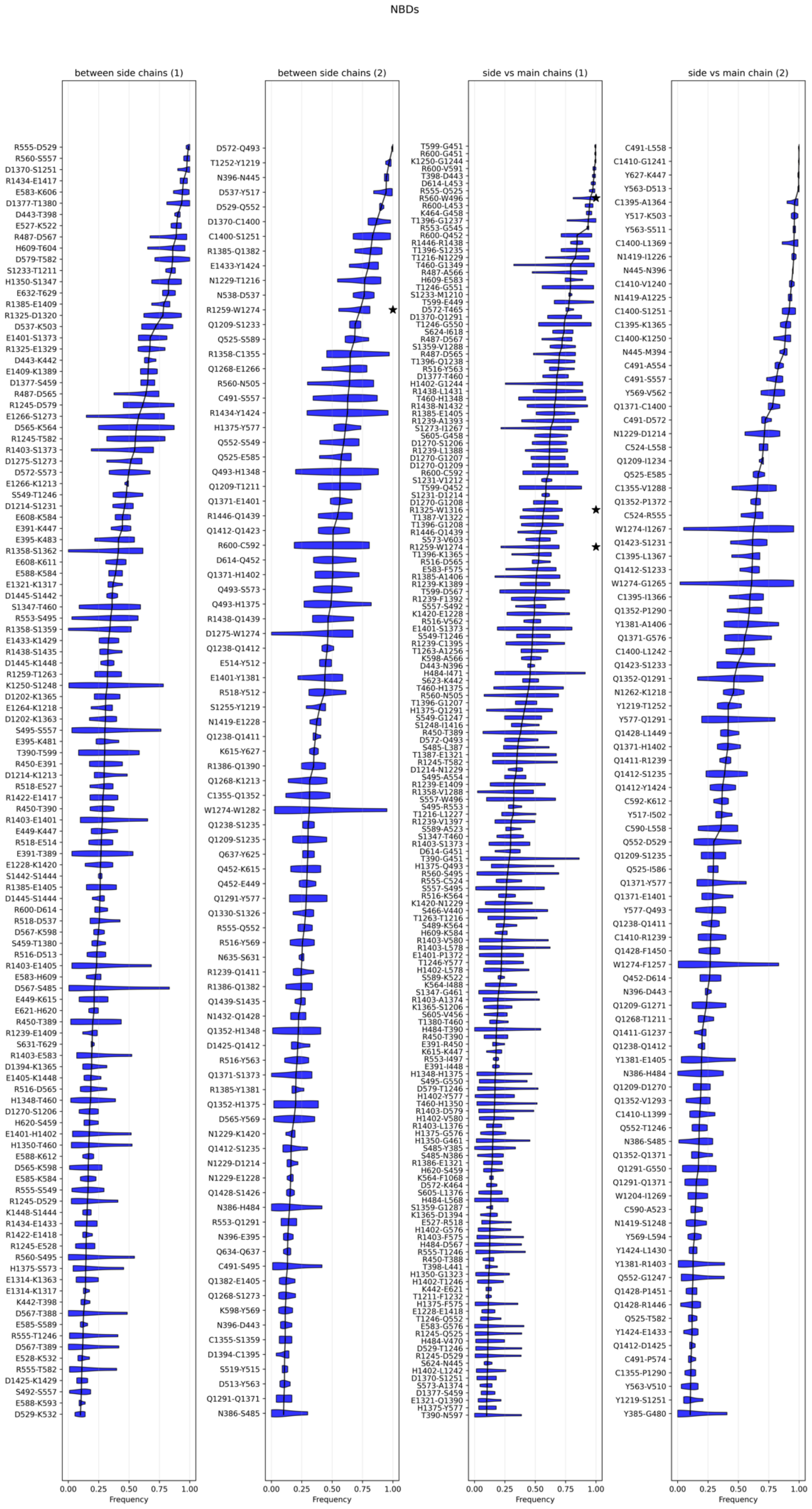
NBDs
between side chains (1)
between side chains (2)
side vs main chains (1)
side vs main chain (2)
Frequency
0.00 0.25 0.50 0.75 1.00

**Figure S8. Mapping of side-chain/side-chain contacts also involved in anion and/or lipid interactions in the CFTR 3D structure – MD simulations in apo systems (4 replica).**–Violin plots showing the distribution of side-chain/side-chain contact frequencies in the 4 replica (Apo conditions) for each amino acid pairs, presented in the descending order of frequency means (bars). Electrostatic interactions with a mean frequency ≥ 0.10 across MD simulations were categorized by regions in the CFTR 3D structure based on the average Z-coordinate of the involved Cα atoms: ECLs, TM helices, Elbow/Lasso/ICLs, ICLs/NBDs and NBDs. Distribut are color-coded based on whether the side chain of an amino acid involved in electrostatic interactions with another amino acid side chain also establishes contacts with anions (cyan), lipids (yellow), both (purple), or neither (light grey). This analysis highlights electrostatic networks potentially coupled to membrane interactions or/and ion coordination, suggesting region-specific regulatory or structural roles. The two panels refer to interactions involving (1) R/K/H/D/S/T, (2) E/N/Q/Y/W/C. Black stars indicated cation-∏ interactions between tryptophan (W) and arginine (R) residues, whereas the red star highlights the contact between H199 and W202.

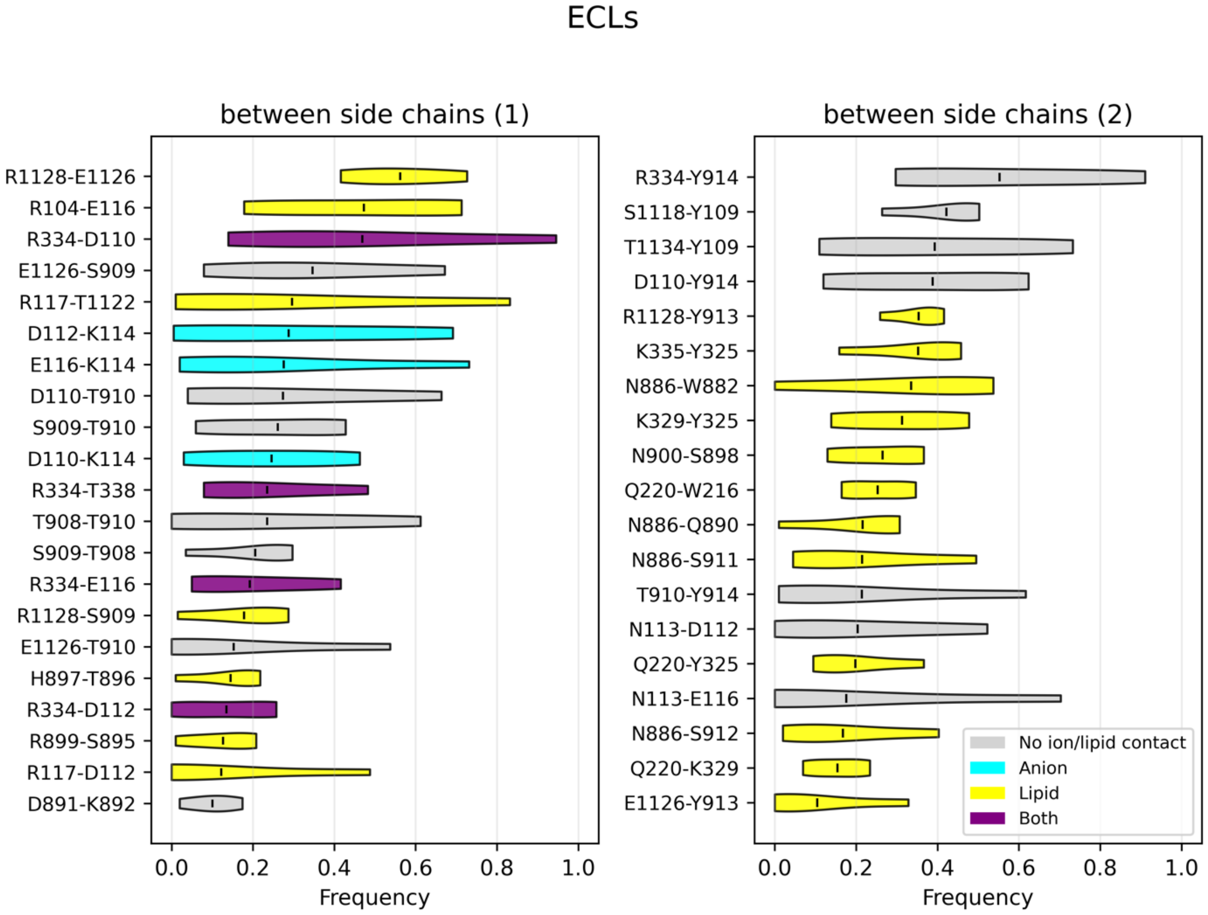

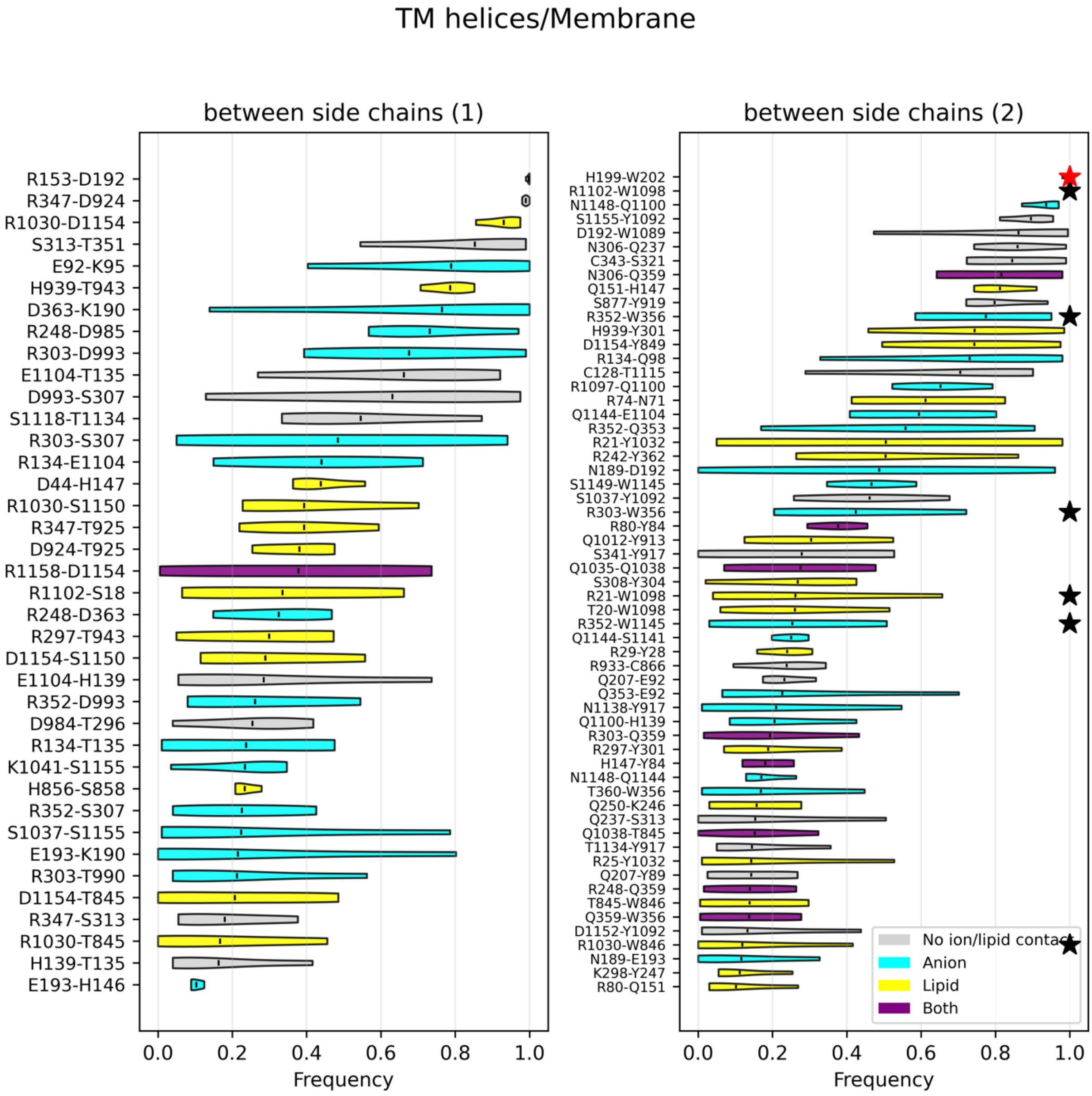
TM helices/Membrane
between side chains (1)
R153-D192
R347-D924
R1030-D1154
S313-T351
E92-K95
H939-T943
D363-K190
R248-D985
R303-D993
E1104-T135
D993-S307
S1118-T1134
R303-S307
R134-E1104
D44-H147
R1030-S1150
R347-T925
D924-T925
R1158-D1154
R1102-S18
R248-D363
R297-T943
D1154-S1150
E1104-H139
R352-D993
D984-T296
R134-T135
K1041-S1155
H856-S858
R352-S307
S1037-S1155
E193-K190
R303-T990
D1154-T845
R347-S313
R1030-T845
H139-T135
E193-H146
0.0
0.2
0.4
0.6
0.8
1.0
Frequency
between side chains (2)
H199-W202
R1102-W1098
N1148-Q1100
S1155-Y1092
D192-W1089
N306-Q237
C343-S321
N306-Q359
Q151-H147
S877-Y919
R352-W356
H939-Y301
D1154-Y849
R134-Q98
C128-T1115
R1097-Q1100
R74-N71
Q1144-E1104
R352-Q353
R21-Y1032
R242-Y362
N189-D192
S1149-W1145
S1037-Y1092
R303-W356
R80-Y84
Q1012-Y913
S341-Y917
Q1035-Q1038
S308-Y304
R21-W1098
T20-W1098
R352-W1145
Q1144-S1141
R29-Y28
R933-C866
Q207-E92
Q353-E92
N1138-Y917
Q1100-H139
R303-Q359
R297-Y301
H147-Y84
N1148-Q1144
T360-W356
Q250-K246
Q237-S313
Q1038-T845
T1134-Y917
R25-Y1032
Q207-Y89
R248-Q359
T845-W846
Q359-W356
D1152-Y1092
R1030-W846
N189-E193
K298-Y247
R80-Q151
No ion/lipid contact
Anion
Lipid
Both
0.0
0.2
0.4
0.6
0.8
1.0
Frequency

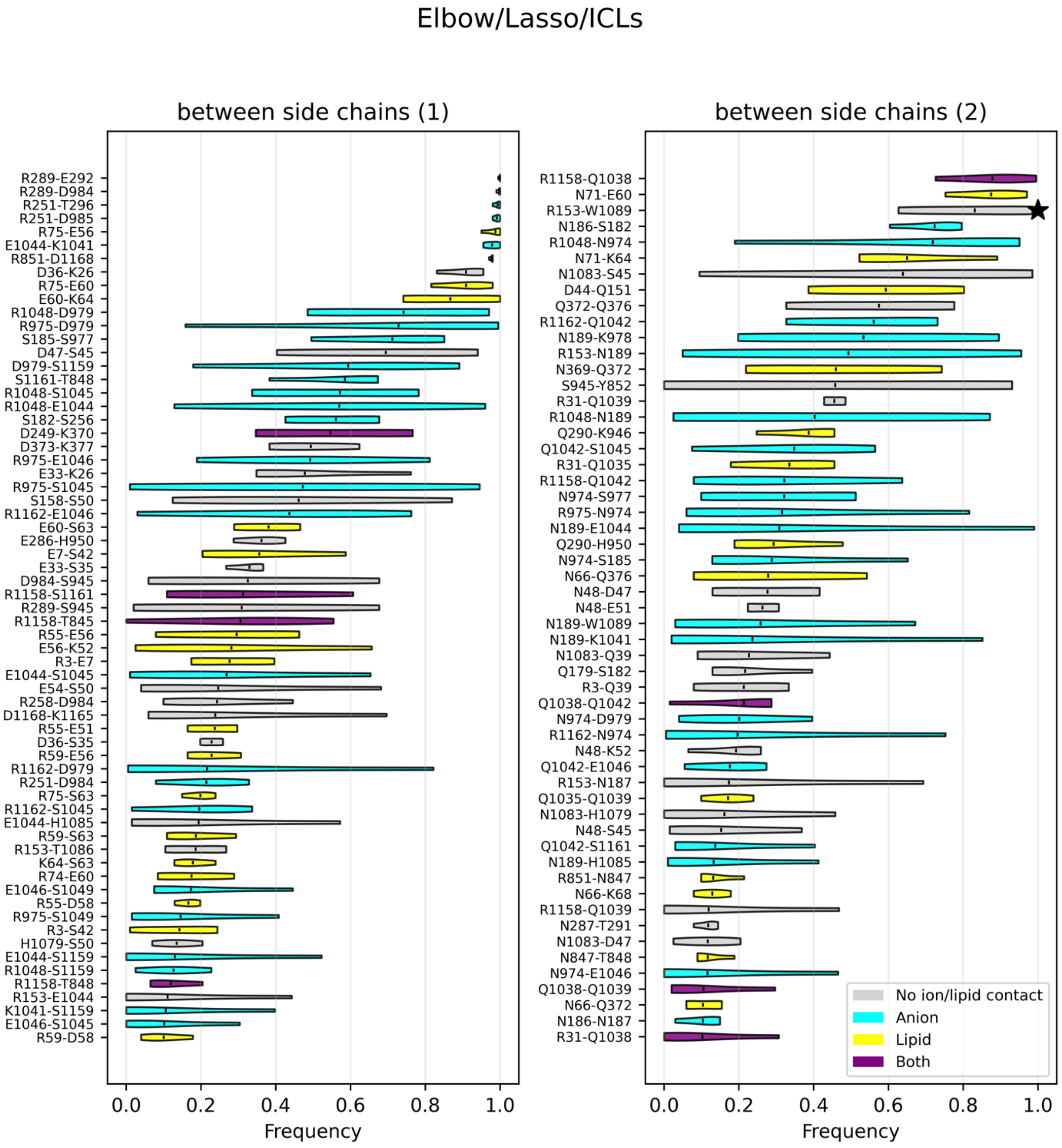
Elbow/Lasso/ICLs
between side chains (1)
between side chains (2)
Frequency
Frequency
No ion/lipid contact
Anion
Lipid
Both

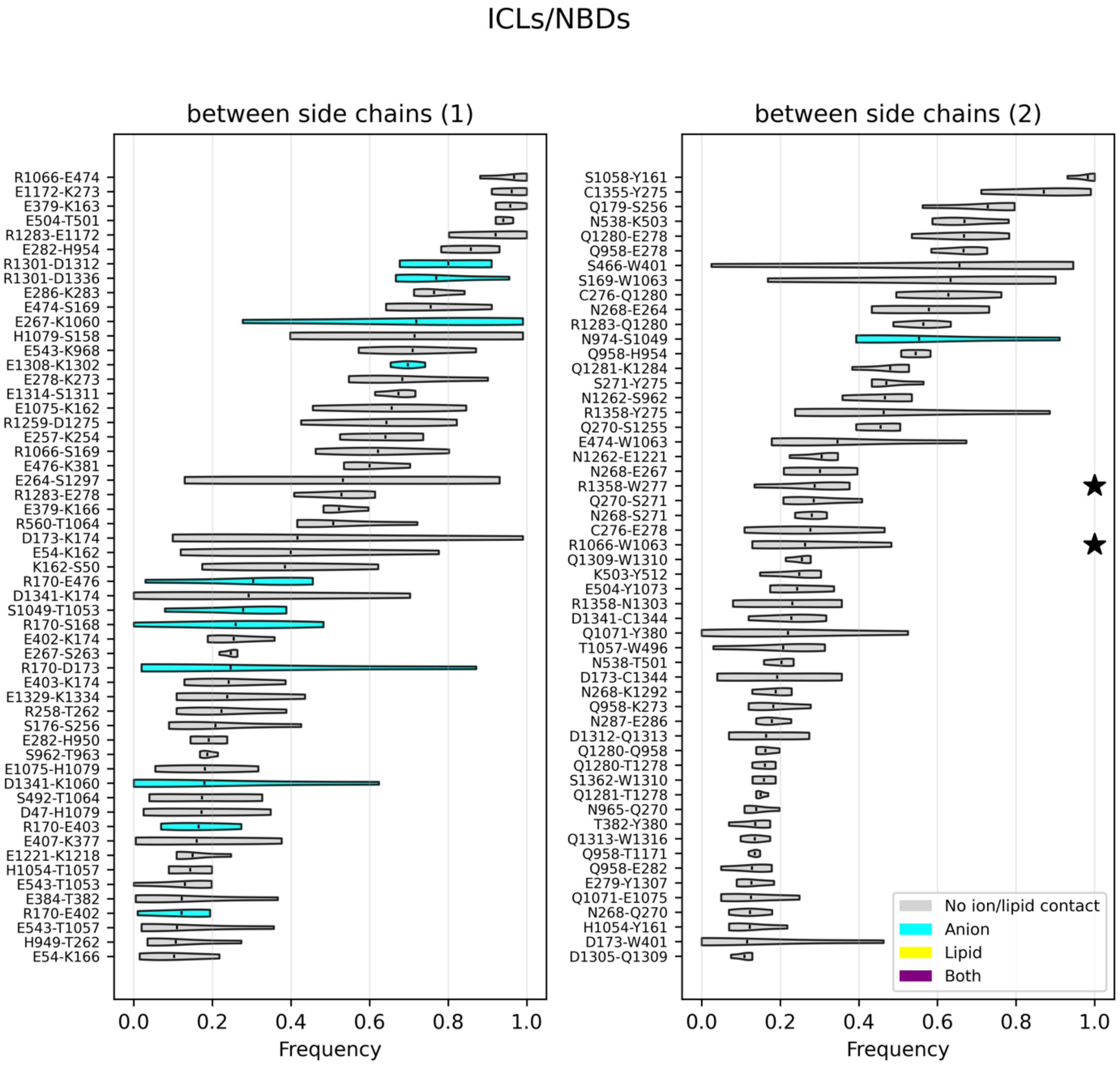
ICLs/NBDs
between side chains (1)
R1066-E474
E1172-K273
E379-K163
E504-T501
R1283-E1172
E282-H954
R1301-D1312
R1301-D1336
E286-K283
E474-S169
E267-K1060
H1079-S158
E543-K968
E1308-K1302
E278-K273
E1314-S1311
E1075-K162
R1259-D1275
E257-K254
R1066-S169
E476-K381
E264-S1297
R1283-E278
E379-K166
R560-T1064
D173-K174
E54-K162
K162-S50
R170-E476
D1341-K174
S1049-T1053
R170-S168
E402-K174
E267-S263
R170-D173
E403-K174
E1329-K1334
R258-T262
S176-S256
E282-H950
S962-T963
E1075-H1079
D1341-K1060
S492-T1064
D47-H1079
R170-E403
E407-K377
E1221-K1218
H1054-T1057
E543-T1053
E384-T382
R170-E402
E543-T1057
H949-T262
E54-K166
0.0
0.2
0.4
0.6
0.8
1.0
Frequency
between side chains (2)
S1058-Y161
C1355-Y275
Q179-S256
N538-K503
Q1280-E278
Q958-E278
S466-W401
S169-W1063
C276-Q1280
N268-E264
R1283-Q1280
N974-S1049
Q958-H954
Q1281-K1284
S271-Y275
N1262-S962
R1358-Y275
Q270-S1255
E474-W1063
N1262-E1221
N268-E267
R1358-W277
Q270-S271
N268-S271
C276-E278
R1066-W1063
Q1309-W1310
K503-Y512
E504-Y1073
R1358-N1303
D1341-C1344
Q1071-Y380
T1057-W496
N538-T501
D173-C1344
N268-K1292
Q958-K273
N287-E286
D1312-Q1313
Q1280-Q958
Q1280-T1278
S1362-W1310
Q1281-T1278
N965-Q270
T382-Y380
Q1313-W1316
Q958-T1171
Q958-E282
E279-Y1307
Q1071-E1075
N268-Q270
H1054-Y161
D173-W401
D1305-Q1309
No ion/lipid contact
Anion
Lipid
Both
0.0
0.2
0.4
0.6
0.8
1.0
Frequency

NBDs

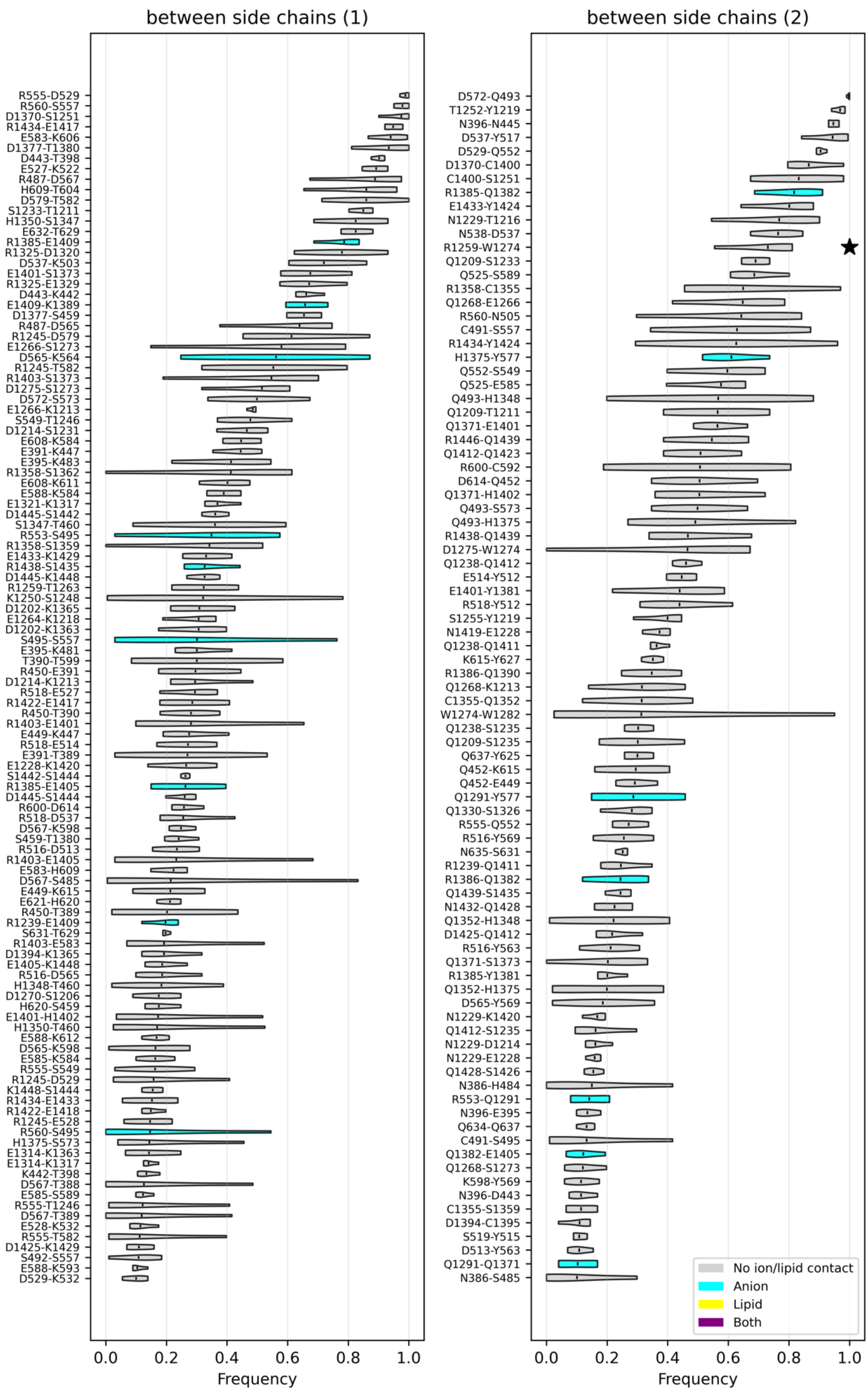

between side chains (1)
between side chains (2)
Frequency
Frequency
No ion/lipid contact
Anion
Lipid
Both

**Figure S9. Detailed views of contacts in the different regions of the human CFTR 3D structure.** These interactions are visualized on the human CFTR 3D structure after 500 ns of MD simulation (apo condition replica 1).

A) **Elbow-lasso-ICLs** (region colored in yellow in Figure 1): networks present at the level of the TM4/TM6 and TM10/TM12. Interactions highlighted in green and pink (TM4/TM6 portal) are stable and transient, respectively.

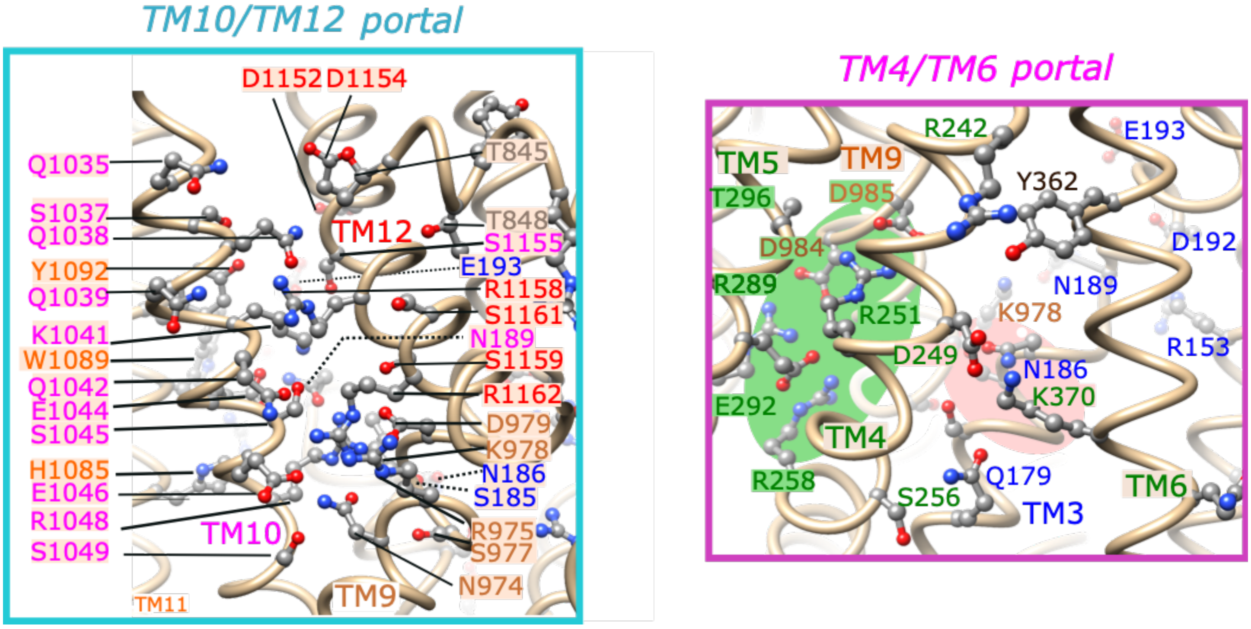


B) **ICLs:NBDs interface** (region colored in green in Figure 1): networks present at the level of the ICL2-ICL3-NBD2 and ICL1-ICL4-NBD1 interfaces. Interactions highlighted in green and pink (TM4/TM6 portal) are stable and transient, respectively.

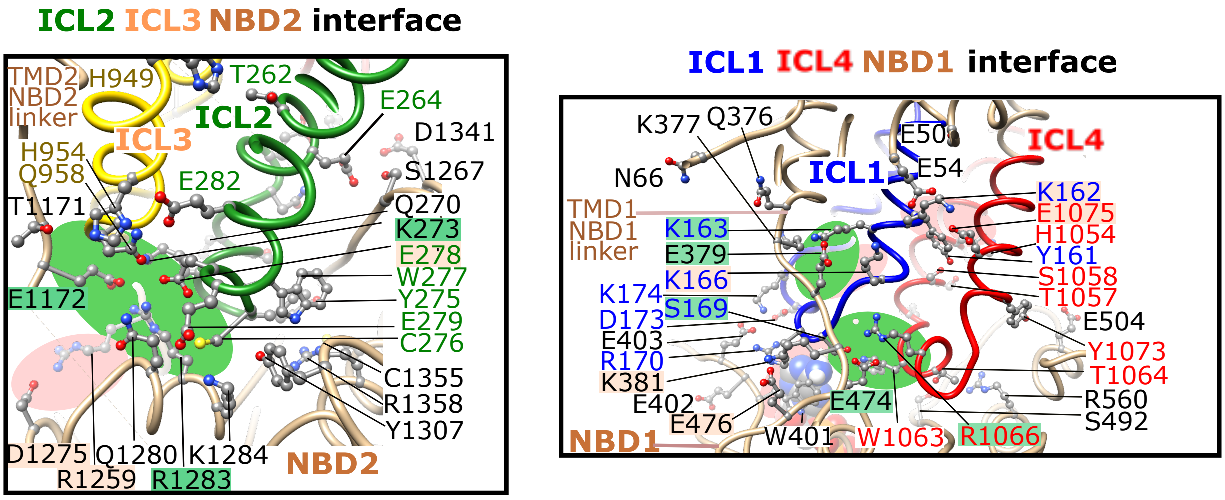

C) **NBD1–NBD2 interface** (region colored in blue in Figure 1). Transient electrostatic interactions involving, among the most striking pairs, the conserved D-loop aspartates ($D579_{NBD1}$ and $D1377_{NBD2}$) and residues just upstream of the Walker A motifs ($S459_{NBD1}$ and $R1245_{NBD2}$), as well as the homologous $Y577_{NBD1}$ and $H1375_{NBD2}$, interacting with each other and with the Q-loop $Q1291_{NBD2}$ and $Q493_{NBD1}$, respectively. $H1348_{NBD2}$ (degenerate ABC signature) can form an alternative bond with $Q493_{NBD1}$ to that established with $H1375_{NBD2}$. These pairs are highlighted in dark blue (NBD1) and magenta (NBD2) within pink circles, relative to residues of the conserved ABC motifs (A-loop, Q-loop, D-loop, H-loop, Walker A, Walker B (E1371Q) and ABC signature, in light blue (NBD1) and orange (NBD2)). Most of these electrostatic interactions observed in the MD simulations (top) were not observed in the cryo-EM 3D structure of human CFTR (bottom, pdb 6OP2). ATP (stick representation) is bound at both ATP-binding sites 1 and 2.

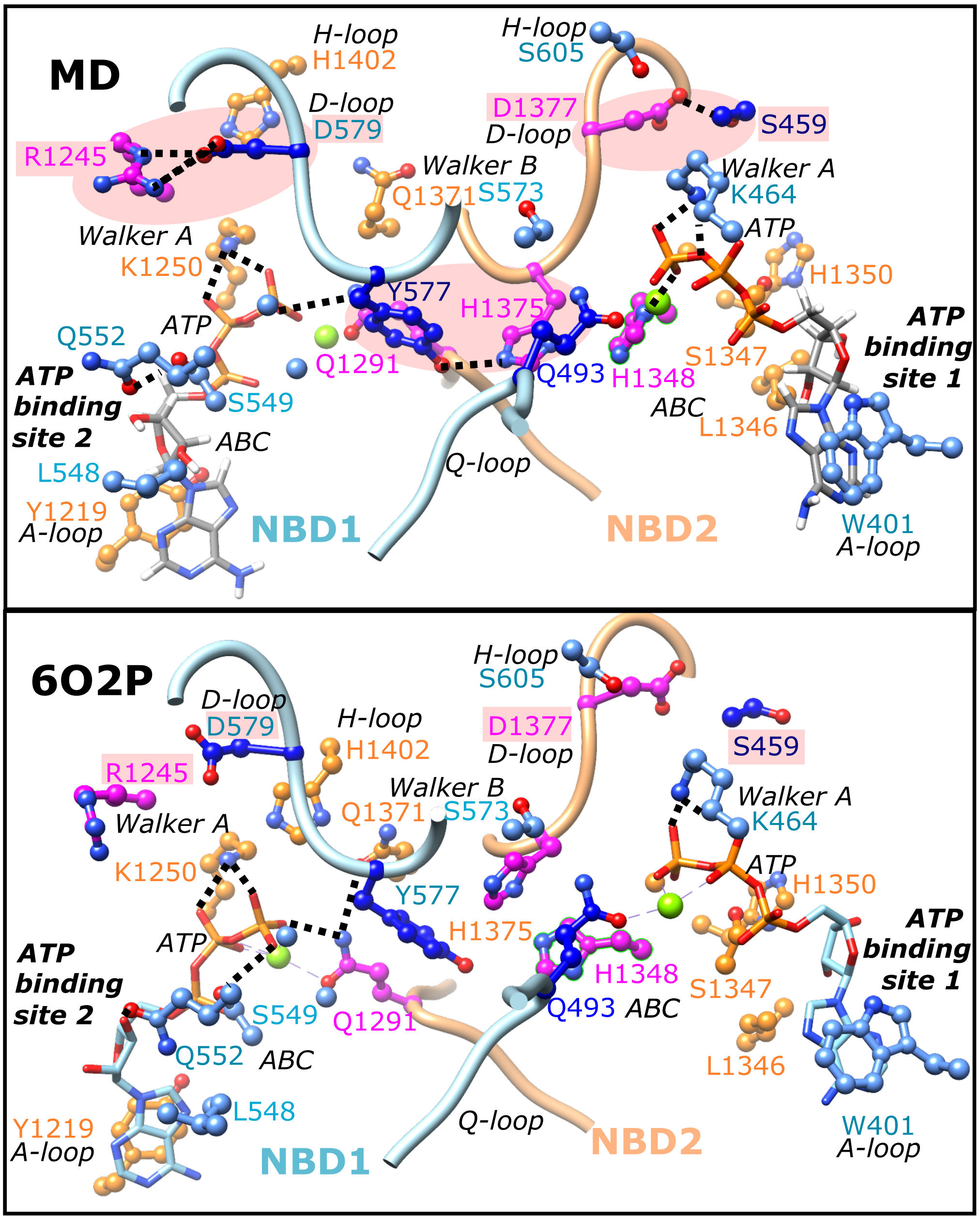

**Figure S10. Residues involved in anion interactions and their overlap with inter-residue and lipid contacts – MD simulations in apo systems (4 replica).** Violin plots showing the distribution of frequencies of residues interacting with anions (chloride (CL) or bicarbonate (CO3)) in the 4 replica (Apo conditions), presented in the descending order of frequency means (bars). In each of the MD simulations, the contact frequencies with anions are summed. Distributions are color-coded based on whether the interacting amino acid is also involved in inter-residue contacts (side-chain/side-chain, red), in lipid interactions (yellow), in both inter-residue and lipid interactions (black) or only in anion contacts (cyan).

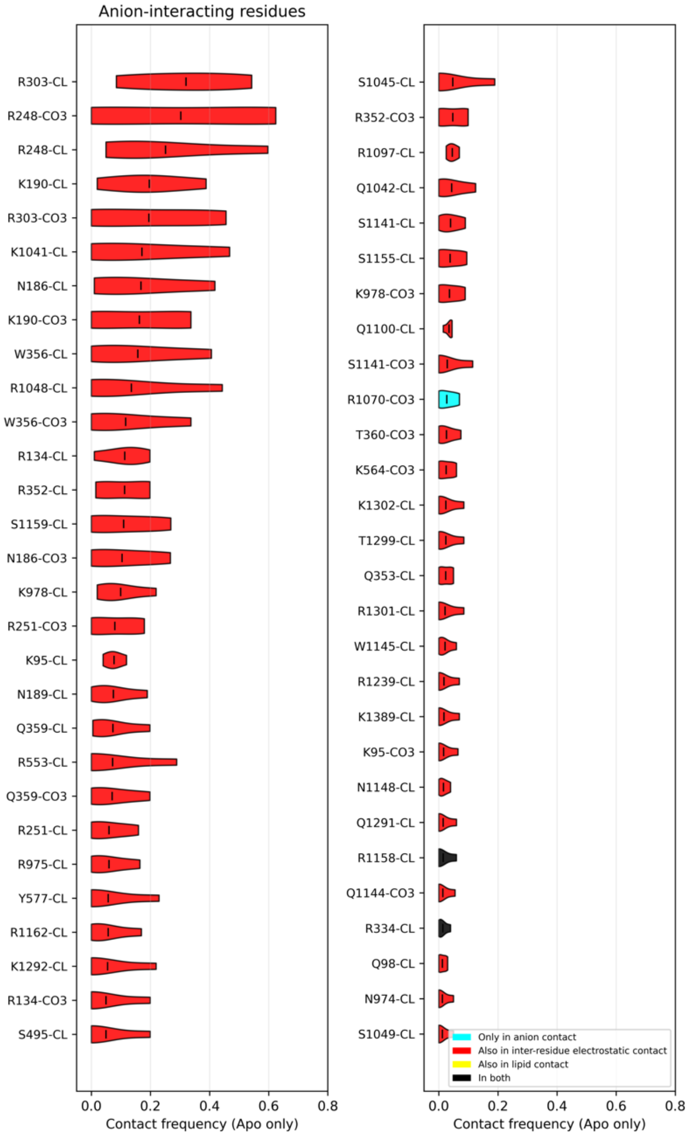

**Figure S11. Most frequent residues involved in lipid headgroups interactions and their overlap with inter-residue and anion contacts – MD simulations in apo systems (4 replica).** Violin plots showing the distribution of frequencies of residues interacting with lipid headgroups in the 4 replica (Apo conditions), presented in the descending order of frequency means (bars). In each of the MD simulations, the contact frequencies with lipids are summed. Colors indicate whether the amino acid is also involved in inter-residue electrostatic interactions (side-chain/side-chain, red), anion interactions (cyan), both inter-residue and anion interactions (black) or only lipid interactions (yellow). Note: the mean frequency for a residue can exceed 1.0 because mean values are summed across lipid headgroup types (e.g., CHL, POC, POE, POS, DSM). POC, POS, POE stand for POPC, POPS and POPE, respectively.

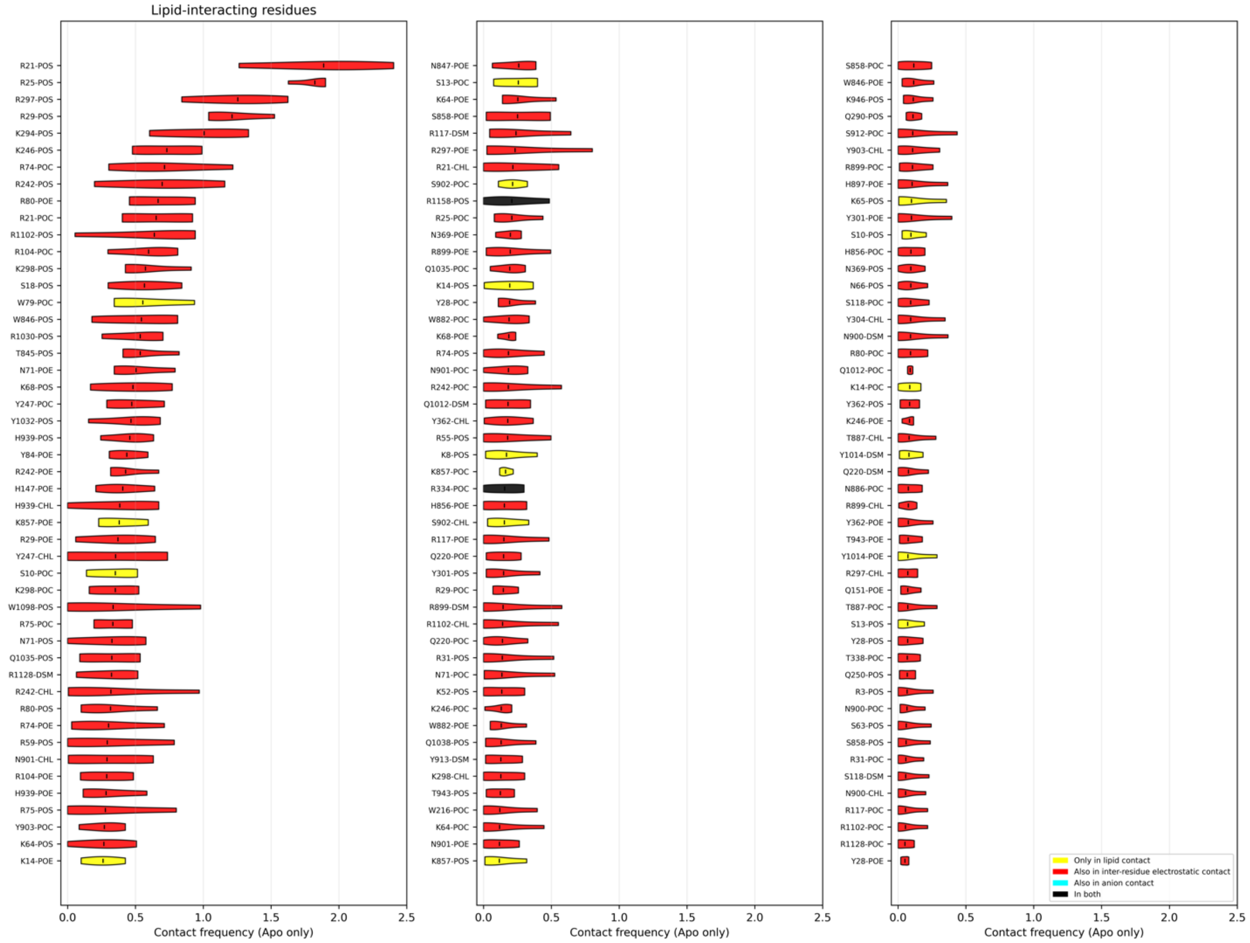

**Figure S12. Comparison of the contact frequencies between the Apo and VX-770-bound conditions.**

**A. Side-chain/side-chain interactions.** Each point represents a specific residue pair; x-axis shows average contact frequency across APO replicates, y-axis across VX replicates. Points above the diagonal indicate increased frequency upon VX-770 binding. Contacts with Δfrequency > 0.3 are labeled.

**B. Interactions with anions.** Total anion contact frequency per amino acid residue, summed across individual simulations and averaged per condition. VX-770 bound vs apo conditions are compared, with points colored by gain (blue) or loss (red) of interaction strength. Contacts with Δfrequency > 0.2 are labeled.

**C. Interactions with lipids (per lipid type).** Lipid contact frequency with charged/polar residue, summed across molecules of the same lipid type within each simulation and averaged per condition (Apo/VX). Residues with VX-770 related shifts (Δfrequency > 0.2) are highlighted. POC, POE and POS stand for POPC, POPE and POPS, respectively.

**A.**

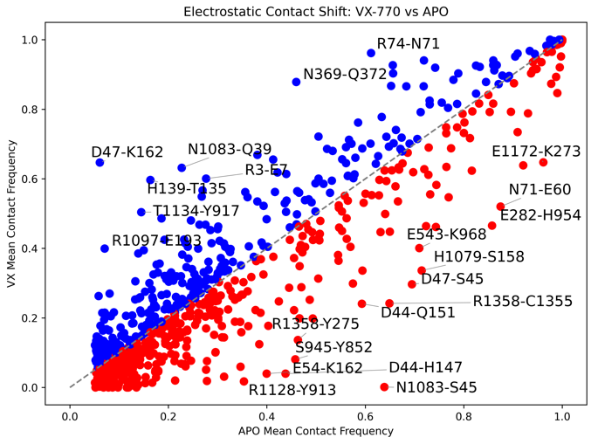


**B.**

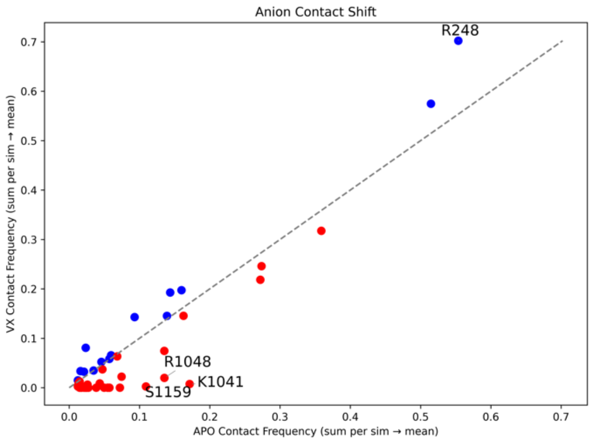

c.

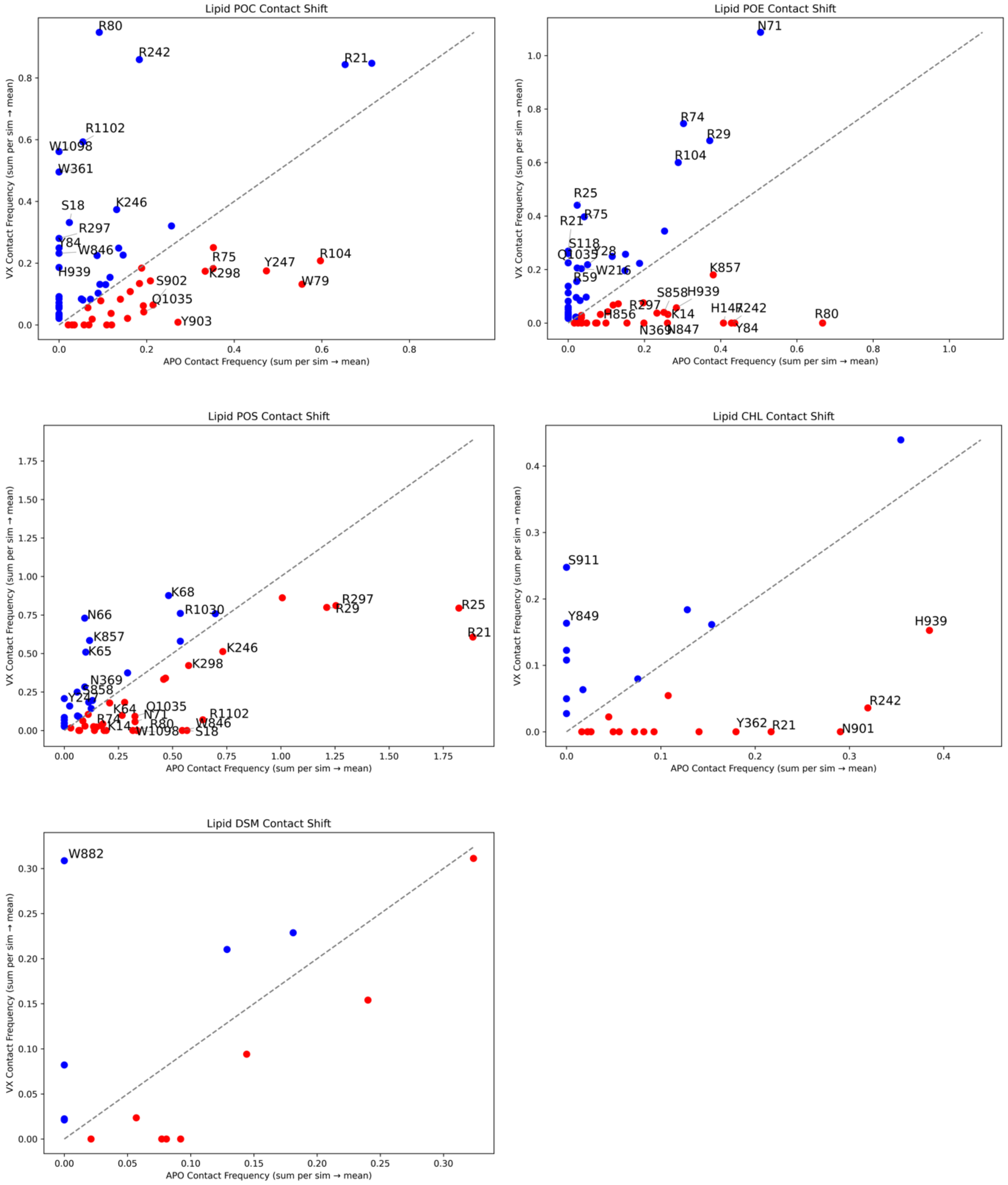
Lipid POC Contact Shift
R80
R242
R21
R1102
W1098
W361
S18
K246
R297
Y84
W846
H939
R75
Y247
R104
R298
S902
W79
Q1035
Y903
Lipid POE Contact Shift
N71
R74
R29
R104
R25
R75
R21
S118
Q1035
Y28
R59
W216
K857
S858
H939
R297
H856
K14
H147
R242
N369
N847
Y84
R80
Lipid POS Contact Shift
K68
N66
R1030
K857
K65
R297
R29
R25
R21
K246
K298
N369
S858
Y24
K64
Q1035
N71
R1102
R74
K14
R80
W846
W1098
S18
Lipid CHL Contact Shift
S911
Y849
H939
R242
Y362
R21
N901
Lipid DSM Contact Shift
W882
VX Contact Frequency (sum per sim → mean)
APO Contact Frequency (sum per sim → mean)

**Figure S13: Residues involved in multiple interaction types.** Bar plot showing the number of residues involved in pairwise or three-way interactions (with another amino acid, anion and/or lipid). Residue names are displayed within each bar.

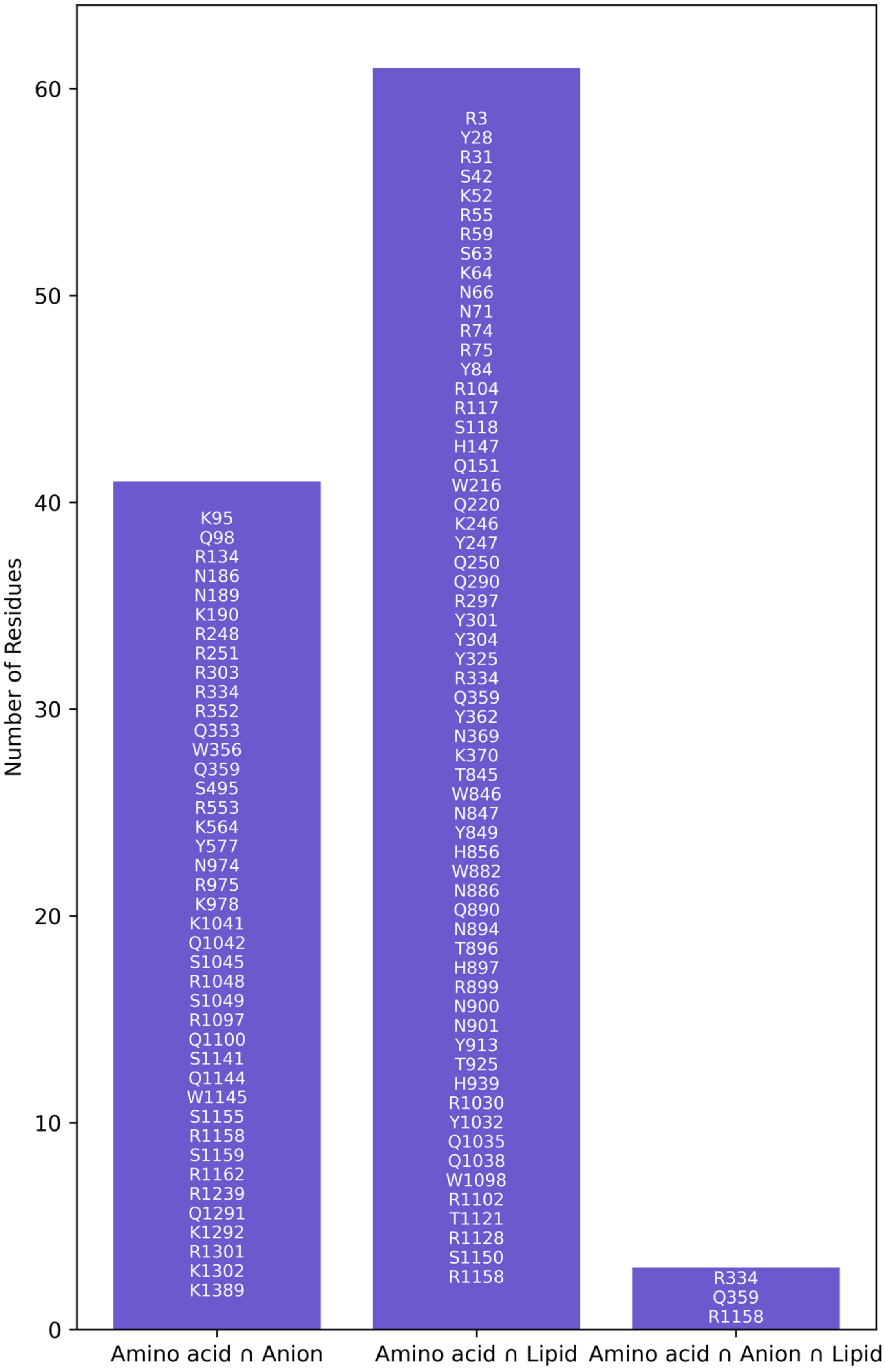

#### Supplementary Tables

**Table S1: MD Simulations of human CFTR.**
Each column corresponds to one simulation. The thermostat and barostat used are indicated. N_CL- is the number of chloride ions and N_HCO3- is the number of bicarbonate ions. Simulation lengths are reported in nanoseconds (ns). Abbreviation: NH = Nose-Hoover.

| | Apo1 | Apo2 | Apo3 | Apo4 | VX1 | VX2 | VX3 | VX4 |
|---|---|---|---|---|---|---|---|---|
| Thermostat | NH | NH | V-rescale | V-rescale | NH | NH | V-rescale | V-rescale |
| Barostat | PR | PR | C-rescale | C-rescale | PR | PR | C-rescale | C-rescale |
| N_CL- | 196 | 196 | 196 | 196 | 195 | 195 | 195 | 195 |
| N_HCO3- | 1 | 1 | 1 | 1 | 1 | 1 | 1 | 1 |
| Sim Time (ns) | 500 | 500 | 1000 | 1000 | 500 | 500 | 1000 | 1000 |
| MD Tool | Gromacs 2020.3 | Gromacs 2023.2 | Gromacs 2023.2 | Gromacs 2023.2 | Gromacs 2020.3 | Gromacs 2023.2 | Gromacs 2023.2 | Gromacs 2023.2 |

**Table S2: Amino acids involved in TM α-helix irregularities and involvement of charged/polar side chains in side-chain/side-chain contacts.**
This table presents CFTR residues that exhibit α-helix irregularities, defined as maintaining an α-helical conformation for less than 80% of the simulation time across molecular dynamics simulations. The columns Apo1–Apo4 and VX1–VX4 report the frequencies of α-helical conformations for each residue in the respective replicates. Residues marked with * indicate an α-helix frequency > 0.8. For charged/polar residues within these irregular regions that participate in amino acid/amino acid contact, the contact pair columns list the interacting partners along with their average contact frequency. Additionally (last columns), the table reports the frequency of contacts formed with anions for these residues.

*See four following pages*

| | Alpha helical conformation | | | | | | | | | Electrostatic contact | | | | | | | | Anion contact | | | | | | | | |
|---|---|---|---|---|---|---|---|---|---|---|---|---|---|---|---|---|---|---|---|---|---|---|---|---|---|
| Residue | Apo1 | Apo2 | Apo3 | Apo4 | VX1 | VX2 | VX3 | VX4 | Contact pair | Apo1 | Apo2 | Apo3 | Apo4 | VX1 | VX2 | VX3 | VX4 | Apo1 | Apo2 | Apo3 | Apo4 | VX1 | VX2 | VX3 | VX4 | TM |
| F78 | * | * | * | * | * | 0.1 | * | * | | | | | | | | | | | | | | | | | | TM1 |
| Y84 | * | * | * | * | 0.8 | * | * | * | | | | | | | | | | 0.0 | 0.0 | 0.0 | 0.0 | 0.0 | 0.0 | 0.0 | 0.0 | TM1 |
| V97 | 0.8 | 0.7 | * | 0.6 | 0.7 | * | 0.6 | 0.8 | | | | | | | | | | | | | | | | | | TM1 |
| L101 | * | * | * | 0.8 | * | * | * | * | | | | | | | | | | | | | | | | | | TM1 |
| A107 | 0.8 | 0.8 | 0.7 | 0.5 | 0.4 | 0.3 | 0.5 | * | | | | | | | | | | | | | | | | | | TM1 |
| R117 | 0.1 | 0.2 | 0.8 | 0.0 | 0.5 | 0.5 | * | 0.7 | | | | | | | | | | | | | | | | | | TM2 |
| | | | | | | | | | R117-D112 | 0.0 | 0.0 | 0.0 | 0.5 | 0.0 | 0.2 | 0.0 | 0.0 | | | | | | | | | |
| | | | | | | | | | R117-E115 | 0.0 | 0.0 | 0.0 | 0.0 | 0.1 | 0.0 | 0.0 | 0.1 | | | | | | | | | |
| | | | | | | | | | N113-R117 | 0.1 | 0.1 | 0.0 | 0.1 | 0.1 | 0.3 | 0.2 | 0.1 | | | | | | | | | |
| | | | | | | | | | Y109-R117 | 0.0 | 0.0 | 0.3 | 0.0 | 0.0 | 0.0 | 0.0 | 0.0 | | | | | | | | | |
| | | | | | | | | | R117-T1122 | 0.8 | 0.3 | 0.0 | 0.1 | 0.1 | 0.2 | 0.1 | 0.0 | | | | | | | | | |
| | | | | | | | | | R117-E1124 | 0.0 | 0.0 | 0.0 | 0.0 | 0.0 | 0.0 | 0.2 | 0.1 | | | | | | | | | |
| S118 | 0.1 | 0.7 | 0.8 | 0.0 | 0.6 | 0.8 | * | * | | | | | | | | | | | | | | | | | | TM2 |
| | | | | | | | | | S118-E116 | 0.0 | 0.3 | 0.0 | 0.0 | 0.0 | 0.0 | 0.0 | 0.0 | | | | | | | | | |
| | | | | | | | | | Y122-S118 | 0.0 | 0.1 | 0.0 | 0.0 | 0.0 | 0.1 | 0.0 | 0.1 | | | | | | | | | |
| I119 | 0.1 | 0.7 | * | 0.2 | 0.7 | * | * | * | | | | | | | | | | | | | | | | | | TM2 |
| L137 | * | * | * | * | 0.7 | * | * | * | | | | | | | | | | | | | | | | | | TM2 |
| L138 | * | * | * | * | 0.7 | * | * | | | | | | | | | | | | | | | | | | | TM2 |
| L165 | 0.6 | 0.6 | 0.8 | 0.8 | * | * | 0.7 | * | | | | | | | | | | | | | | | | | | TM2 |
| N186 | * | 0.8 | 0.8 | * | * | 0.7 | 0.7 | 0.6 | | | | | | | | | | 0.0 | 0.3 | 0.4 | 0.4 | 0.2 | 0.2 | 0.0 | 0.4 | TM3 |
| | | | | | | | | | N186-S182 | 0.6 | 0.7 | 0.8 | 0.8 | 0.4 | 0.6 | 0.5 | 0.3 | | | | | | | | | |
| | | | | | | | | | N186-S185 | 0.1 | 0.0 | 0.0 | 0.0 | 0.1 | 0.1 | 0.2 | 0.1 | | | | | | | | | |
| | | | | | | | | | N186-N187 | 0.0 | 0.1 | 0.1 | 0.1 | 0.2 | 0.0 | 0.1 | 0.4 | | | | | | | | | |
| | | | | | | | | | N186-N189 | 0.0 | 0.0 | 0.0 | 0.0 | 0.1 | 0.0 | 0.1 | 0.0 | | | | | | | | | |
| | | | | | | | | | N186-K190 | 0.0 | 0.1 | 0.1 | 0.1 | 0.0 | 0.0 | 0.0 | 0.3 | | | | | | | | | |
| | | | | | | | | | N186-R248 | 0.1 | 0.0 | 0.0 | 0.0 | 0.1 | 0.1 | 0.0 | 0.0 | | | | | | | | | |
| | | | | | | | | | N186-S256 | 0.0 | 0.1 | 0.0 | 0.0 | 0.0 | 0.0 | 0.3 | 0.0 | | | | | | | | | |
| | | | | | | | | | N186-D363 | 0.0 | 0.0 | 0.0 | 0.0 | 0.1 | 0.0 | 0.0 | 0.1 | | | | | | | | | |
| | | | | | | | | | N186-S977 | 0.0 | 0.0 | 0.1 | 0.0 | 0.1 | 0.0 | 0.1 | 0.0 | | | | | | | | | |
| N187 | * | 0.2 | 0.2 | 0.7 | 0.5 | 0.3 | 0.3 | 0.4 | | | | | | | | | | | | | | | | | | TM3 |
| | | | | | | | | | N187-R153 | 0.7 | 0.0 | 0.0 | 0.0 | 0.2 | 0.0 | 0.0 | 0.0 | | | | | | | | | |
| | | | | | | | | | N186-N187 | 0.0 | 0.1 | 0.1 | 0.1 | 0.2 | 0.0 | 0.1 | 0.4 | | | | | | | | | |
| L188 | 0.0 | 0.1 | 0.0 | 0.4 | 0.3 | 0.0 | 0.1 | 0.0 | | | | | | | | | | | | | | | | | | TM3 |
| N189 | 0.0 | 0.0 | 0.0 | 0.0 | 0.2 | 0.0 | 0.0 | 0.0 | | | | | | | | | | 0.0 | 0.1 | 0.0 | 0.2 | 0.0 | 0.0 | 0.0 | 0.0 | TM3 |
| | | | | | | | | | N189-R153 | 0.0 | 0.3 | 0.7 | 1.0 | 0.1 | 0.4 | 0.1 | 0.2 | | | | | | | | | |
| | | | | | | | | | N189-S185 | 0.0 | 0.0 | 0.0 | 0.0 | 0.1 | 0.0 | 0.0 | 0.0 | | | | | | | | | |
| | | | | | | | | | N186-N189 | 0.0 | 0.0 | 0.0 | 0.0 | 0.1 | 0.0 | 0.1 | 0.0 | | | | | | | | | |
| | | | | | | | | | N189-D192 | 0.0 | 0.3 | 0.7 | 1.0 | 0.0 | 0.3 | 0.0 | 0.7 | | | | | | | | | |
| | | | | | | | | | N189-E193 | 0.0 | 0.3 | 0.1 | 0.0 | 0.0 | 0.0 | 0.0 | 0.1 | | | | | | | | | |
| | | | | | | | | | N189-K978 | 0.2 | 0.7 | 0.3 | 0.9 | 0.6 | 0.6 | 0.9 | 0.7 | | | | | | | | | |
| | | | | | | | | | N189-K1041 | 0.9 | 0.0 | 0.0 | 0.0 | 0.2 | 0.1 | 0.0 | 0.1 | | | | | | | | | |
| | | | | | | | | | N189-E1044 | 1.0 | 0.1 | 0.1 | 0.0 | 0.6 | 0.4 | 0.1 | 0.3 | | | | | | | | | |
| | | | | | | | | | N189-R1048 | 0.9 | 0.1 | 0.0 | 0.6 | 0.5 | 0.0 | 0.0 | 0.1 | | | | | | | | | |
| | | | | | | | | | N189-H1085 | 0.0 | 0.1 | 0.4 | 0.0 | 0.3 | 0.3 | 0.6 | 0.0 | | | | | | | | | |
| | | | | | | | | | N189-W1089 | 0.0 | 0.3 | 0.7 | 0.1 | 0.1 | 0.6 | 0.1 | 0.0 | | | | | | | | | |
| K190 | 0.1 | 0.3 | 0.3 | 0.0 | 0.3 | 0.4 | 0.0 | 0.0 | | | | | | | | | | 0.2 | 0.4 | 0.5 | 0.4 | 0.2 | 0.5 | 0.1 | 0.4 | TM3 |
| | | | | | | | | | N186-K190 | 0.0 | 0.1 | 0.1 | 0.1 | 0.0 | 0.0 | 0.0 | 0.3 | | | | | | | | | |
| | | | | | | | | | K190-E193 | 0.8 | 0.0 | 0.1 | 0.0 | 0.6 | 0.0 | 0.4 | 0.0 | | | | | | | | | |
| | | | | | | | | | K190-D363 | 0.1 | 1.0 | 0.9 | 1.0 | 0.1 | 1.0 | 0.4 | 1.0 | | | | | | | | | |
| | | | | | | | | | K190-D1152 | 0.3 | 0.0 | 0.0 | 0.0 | 0.0 | 0.0 | 0.0 | 0.0 | | | | | | | | | |
| F191 | * | 0.3 | 0.3 | 0.1 | * | 0.4 | 0.3 | 0.1 | | | | | | | | | | | | | | | | | | TM3 |
| D192 | * | 0.3 | 0.3 | 0.1 | * | 0.5 | 0.5 | 0.1 | | | | | | | | | | | | | | | | | | TM3 |
| | | | | | | | | | H146-D192 | 0.0 | 0.0 | 0.0 | 0.0 | 0.4 | 0.0 | 0.0 | 0.0 | | | | | | | | | |
| | | | | | | | | | N189-D192 | 0.0 | 0.3 | 0.7 | 1.0 | 0.0 | 0.3 | 0.0 | 0.7 | | | | | | | | | |
| | | | | | | | | | R153-D192 | 1.0 | 1.0 | 1.0 | 1.0 | 0.8 | 1.0 | 1.0 | 1.0 | | | | | | | | | |
| | | | | | | | | | W1089-D192 | 1.0 | 1.0 | 1.0 | 0.5 | 1.0 | 1.0 | 1.0 | 0.7 | | | | | | | | | |
| E193 | * | 0.3 | 0.3 | 0.1 | * | 0.6 | 0.5 | 0.1 | | | | | | | | | | | | | | | | | | TM3 |
| | | | | | | | | | H146-E193 | 0.1 | 0.1 | 0.1 | 0.1 | 0.0 | 0.0 | 0.0 | 0.1 | | | | | | | | | |
| | | | | | | | | | K190-E193 | 0.8 | 0.0 | 0.1 | 0.0 | 0.6 | 0.0 | 0.4 | 0.0 | | | | | | | | | |
| | | | | | | | | | N189-E193 | 0.0 | 0.3 | 0.1 | 0.0 | 0.0 | 0.0 | 0.0 | 0.1 | | | | | | | | | |
| | | | | | | | | | R1097-E193 | 0.1 | 0.1 | 0.0 | 0.0 | 0.3 | 0.6 | 0.3 | 0.3 | | | | | | | | | |
| G194 | * | 0.5 | 0.5 | * | * | * | 0.7 | 0.3 | | | | | | | | | | | | | | | | | | TM3 |
| A198 | * | * | * | * | * | 0.7 | * | 0.8 | | | | | | | | | | | | | | | | | | TM3 |
| H199 | 0.2 | 0.4 | 0.4 | 0.5 | 0.4 | 0.3 | 0.2 | 0.3 | | | | | | | | | | | | | | | | | | TM3 |
| | | | | | | | | | W202-H199 | 1.0 | 1.0 | 1.0 | 1.0 | 1.0 | 0.9 | 0.9 | 0.8 | | | | | | | | | |
| F200 | 0.2 | 0.5 | 0.6 | 0.6 | 0.6 | 0.7 | 0.7 | 0.7 | | | | | | | | | | | | | | | | | | TM3 |

| | Alpha helical conformation | | | | | | | | | Electrostatic contact | | | | | | | | Anion contact | | | | | | | | |
|---|---|---|---|---|---|---|---|---|---|---|---|---|---|---|---|---|---|---|---|---|---|---|---|---|---|---|
| Residue | Apo1 | Apo2 | Apo3 | Apo4 | VX1 | VX2 | VX3 | VX4 | Contact pair | Apo1 | Apo2 | Apo3 | Apo4 | VX1 | VX2 | VX3 | VX4 | Apo1 | Apo2 | Apo3 | Apo4 | VX1 | VX2 | VX3 | VX4 | TM |
| **V201** | 0.2 | 0.5 | 0.5 | 0.6 | 0.6 | 0.6 | 0.7 | 0.7 | | | | | | | | | | | | | | | | | | TM3 |
| **W202** | 0.2 | 0.5 | 0.5 | 0.6 | 0.6 | 0.6 | 0.7 | 0.7 | | | | | | | | | | | | | | | | | | TM3 |
| | | | | | | | | | W202-H199 | 1.0 | 1.0 | 1.0 | 1.0 | 1.0 | 0.9 | 0.9 | 0.8 | | | | | | | | | |
| **I203** | 0.7 | * | * | * | * | * | * | * | | | | | | | | | | | | | | | | | | TM3 |
| **I215** | * | * | * | * | * | * | * | 0.8 | | | | | | | | | | | | | | | | | | TM3 |
| **A223** | * | * | 0.8 | 0.8 | * | * | 0.6 | * | | | | | | | | | | | | | | | | | | TM4 |
| **S313** | 0.5 | 0.2 | 0.4 | 0.6 | 0.5 | 0.8 | 0.7 | 0.5 | | | | | | | | | | | | | | | | | | TM5 |
| | | | | | | | | | C343-S313 | 0.1 | 0.0 | 0.0 | 0.0 | 0.0 | 0.0 | 0.0 | 0.0 | | | | | | | | | |
| | | | | | | | | | Q237-S313 | 0.1 | 0.5 | 0.0 | 0.0 | 0.3 | 0.0 | 0.2 | 0.0 | | | | | | | | | |
| | | | | | | | | | R347-S313 | 0.4 | 0.2 | 0.1 | 0.1 | 0.1 | 0.0 | 0.0 | 0.0 | | | | | | | | | |
| | | | | | | | | | S313-T351 | 0.9 | 0.5 | 1.0 | 1.0 | 0.4 | 0.8 | 1.0 | 1.0 | | | | | | | | | |
| **G314** | 0.8 | 0.1 | * | * | * | 0.7 | * | * | | | | | | | | | | | | | | | | | | TM5 |
| **F315** | * | * | * | * | * | 0.7 | * | * | | | | | | | | | | | | | | | | | | TM5 |
| **V322** | * | * | * | * | * | * | * | 0.6 | | | | | | | | | | | | | | | | | | TM5 |
| **L323** | * | * | * | * | * | 0.5 | * | 0.6 | | | | | | | | | | | | | | | | | | TM5 |
| **P324** | * | * | * | * | * | 0.5 | * | 0.6 | | | | | | | | | | | | | | | | | | TM5 |
| **Y325** | * | * | * | * | * | 0.6 | * | 0.6 | | | | | | | | | | | | | | | | | | TM5 |
| | | | | | | | | | Q220-Y325 | 0.1 | 0.4 | 0.2 | 0.1 | 0.0 | 0.4 | 0.2 | 0.2 | | | | | | | | | |
| | | | | | | | | | Y325-K329 | 0.2 | 0.1 | 0.5 | 0.4 | 0.2 | 0.1 | 0.2 | 0.0 | | | | | | | | | |
| | | | | | | | | | Y325-K335 | 0.4 | 0.2 | 0.5 | 0.4 | 0.4 | 0.3 | 0.5 | 0.0 | | | | | | | | | |
| **A326** | * | * | * | * | * | 0.4 | * | 0.6 | | | | | | | | | | | | | | | | | | TM5 |
| **L327** | * | 0.8 | * | * | * | 0.3 | * | 0.2 | | | | | | | | | | | | | | | | | | TM5 |
| **I328** | * | 0.8 | * | * | * | 0.3 | * | 0.2 | | | | | | | | | | | | | | | | | | TM5 |
| **K329** | 0.6 | 0.3 | 0.6 | 0.7 | 0.5 | 0.1 | 0.3 | 0.1 | | | | | | | | | | | | | | | | | | TM5 |
| | | | | | | | | | N900-K329 | 0.0 | 0.0 | 0.0 | 0.0 | 0.0 | 0.0 | 0.0 | 0.1 | | | | | | | | | |
| | | | | | | | | | Q220-K329 | 0.1 | 0.2 | 0.1 | 0.2 | 0.1 | 0.0 | 0.0 | 0.0 | | | | | | | | | |
| | | | | | | | | | Y325-K329 | 0.2 | 0.1 | 0.5 | 0.4 | 0.2 | 0.1 | 0.2 | 0.0 | | | | | | | | | |
| **T351** | 0.5 | 0.6 | 0.6 | 0.5 | 0.6 | 0.6 | 0.4 | 0.6 | | | | | | | | | | | | | | | | | | TM6 |
| | | | | | | | | | Q237-T351 | 0.0 | 0.0 | 0.0 | 0.0 | 0.1 | 0.1 | 0.0 | 0.0 | | | | | | | | | |
| | | | | | | | | | R347-T351 | 0.2 | 0.1 | 0.0 | 0.0 | 0.0 | 0.0 | 0.0 | 0.0 | | | | | | | | | |
| | | | | | | | | | S313-T351 | 0.9 | 0.5 | 1.0 | 1.0 | 0.4 | 0.8 | 1.0 | 1.0 | | | | | | | | | |
| **L375** | * | * | 0.8 | 0.8 | * | * | * | * | | | | | | | | | | | | | | | | | | TM6 |
| **Q376** | 0.2 | 0.2 | 0.2 | 0.2 | 0.2 | 0.2 | 0.1 | 0.2 | | | | | | | | | | | | | | | | | | TM6 |
| | | | | | | | | | Q376-D373 | 0.1 | 0.0 | 0.1 | 0.1 | 0.0 | 0.1 | 0.1 | 0.2 | | | | | | | | | |
| | | | | | | | | | N66-Q376 | 0.1 | 0.1 | 0.5 | 0.4 | 0.0 | 0.0 | 0.0 | 0.0 | | | | | | | | | |
| | | | | | | | | | Q372-Q376 | 0.5 | 0.3 | 0.7 | 0.8 | 0.2 | 0.6 | 0.2 | 0.2 | | | | | | | | | |
| | | | | | | | | | W57-Q376 | 0.0 | 0.1 | 0.0 | 0.0 | 0.5 | 0.1 | 0.6 | 0.3 | | | | | | | | | |
| **S858** | * | * | * | * | 0.5 | * | * | 0.8 | | | | | | | | | | | | | | | | | | TM7 |
| | | | | | | | | | H856-S858 | 0.2 | 0.2 | 0.3 | 0.2 | 0.3 | 0.4 | 0.1 | 0.2 | | | | | | | | | |
| **S909** | 0.1 | 0.0 | 0.0 | 0.4 | 0.0 | 0.0 | 0.0 | * | | | | | | | | | | | | | | | | | | TM8 |
| | | | | | | | | | S909-D110 | 0.0 | 0.0 | 0.0 | 0.0 | 0.0 | 0.0 | 0.4 | 0.0 | | | | | | | | | |
| | | | | | | | | | K892-S909 | 0.0 | 0.0 | 0.1 | 0.0 | 0.1 | 0.0 | 0.0 | 0.0 | | | | | | | | | |
| | | | | | | | | | N894-S909 | 0.0 | 0.0 | 0.1 | 0.0 | 0.2 | 0.0 | 0.0 | 0.0 | | | | | | | | | |
| | | | | | | | | | Q890-S909 | 0.0 | 0.1 | 0.0 | 0.0 | 0.1 | 0.0 | 0.0 | 0.0 | | | | | | | | | |
| | | | | | | | | | R1128-S909 | 0.3 | 0.2 | 0.2 | 0.0 | 0.0 | 0.0 | 0.6 | 0.1 | | | | | | | | | |
| | | | | | | | | | T896-S909 | 0.0 | 0.2 | 0.0 | 0.0 | 0.0 | 0.0 | 0.0 | 0.0 | | | | | | | | | |
| | | | | | | | | | T908-S909 | 0.3 | 0.2 | 0.2 | 0.0 | 0.1 | 0.1 | 0.0 | 0.0 | | | | | | | | | |
| | | | | | | | | | S909-T910 | 0.1 | 0.2 | 0.4 | 0.4 | 0.1 | 0.1 | 0.4 | 0.2 | | | | | | | | | |
| | | | | | | | | | S909-E1126 | 0.3 | 0.1 | 0.7 | 0.4 | 0.0 | 0.1 | 0.0 | 0.1 | | | | | | | | | |
| **T910** | 0.1 | 0.0 | 0.0 | 0.7 | 0.0 | 0.0 | 0.0 | * | | | | | | | | | | | | | | | | | | TM8 |
| | | | | | | | | | T910-D110 | 0.7 | 0.0 | 0.1 | 0.3 | 0.0 | 0.0 | 0.9 | 0.0 | | | | | | | | | |
| | | | | | | | | | N894-T910 | 0.0 | 0.0 | 0.0 | 0.0 | 0.2 | 0.0 | 0.0 | 0.0 | | | | | | | | | |
| | | | | | | | | | R1128-T910 | 0.0 | 0.2 | 0.1 | 0.0 | 0.0 | 0.0 | 0.9 | 0.0 | | | | | | | | | |
| | | | | | | | | | R334-T910 | 0.1 | 0.0 | 0.0 | 0.0 | 0.0 | 0.0 | 0.0 | 0.0 | | | | | | | | | |
| | | | | | | | | | S909-T910 | 0.1 | 0.2 | 0.4 | 0.4 | 0.1 | 0.1 | 0.4 | 0.2 | | | | | | | | | |
| | | | | | | | | | T908-T910 | 0.3 | 0.0 | 0.6 | 0.0 | 0.5 | 0.0 | 0.4 | 0.0 | | | | | | | | | |
| | | | | | | | | | Y914-T910 | 0.0 | 0.1 | 0.6 | 0.1 | 0.0 | 0.0 | 0.1 | 0.0 | | | | | | | | | |
| | | | | | | | | | T910-S911 | 0.0 | 0.0 | 0.0 | 0.1 | 0.1 | 0.3 | 0.0 | 0.0 | | | | | | | | | |
| | | | | | | | | | T910-S912 | 0.0 | 0.1 | 0.0 | 0.0 | 0.0 | 0.1 | 0.0 | 0.0 | | | | | | | | | |
| | | | | | | | | | T910-E1126 | 0.0 | 0.0 | 0.5 | 0.1 | 0.0 | 0.0 | 0.0 | 0.0 | | | | | | | | | |
| **S911** | * | 0.7 | * | 0.8 | 0.1 | 0.3 | * | * | | | | | | | | | | | | | | | | | | TM8 |
| | | | | | | | | | N886-S911 | 0.1 | 0.5 | 0.0 | 0.2 | 0.3 | 0.1 | 0.0 | 0.1 | | | | | | | | | |
| | | | | | | | | | Q890-S911 | 0.0 | 0.0 | 0.3 | 0.0 | 0.0 | 0.0 | 0.0 | 0.0 | | | | | | | | | |
| | | | | | | | | | R334-S911 | 0.0 | 0.0 | 0.0 | 0.0 | 0.0 | 0.2 | 0.0 | 0.0 | | | | | | | | | |
| | | | | | | | | | T908-S911 | 0.2 | 0.0 | 0.0 | 0.0 | 0.1 | 0.0 | 0.2 | 0.0 | | | | | | | | | |
| | | | | | | | | | T910-S911 | 0.0 | 0.0 | 0.0 | 0.1 | 0.1 | 0.3 | 0.0 | 0.0 | | | | | | | | | |

| | Alpha helical conformation | | | | | | | | | Electrostatic contact | | | | | | | | Anion contact | | | | | | | | |
| --- | --- | --- | --- | --- | --- | --- | --- | --- | --- | --- | --- | --- | --- | --- | --- | --- | --- | --- | --- | --- | --- | --- | --- | --- | --- | --- |
| Residue | Apo1 | Apo2 | Apo3 | Apo4 | VX1 | VX2 | VX3 | VX4 | Contact pair | Apo1 | Apo2 | Apo3 | Apo4 | VX1 | VX2 | VX3 | VX4 | Apo1 | Apo2 | Apo3 | Apo4 | VX1 | VX2 | VX3 | VX4 | TM |
| | | | | | | | | | Y913-S911 | 0.0 | 0.0 | 0.0 | 0.0 | 0.0 | 0.1 | 0.0 | 0.0 | | | | | | | | | |
| | | | | | | | | | S911-S912 | 0.0 | 0.0 | 0.0 | 0.0 | 0.0 | 0.0 | 0.1 | 0.0 | | | | | | | | | |
| S912 | * | * | * | 0.8 | 0.1 | 0.7 | * | * | | | | | | | | | | | | | | | | | | TM8 |
| | | | | | | | | | N886-S912 | 0.1 | 0.2 | 0.0 | 0.4 | 0.1 | 0.2 | 0.2 | 0.5 | | | | | | | | | |
| | | | | | | | | | Q890-S912 | 0.0 | 0.0 | 0.2 | 0.0 | 0.0 | 0.0 | 0.1 | 0.0 | | | | | | | | | |
| | | | | | | | | | R1128-S912 | 0.1 | 0.0 | 0.0 | 0.1 | 0.0 | 0.2 | 0.0 | 0.6 | | | | | | | | | |
| | | | | | | | | | S911-S912 | 0.0 | 0.0 | 0.0 | 0.0 | 0.0 | 0.0 | 0.1 | 0.0 | | | | | | | | | |
| | | | | | | | | | T910-S912 | 0.0 | 0.1 | 0.0 | 0.0 | 0.0 | 0.1 | 0.0 | 0.0 | | | | | | | | | |
| Y913 | * | * | * | 0.8 | 0.1 | * | * | * | | | | | | | | | | | | | | | | | | TM8 |
| | | | | | | | | | Y913-D110 | 0.1 | 0.1 | 0.0 | 0.2 | 0.3 | 0.8 | 0.0 | 0.0 | | | | | | | | | |
| | | | | | | | | | Y913-R334 | 0.0 | 0.0 | 0.0 | 0.0 | 0.1 | 0.2 | 0.0 | 0.0 | | | | | | | | | |
| | | | | | | | | | Y913-S911 | 0.0 | 0.0 | 0.0 | 0.0 | 0.0 | 0.1 | 0.0 | 0.0 | | | | | | | | | |
| | | | | | | | | | Y913-Y914 | 0.1 | 0.1 | 0.0 | 0.0 | 0.9 | 0.0 | 0.0 | 0.0 | | | | | | | | | |
| | | | | | | | | | Y913-Y917 | 0.0 | 0.0 | 0.0 | 0.0 | 0.0 | 0.0 | 0.0 | 0.1 | | | | | | | | | |
| | | | | | | | | | Y913-Q1012 | 0.2 | 0.5 | 0.4 | 0.1 | 0.0 | 0.0 | 0.9 | 0.0 | | | | | | | | | |
| | | | | | | | | | Y913-E1126 | 0.0 | 0.1 | 0.0 | 0.3 | 0.0 | 0.3 | 0.0 | 0.0 | | | | | | | | | |
| | | | | | | | | | Y913-R1128 | 0.4 | 0.4 | 0.4 | 0.3 | 0.0 | 0.0 | 0.0 | 0.0 | | | | | | | | | |
| V922 | 0.4 | 0.4 | 0.4 | 0.7 | 0.5 | 0.5 | 0.5 | 0.4 | | | | | | | | | | | | | | | | | | TM8 |
| L927 | 0.0 | 0.1 | 0.0 | 0.2 | 0.2 | 0.3 | 0.4 | 0.5 | | | | | | | | | | | | | | | | | | TM8 |
| G930 | * | 0.0 | * | 0.4 | * | * | * | * | | | | | | | | | | | | | | | | | | TM8 |
| F931 | 0.6 | 0.4 | 0.0 | 0.5 | 0.7 | 0.5 | 0.8 | 0.7 | | | | | | | | | | | | | | | | | | TM8 |
| F932 | 0.6 | 0.4 | 0.0 | 0.5 | 0.7 | 0.5 | 0.8 | 0.7 | | | | | | | | | | | | | | | | | | TM8 |
| R933 | 0.6 | 0.4 | 0.0 | 0.5 | 0.7 | 0.5 | 0.8 | 0.7 | | | | | | | | | | | | | | | | | | TM8 |
| | | | | | | | | | R933-E873 | 1.0 | 1.0 | 1.0 | 1.0 | 1.0 | 1.0 | 1.0 | 1.0 | | | | | | | | | |
| | | | | | | | | | C866-R933 | 0.3 | 0.3 | 0.1 | 0.3 | 0.5 | 0.1 | 0.1 | 0.0 | | | | | | | | | |
| | | | | | | | | | Q996-R933 | 0.0 | 0.0 | 0.0 | 0.0 | 0.7 | 0.0 | 0.0 | 0.0 | | | | | | | | | |
| K946 | * | * | * | 0.8 | * | * | * | * | | | | | | | | | | | | | | | | | | TM8 |
| | | | | | | | | | Q290-K946 | 0.2 | 0.5 | 0.5 | 0.4 | 0.2 | 0.2 | 0.1 | 0.3 | | | | | | | | | |
| Q958 | 0.4 | 0.4 | 0.6 | 0.5 | 0.4 | 0.4 | 0.6 | 0.3 | | | | | | | | | | | | | | | | | | TM8 |
| | | | | | | | | | Q958-K273 | 0.3 | 0.2 | 0.1 | 0.1 | 0.2 | 0.5 | 0.1 | 0.7 | | | | | | | | | |
| | | | | | | | | | Q958-E278 | 0.6 | 0.7 | 0.7 | 0.7 | 0.3 | 0.5 | 0.7 | 0.6 | | | | | | | | | |
| | | | | | | | | | Q958-E282 | 0.2 | 0.0 | 0.2 | 0.1 | 0.0 | 0.0 | 0.1 | 0.0 | | | | | | | | | |
| | | | | | | | | | Q958-H954 | 0.6 | 0.5 | 0.5 | 0.6 | 0.5 | 0.3 | 0.3 | 0.4 | | | | | | | | | |
| | | | | | | | | | Q958-T1171 | 0.1 | 0.1 | 0.1 | 0.1 | 0.2 | 0.1 | 0.2 | 0.0 | | | | | | | | | |
| | | | | | | | | | Q958-E1172 | 0.1 | 0.1 | 0.0 | 0.1 | 0.1 | 0.1 | 0.0 | 0.0 | | | | | | | | | |
| | | | | | | | | | Q958-Q1280 | 0.2 | 0.1 | 0.2 | 0.1 | 0.3 | 0.1 | 0.1 | 0.0 | | | | | | | | | |
| F976 | * | * | * | 0.7 | * | 0.8 | * | * | | | | | | | | | | | | | | | | | | TM9 |
| S977 | 0.6 | 0.7 | * | 0.6 | * | 0.3 | * | 0.7 | | | | | | | | | | | | | | | | | | TM9 |
| | | | | | | | | | N186-S977 | 0.0 | 0.0 | 0.1 | 0.0 | 0.1 | 0.0 | 0.1 | 0.0 | | | | | | | | | |
| | | | | | | | | | N974-S977 | 0.5 | 0.1 | 0.5 | 0.2 | 0.2 | 0.5 | 0.2 | 0.0 | | | | | | | | | |
| | | | | | | | | | R1048-S977 | 0.0 | 0.0 | 0.0 | 0.0 | 0.0 | 0.1 | 0.0 | 0.0 | | | | | | | | | |
| | | | | | | | | | S182-S977 | 0.0 | 0.0 | 0.0 | 0.0 | 0.2 | 0.0 | 0.0 | 0.2 | | | | | | | | | |
| | | | | | | | | | ùù^ùù`ù | 0.5 | 0.7 | 0.9 | 0.8 | 0.8 | 0.7 | 0.7 | 1.0 | | | | | | | | | |
| K978 | * | * | 0.7 | 0.8 | 0.2 | 0.8 | 0.1 | * | | | | | | | | | | 0.0 | 0.2 | 0.1 | 0.2 | 0.1 | 0.1 | 0.0 | 0.1 | TM9 |
| | | | | | | | | | K978-S185 | 0.2 | 0.0 | 0.0 | 0.0 | 0.1 | 0.1 | 0.3 | 0.1 | | | | | | | | | |
| | | | | | | | | | N189-K978 | 0.2 | 0.7 | 0.3 | 0.9 | 0.6 | 0.6 | 0.9 | 0.7 | | | | | | | | | |
| D979 | * | * | * | * | 0.6 | 0.8 | 0.4 | * | | | | | | | | | | | | | | | | | | TM9 |
| | | | | | | | | | N974-D979 | 0.4 | 0.1 | 0.0 | 0.2 | 0.1 | 0.4 | 0.0 | 0.7 | | | | | | | | | |
| | | | | | | | | | R1048-D979 | 0.5 | 1.0 | 0.9 | 0.6 | 0.8 | 1.0 | 1.0 | 0.9 | | | | | | | | | |
| | | | | | | | | | R1162-D979 | 0.8 | 0.0 | 0.0 | 0.0 | 0.1 | 0.9 | 0.2 | 0.1 | | | | | | | | | |
| | | | | | | | | | R975-D979 | 0.2 | 0.9 | 1.0 | 0.9 | 0.7 | 0.6 | 0.9 | 0.5 | | | | | | | | | |
| | | | | | | | | | S1045-D979 | 0.0 | 0.0 | 0.0 | 0.1 | 0.2 | 0.3 | 0.1 | 0.0 | | | | | | | | | |
| | | | | | | | | | S1159-D979 | 0.9 | 0.6 | 0.2 | 0.7 | 0.8 | 0.6 | 0.7 | 0.6 | | | | | | | | | |
| I980 | * | * | * | * | 0.6 | 0.8 | 0.4 | * | | | | | | | | | | | | | | | | | | TM9 |
| A981 | * | * | * | * | 0.7 | * | 0.4 | * | | | | | | | | | | | | | | | | | | TM9 |
| I982 | * | * | * | * | 0.7 | * | 0.4 | * | | | | | | | | | | | | | | | | | | TM9 |
| L983 | * | * | * | * | 0.6 | * | 0.4 | * | | | | | | | | | | | | | | | | | | TM9 |
| D984 | * | * | * | * | 0.6 | * | 0.4 | * | | | | | | | | | | | | | | | | | | TM9 |
| | | | | | | | | | H949-D984 | 0.0 | 0.0 | 0.0 | 0.0 | 0.2 | 0.0 | 0.3 | 0.0 | | | | | | | | | |
| | | | | | | | | | R251-D984 | 0.1 | 0.2 | 0.3 | 0.2 | 0.1 | 0.1 | 0.3 | 0.3 | | | | | | | | | |
| | | | | | | | | | R258-D984 | 0.4 | 0.2 | 0.1 | 0.2 | 0.3 | 0.0 | 0.4 | 0.2 | | | | | | | | | |
| | | | | | | | | | R289-D984 | 1.0 | 1.0 | 1.0 | 1.0 | 1.0 | 1.0 | 1.0 | 1.0 | | | | | | | | | |
| | | | | | | | | | S945-D984 | 0.1 | 0.5 | 0.7 | 0.1 | 0.0 | 1.0 | 0.1 | 0.4 | | | | | | | | | |
| | | | | | | | | | T296-D984 | 0.0 | 0.3 | 0.4 | 0.3 | 0.1 | 0.4 | 0.1 | 0.4 | | | | | | | | | |
| D985 | 0.1 | 0.0 | 0.0 | 0.1 | 0.1 | 0.5 | 0.1 | 0.0 | | | | | | | | | | | | | | | | | | TM9 |
| | | | | | | | | | R248-D985 | 1.0 | 0.7 | 0.7 | 0.6 | 0.5 | 0.1 | 0.8 | 0.8 | | | | | | | | | |

| | Alpha helical conformation | | | | | | | | | Electrostatic contact | | | | | | | | Anion contact | | | | | | | | |
|---|---|---|---|---|---|---|---|---|---|---|---|---|---|---|---|---|---|---|---|---|---|---|---|---|---|---|
| **Residue** | **Apo1** | **Apo2** | **Apo3** | **Apo4** | **VX1** | **VX2** | **VX3** | **VX4** | **Contact pair** | **Apo1** | **Apo2** | **Apo3** | **Apo4** | **VX1** | **VX2** | **VX3** | **VX4** | **Apo1** | **Apo2** | **Apo3** | **Apo4** | **VX1** | **VX2** | **VX3** | **VX4** | **TM** |
| | | | | | | | | | R251-D985 | 1.0 | 1.0 | 1.0 | 1.0 | 1.0 | 1.0 | 1.0 | 1.0 | | | | | | | | | |
| | | | | | | | | | R303-D985 | 0.0 | 0.0 | 0.0 | 0.0 | 0.0 | 0.4 | 0.0 | 0.0 | | | | | | | | | |
| **G1003** | * | 0.8 | * | * | * | * | * | * | | | | | | | | | | | | | | | | | | TM9 |
| **A1004** | * | 0.6 | * | * | * | * | * | * | | | | | | | | | | | | | | | | | | TM9 |
| **V1010** | 0.6 | * | 0.7 | 0.5 | 0.6 | 0.8 | 0.8 | 0.4 | | | | | | | | | | | | | | | | | | TM9 |
| **L1043** | * | * | 0.6 | * | * | * | 0.7 | * | | | | | | | | | | | | | | | | | | TM10 |
| **E1046** | 0.7 | * | * | * | * | * | * | * | | | | | | | | | | | | | | | | | | TM10 |
| | | | | | | | | | N974-E1046 | 0.5 | 0.0 | 0.0 | 0.0 | 0.0 | 0.0 | 0.0 | 0.0 | | | | | | | | | |
| | | | | | | | | | Q1042-E1046 | 0.1 | 0.2 | 0.1 | 0.3 | 0.2 | 0.8 | 0.0 | 0.4 | | | | | | | | | |
| | | | | | | | | | R1162-E1046 | 0.8 | 0.4 | 0.0 | 0.6 | 0.3 | 0.9 | 0.1 | 0.8 | | | | | | | | | |
| | | | | | | | | | R975-E1046 | 0.8 | 0.5 | 0.2 | 0.5 | 0.4 | 0.7 | 0.1 | 0.6 | | | | | | | | | |
| | | | | | | | | | S1045-E1046 | 0.0 | 0.0 | 0.3 | 0.1 | 0.1 | 0.0 | 0.2 | 0.1 | | | | | | | | | |
| | | | | | | | | | S1049-E1046 | 0.4 | 0.1 | 0.1 | 0.1 | 0.0 | 0.0 | 0.1 | 0.1 | | | | | | | | | |
| **G1047** | 0.6 | 0.8 | * | 0.7 | 0.8 | 0.7 | * | 0.6 | | | | | | | | | | | | | | | | | | TM10 |
| **R1048** | 0.4 | 0.6 | * | 0.7 | 0.8 | * | * | 0.8 | | | | | | | | | | 0.0 | 0.1 | 0.0 | 0.4 | 0.1 | 0.0 | 0.0 | 0.0 | TM10 |
| | | | | | | | | | R1048-S185 | 0.1 | 0.0 | 0.0 | 0.0 | 0.1 | 0.0 | 0.0 | 0.0 | | | | | | | | | |
| | | | | | | | | | R1048-S977 | 0.0 | 0.0 | 0.0 | 0.0 | 0.0 | 0.1 | 0.0 | 0.0 | | | | | | | | | |
| | | | | | | | | | R1048-D979 | 0.5 | 1.0 | 0.9 | 0.6 | 0.8 | 1.0 | 1.0 | 0.9 | | | | | | | | | |
| | | | | | | | | | R1048-E1044 | 0.7 | 0.1 | 0.5 | 1.0 | 0.5 | 0.4 | 0.5 | 0.1 | | | | | | | | | |
| | | | | | | | | | R1048-S1045 | 0.8 | 0.3 | 0.4 | 0.8 | 0.6 | 1.0 | 0.6 | 0.7 | | | | | | | | | |
| | | | | | | | | | N189-R1048 | 0.9 | 0.1 | 0.0 | 0.6 | 0.5 | 0.0 | 0.0 | 0.1 | | | | | | | | | |
| | | | | | | | | | N974-R1048 | 0.9 | 1.0 | 0.8 | 0.2 | 0.9 | 1.0 | 1.0 | 0.9 | | | | | | | | | |
| | | | | | | | | | R1048-S1159 | 0.1 | 0.2 | 0.0 | 0.1 | 0.2 | 0.0 | 0.1 | 0.5 | | | | | | | | | |
| **P1072** | * | 0.5 | * | * | * | * | 0.4 | * | | | | | | | | | | | | | | | | | | TM11 |
| **Y1073** | * | 0.5 | * | * | * | * | 0.4 | * | | | | | | | | | | | | | | | | | | TM11 |
| | | | | | | | | | Y1073-E504 | 0.2 | 0.3 | 0.2 | 0.3 | 0.2 | 0.3 | 0.3 | 0.3 | | | | | | | | | |
| | | | | | | | | | N505-Y1073 | 0.0 | 0.1 | 0.0 | 0.0 | 0.0 | 0.0 | 0.0 | 0.0 | | | | | | | | | |
| **F1074** | * | 0.5 | * | * | * | * | 0.4 | * | | | | | | | | | | | | | | | | | | TM11 |
| **N1138** | * | * | * | * | 0.7 | 0.8 | 0.8 | * | | | | | | | | | | 0.0 | 0.0 | 0.0 | 0.0 | 0.0 | 0.0 | 0.1 | 0.0 | TM12 |
| | | | | | | | | | Y917-N1138 | 0.0 | 0.0 | 0.5 | 0.3 | 0.0 | 0.0 | 0.6 | 0.6 | | | | | | | | | |
| | | | | | | | | | N1138-S1141 | 0.1 | 0.1 | 0.0 | 0.0 | 0.0 | 0.0 | 0.1 | 0.0 | | | | | | | | | |
| | | | | | | | | | N1138-T1142 | 0.1 | 0.0 | 0.1 | 0.0 | 0.0 | 0.0 | 0.0 | 0.1 | | | | | | | | | |
| **I1139** | * | 0.6 | * | 0.8 | 0.5 | 0.5 | 0.7 | * | | | | | | | | | | | | | | | | | | TM12 |
| **V1160** | * | * | 0.4 | * | * | * | * | * | | | | | | | | | | | | | | | | | | TM12 |
| **S1161** | * | * | 0.7 | * | * | * | * | * | | | | | | | | | | 0.0 | 0.0 | 0.0 | 0.0 | 0.0 | 0.0 | 0.0 | 0.0 | TM12 |
| | | | | | | | | | K1165-S1161 | 0.1 | 0.1 | 0.0 | 0.1 | 0.1 | 0.0 | 0.1 | 0.0 | | | | | | | | | |
| | | | | | | | | | Q1042-S1161 | 0.0 | 0.0 | 0.4 | 0.1 | 0.0 | 0.0 | 0.0 | 0.0 | | | | | | | | | |
| | | | | | | | | | R1158-S1161 | 0.4 | 0.1 | 0.6 | 0.2 | 0.1 | 0.9 | 0.0 | 0.7 | | | | | | | | | |
| | | | | | | | | | T848-S1161 | 0.7 | 0.7 | 0.4 | 0.6 | 0.8 | 0.8 | 0.8 | 0.7 | | | | | | | | | |
| **R1162** | * | * | 0.7 | * | * | * | * | * | | | | | | | | | | 0.0 | 0.0 | 0.0 | 0.2 | 0.2 | 0.0 | 0.1 | 0.0 | TM12 |
| | | | | | | | | | R1162-D979 | 0.8 | 0.0 | 0.0 | 0.0 | 0.1 | 0.9 | 0.2 | 0.1 | | | | | | | | | |
| | | | | | | | | | R1162-E1044 | 0.0 | 0.0 | 0.0 | 0.0 | 0.0 | 0.0 | 0.1 | 0.0 | | | | | | | | | |
| | | | | | | | | | R1162-S1045 | 0.3 | 0.2 | 0.0 | 0.2 | 0.5 | 0.3 | 0.3 | 0.2 | | | | | | | | | |
| | | | | | | | | | R1162-E1046 | 0.8 | 0.4 | 0.0 | 0.6 | 0.3 | 0.9 | 0.1 | 0.8 | | | | | | | | | |
| | | | | | | | | | R1162-S1159 | 0.1 | 0.0 | 0.0 | 0.0 | 0.0 | 0.0 | 0.2 | 0.0 | | | | | | | | | |
| | | | | | | | | | N974-R1162 | 0.8 | 0.0 | 0.0 | 0.0 | 0.0 | 0.9 | 0.0 | 0.1 | | | | | | | | | |
| | | | | | | | | | Q1042-R1162 | 0.3 | 0.7 | 0.5 | 0.7 | 0.5 | 0.9 | 0.4 | 0.8 | | | | | | | | | |
| **I1167** | * | * | 0.7 | * | * | * | * | * | | | | | | | | | | | | | | | | | | TM12 |

**Table S3: Per-residue changes in amino acid / anion / lipid contacts in MD simulations in the presence of VX-770.** This table reports the difference between the total frequency of amino acid/anion/lipid contacts (summed by MD simulation for anion and lipid contacts) for each residue averaged over MD simulations in presence of VX-770 (vx_mean) and MD simulations in apo conditions (apo_mean). This difference (vx_mean − apo_mean) quantifies the shift in contacts upon VX-770 binding. Positive values indicate residues forming more contacts in the VX-770-bound state, while negative values indicate loss of contacts.

**Side-chain/Side chain contacts**

| Increased frequency in presence of VX-770 | | | | Decreased frequency in presence of VX-770 | | | |
|---|---|---|---|---|---|---|---|
| **Contact pair** | **apo_mean** | **vx_mean** | **delta** | **Contact pair** | **apo_mean** | **vx_mean** | **delta** |
| D47-K162 | 0.06 | 0.65 | 0.59 | N1083-S45 | 0.64 | 0.00 | -0.64 |
| H139-T135 | 0.16 | 0.60 | 0.43 | R1358-C1355 | 0.65 | 0.24 | -0.41 |
| N369-Q372 | 0.46 | 0.88 | 0.42 | D44-H147 | 0.44 | 0.04 | -0.40 |
| N1083-Q39 | 0.23 | 0.63 | 0.40 | D47-S45 | 0.69 | 0.30 | -0.40 |
| T1134-Y917 | 0.14 | 0.50 | 0.36 | E282-H954 | 0.86 | 0.47 | -0.39 |
| R74-N71 | 0.61 | 0.96 | 0.35 | H1079-S158 | 0.71 | 0.34 | -0.38 |
| R1097-E193 | 0.07 | 0.40 | 0.33 | S945-Y852 | 0.46 | 0.08 | -0.38 |
| R3-E7 | 0.28 | 0.60 | 0.32 | E54-K162 | 0.40 | 0.04 | -0.36 |
| R153-T1086 | 0.19 | 0.49 | 0.30 | N71-E60 | 0.87 | 0.52 | -0.35 |
| E391-T389 | 0.27 | 0.57 | 0.30 | D44-Q151 | 0.59 | 0.24 | -0.35 |
| D924-T925 | 0.38 | 0.67 | 0.29 | R1128-Y913 | 0.35 | 0.02 | -0.34 |
| S308-Y304 | 0.27 | 0.55 | 0.28 | R1358-Y275 | 0.46 | 0.14 | -0.33 |
| S466-W401 | 0.66 | 0.93 | 0.27 | E1172-K273 | 0.96 | 0.65 | -0.31 |
| E1075-K162 | 0.66 | 0.90 | 0.25 | E543-K968 | 0.71 | 0.40 | -0.31 |
| N965-Q270 | 0.14 | 0.39 | 0.25 | R153-N189 | 0.49 | 0.20 | -0.29 |
| N386-H484 | 0.15 | 0.39 | 0.24 | E1126-S909 | 0.35 | 0.06 | -0.29 |
| R1358-S1362 | 0.41 | 0.66 | 0.24 | W1274-W1282 | 0.31 | 0.03 | -0.28 |
| D110-K114 | 0.25 | 0.48 | 0.23 | R1283-E1172 | 0.92 | 0.64 | -0.28 |
| R334-E116 | 0.19 | 0.42 | 0.23 | D1154-Y849 | 0.74 | 0.46 | -0.28 |
| R1048-N974 | 0.72 | 0.94 | 0.22 | Q372-Q376 | 0.57 | 0.30 | -0.28 |

**Contacts with anions**

| Increased frequency in presence of VX-770 | | | | Decreased frequency in presence of VX-770 | | | |
|---|---|---|---|---|---|---|---|
| Residue | apo_mean | vx_mean | delta | Residue | apo_mean | vx_mean | delta |
| R248 | 0.55 | 0.70 | 0.15 | K1041 | 0.17 | 0.01 | -0.16 |
| R303 | 0.51 | 0.57 | 0.06 | R1048 | 0.14 | 0.02 | -0.12 |
| Q353 | 0.02 | 0.08 | 0.06 | S1159 | 0.11 | 0.00 | -0.11 |
| K95 | 0.09 | 0.14 | 0.05 | R553 | 0.07 | 0.00 | -0.07 |
| Q359 | 0.14 | 0.19 | 0.05 | K978 | 0.14 | 0.07 | -0.06 |
| R352 | 0.16 | 0.20 | 0.04 | Y577 | 0.06 | 0.00 | -0.06 |
| N1148 | 0.02 | 0.03 | 0.02 | K1292 | 0.05 | 0.00 | -0.05 |
| W1145 | 0.02 | 0.03 | 0.01 | N186 | 0.27 | 0.22 | -0.05 |
| R1097 | 0.05 | 0.05 | 0.01 | N189 | 0.07 | 0.02 | -0.05 |
| R251 | 0.14 | 0.15 | 0.01 | S495 | 0.05 | 0.00 | -0.05 |
| R975 | 0.06 | 0.07 | 0.01 | K190 | 0.36 | 0.32 | -0.04 |
| Q98 | 0.01 | 0.01 | 0.00 | S1155 | 0.04 | 0.00 | -0.04 |
| R1162 | 0.06 | 0.06 | 0.00 | Q1042 | 0.04 | 0.01 | -0.03 |
| Q1100 | 0.03 | 0.03 | 0.00 | W356 | 0.27 | 0.25 | -0.03 |
| R1158 | 0.01 | 0.01 | 0.00 | R1070 | 0.03 | 0.00 | -0.03 |
| R334 | 0.01 | 0.01 | 0.00 | K564 | 0.02 | 0.00 | -0.02 |
| S1141 | 0.07 | 0.06 | 0.00 | T1299 | 0.02 | 0.00 | -0.02 |
| S1049 | 0.01 | 0.00 | -0.01 | K1302 | 0.02 | 0.00 | -0.02 |
| N974 | 0.01 | 0.00 | -0.01 | R1301 | 0.02 | 0.00 | -0.02 |
| S1045 | 0.05 | 0.04 | -0.01 | T360 | 0.03 | 0.01 | -0.02 |

**Contacts with POPC**

| Increased frequency in presence of VX-770 | | | | Decreased frequency in presence of VX-770 | | | |
|---|---|---|---|---|---|---|---|
| Residue | apo_mean | vx_mean | delta | Residue | apo_mean | vx_mean | delta |
| R80 | 0.09 | 0.95 | 0.86 | W79 | 0.55 | 0.13 | -0.42 |
| R242 | 0.18 | 0.86 | 0.68 | R104 | 0.60 | 0.21 | -0.39 |
| W1098 | 0.00 | 0.56 | 0.56 | Y247 | 0.47 | 0.17 | -0.30 |
| R1102 | 0.05 | 0.59 | 0.54 | Y903 | 0.27 | 0.01 | -0.26 |
| W361 | 0.00 | 0.50 | 0.50 | K298 | 0.35 | 0.18 | -0.17 |
| S18 | 0.02 | 0.33 | 0.31 | R75 | 0.33 | 0.17 | -0.16 |
| R297 | 0.00 | 0.28 | 0.28 | Q1035 | 0.19 | 0.04 | -0.15 |
| Y84 | 0.00 | 0.25 | 0.25 | S902 | 0.21 | 0.06 | -0.15 |
| K246 | 0.13 | 0.37 | 0.24 | R334 | 0.16 | 0.02 | -0.14 |
| W846 | 0.00 | 0.23 | 0.23 | Y28 | 0.19 | 0.06 | -0.13 |
| R21 | 0.65 | 0.84 | 0.19 | W216 | 0.12 | 0.00 | -0.12 |
| H939 | 0.00 | 0.19 | 0.19 | S912 | 0.11 | 0.00 | -0.11 |
| K14 | 0.09 | 0.22 | 0.14 | S10 | 0.35 | 0.25 | -0.10 |
| R74 | 0.71 | 0.85 | 0.13 | K64 | 0.12 | 0.04 | -0.08 |
| N71 | 0.14 | 0.25 | 0.11 | T338 | 0.07 | 0.00 | -0.07 |
| H147 | 0.00 | 0.09 | 0.09 | R25 | 0.21 | 0.14 | -0.07 |
| R59 | 0.00 | 0.09 | 0.09 | R31 | 0.06 | 0.00 | -0.06 |
| H199 | 0.00 | 0.08 | 0.08 | N886 | 0.08 | 0.02 | -0.06 |
| R29 | 0.15 | 0.23 | 0.08 | Q220 | 0.14 | 0.08 | -0.06 |
| Y301 | 0.00 | 0.07 | 0.07 | K857 | 0.16 | 0.11 | -0.05 |

**Contacts with POPE**

| Increased frequency in presence of VX-770 | | | | Decreased frequency in presence of VX-770 | | | |
|---|---|---|---|---|---|---|---|
| **Residue** | **apo_mean** | **vx_mean** | **delta** | **Residue** | **apo_mean** | **vx_mean** | **delta** |
| N71 | 0.50 | 1.09 | 0.58 | R80 | 0.67 | 0.00 | -0.67 |
| R74 | 0.30 | 0.75 | 0.44 | Y84 | 0.44 | 0.00 | -0.44 |
| R25 | 0.02 | 0.44 | 0.42 | R242 | 0.43 | 0.00 | -0.43 |
| R75 | 0.04 | 0.40 | 0.35 | H147 | 0.41 | 0.00 | -0.41 |
| R104 | 0.29 | 0.60 | 0.31 | N847 | 0.26 | 0.00 | -0.26 |
| R29 | 0.37 | 0.68 | 0.31 | K14 | 0.26 | 0.03 | -0.23 |
| R21 | 0.00 | 0.27 | 0.27 | H939 | 0.28 | 0.06 | -0.23 |
| S118 | 0.00 | 0.26 | 0.26 | S858 | 0.25 | 0.04 | -0.21 |
| Q1035 | 0.00 | 0.23 | 0.23 | K857 | 0.38 | 0.18 | -0.20 |
| R59 | 0.02 | 0.21 | 0.18 | N369 | 0.20 | 0.00 | -0.20 |
| W216 | 0.03 | 0.20 | 0.17 | R297 | 0.23 | 0.04 | -0.20 |
| Y28 | 0.05 | 0.22 | 0.17 | H856 | 0.15 | 0.00 | -0.15 |
| S10 | 0.00 | 0.14 | 0.14 | R899 | 0.20 | 0.08 | -0.12 |
| W846 | 0.12 | 0.25 | 0.13 | Y301 | 0.10 | 0.00 | -0.10 |
| K298 | 0.02 | 0.16 | 0.13 | Y362 | 0.08 | 0.00 | -0.08 |
| N66 | 0.00 | 0.11 | 0.11 | T943 | 0.07 | 0.00 | -0.07 |
| R117 | 0.15 | 0.26 | 0.11 | Y1014 | 0.07 | 0.00 | -0.07 |
| K64 | 0.25 | 0.34 | 0.09 | Q151 | 0.07 | 0.00 | -0.07 |
| R1030 | 0.00 | 0.08 | 0.08 | H897 | 0.10 | 0.04 | -0.06 |
| Y903 | 0.02 | 0.10 | 0.07 | W882 | 0.13 | 0.07 | -0.06 |

**Contacts with POPS**

| Increased frequency in presence of VX-770 | | | | Decreased frequency in presence of VX-770 | | | |
|---|---|---|---|---|---|---|---|
| **Residue** | **apo_mean** | **vx_mean** | **delta** | **Residue** | **apo_mean** | **vx_mean** | **delta** |
| N66 | 0.09 | 0.73 | 0.64 | R21 | 1.89 | 0.61 | -1.28 |
| K857 | 0.12 | 0.58 | 0.47 | R25 | 1.82 | 0.79 | -1.03 |
| K65 | 0.10 | 0.51 | 0.41 | R1102 | 0.64 | 0.07 | -0.57 |
| K68 | 0.48 | 0.88 | 0.39 | S18 | 0.57 | 0.00 | -0.57 |
| R1030 | 0.54 | 0.76 | 0.22 | W846 | 0.54 | 0.00 | -0.54 |
| Y247 | 0.00 | 0.21 | 0.21 | R297 | 1.25 | 0.81 | -0.44 |
| S858 | 0.06 | 0.25 | 0.19 | R29 | 1.21 | 0.80 | -0.41 |
| N369 | 0.09 | 0.28 | 0.19 | W1098 | 0.34 | 0.00 | -0.34 |
| H856 | 0.02 | 0.16 | 0.13 | R80 | 0.32 | 0.00 | -0.32 |
| S1150 | 0.00 | 0.08 | 0.08 | N71 | 0.33 | 0.06 | -0.27 |
| R59 | 0.29 | 0.37 | 0.08 | Q1035 | 0.33 | 0.09 | -0.23 |
| K946 | 0.11 | 0.18 | 0.07 | K246 | 0.73 | 0.51 | -0.22 |
| W79 | 0.00 | 0.07 | 0.07 | K14 | 0.19 | 0.00 | -0.19 |
| Q1038 | 0.13 | 0.19 | 0.06 | R74 | 0.18 | 0.00 | -0.18 |
| R242 | 0.70 | 0.76 | 0.06 | K64 | 0.27 | 0.10 | -0.17 |
| Y849 | 0.00 | 0.05 | 0.05 | K298 | 0.57 | 0.42 | -0.15 |
| T845 | 0.54 | 0.58 | 0.04 | K294 | 1.01 | 0.86 | -0.15 |
| S63 | 0.06 | 0.09 | 0.03 | R31 | 0.14 | 0.00 | -0.14 |
| K370 | 0.00 | 0.03 | 0.03 | K8 | 0.17 | 0.03 | -0.14 |
| W361 | 0.00 | 0.03 | 0.03 | R55 | 0.18 | 0.04 | -0.14 |

##### Contacts with CHL

Increased frequency in presence of VX-770

| Residue | apo_mean | vx_mean | delta |
|---|---|---|---|
| S911 | 0.00 | 0.25 | 0.25 |
| Y849 | 0.00 | 0.16 | 0.16 |
| R80 | 0.00 | 0.12 | 0.12 |
| R74 | 0.00 | 0.11 | 0.11 |
| Y247 | 0.35 | 0.44 | 0.08 |
| K298 | 0.13 | 0.18 | 0.06 |
| R104 | 0.00 | 0.05 | 0.05 |
| S858 | 0.02 | 0.06 | 0.05 |
| W79 | 0.00 | 0.03 | 0.03 |
| S902 | 0.15 | 0.16 | 0.01 |
| R899 | 0.08 | 0.08 | 0.00 |

Decreased frequency in presence of VX-770

| Residue | apo_mean | vx_mean | delta |
|---|---|---|---|
| N901 | 0.29 | 0.00 | -0.29 |
| R242 | 0.32 | 0.04 | -0.28 |
| H939 | 0.38 | 0.15 | -0.23 |
| R21 | 0.22 | 0.00 | -0.22 |
| Y362 | 0.18 | 0.00 | -0.18 |
| R1102 | 0.14 | 0.00 | -0.14 |
| Y304 | 0.09 | 0.00 | -0.09 |
| T887 | 0.08 | 0.00 | -0.08 |
| R297 | 0.07 | 0.00 | -0.07 |
| N900 | 0.06 | 0.00 | -0.06 |
| Y903 | 0.11 | 0.05 | -0.05 |
| T925 | 0.05 | 0.00 | -0.05 |
| T1121 | 0.03 | 0.00 | -0.03 |
| Y301 | 0.04 | 0.02 | -0.02 |
| Y28 | 0.02 | 0.00 | -0.02 |
| W846 | 0.02 | 0.00 | -0.02 |
| Q359 | 0.02 | 0.00 | -0.02 |
| K14 | 0.02 | 0.00 | -0.02 |

##### Contacts with DSM

Increased frequency in presence of VX-770

| Residue | apo_mean | vx_mean | delta |
|---|---|---|---|
| W882 | 0.00 | 0.31 | 0.31 |
| N886 | 0.00 | 0.08 | 0.08 |
| Y913 | 0.13 | 0.21 | 0.08 |
| Q1012 | 0.18 | 0.23 | 0.05 |
| N894 | 0.00 | 0.02 | 0.02 |
| R104 | 0.00 | 0.02 | 0.02 |

Decreased frequency in presence of VX-770

| Residue | apo_mean | vx_mean | delta |
|---|---|---|---|
| N900 | 0.09 | 0.00 | -0.09 |
| R117 | 0.24 | 0.15 | -0.09 |
| Y1014 | 0.08 | 0.00 | -0.08 |
| Q220 | 0.08 | 0.00 | -0.08 |
| R899 | 0.14 | 0.09 | -0.05 |
| S118 | 0.06 | 0.02 | -0.03 |
| Y903 | 0.02 | 0.00 | -0.02 |
| R1128 | 0.32 | 0.31 | -0.01 |

**Table S4: Portals and exits observed at the end of the MD simulations.** The portals/exits are identified regardless of the rotamers of the amino acids lining these structures, except the conditional ones, which are identified only for specific rotamers of one or more amino acids. It should be noted that the ionic flux should be significantly reduced, or even null, depending on the number of amino acids involved, the number of allowed rotamers, and their locations. The diameter of the passage is considered large, medium or small if it is larger, similar, or smaller than a chloride ion, respectively.

| MD | Portal TM4/TM6 | Portal TM10/TM12 | Extracellular exit TM1/TM6 | Extracellular exit ECL1/ECL6 | Lipid associated with R899 (ECL4) |
|---|---|---|---|---|---|
| ***Amino acids involved in the portals*** | *R242, K246, D249-K370, K254* | *R975, R1158, R1162, Q1042, S1045* | | | |
| **Apo1 500 ns** | Large (1 entry)) | Small/conditional (R1162, S1159) | Large/conditional (F337) | Not observed | no, R899 linked to the K329 carbonyl oxygen atom |
| **Apo2 500 ns** | Large (2 entries) | Large | Not observed | Not observed | yes |
| **Apo3 1000 ns** | Large (2 entries) | Large/conditional (Q1042) | Small/conditional (L102) | Large/conditional (R117, linked to a phospholipid) | yes |
| **Apo4 1000 ns** | Large (2 entries) | Large/conditional (K1041, S1159) | Not observed | Medium/conditional (L102, I106, L127, T1134, N1138) | No, R899 linked to the K892 and T896 carbonyl oxygen atoms |
| **VX1 500 ns** | Large (1 entry) | Large/conditional (R1162, S1159) | Small | Not observed | No |
| **VX2 500 ns** | Large (2 entries) | Medium/conditional (E1042, S1045) | Medium | Not observed | No |
| **VX3 1000 ns** | Large (1 entry) | Medium | Not observed | Small/conditional (N113) | No, R899 linked to D891 |
| **VX4 1000 ns** | Large (1 entry) (Conditional, D249) | Not observed | Not observed (closed by F337) | Not observed | No |

**Table S5: Amino acid couples with side chain/side chain contacts and homologous positions in human ABCC4.** The list refers to the amino acids pairs reported in Figures S7 and S8, excluding (i) close neighborhood contacts on the same TMs/coupling helices (CHs), (ii) internal contacts within the NBDs (only inter-NBDs contacts are reported) and (iii) internal contacts within the ECLs (only inter-ECLs contacts are reported). Stable contacts are reported in bold on the interacting partner (CFTR(B)). Amino acid pairs are only listed once in the table, unless each of them is involved in contacts with other amino acids. The sequences of human CFTR (UniPt: P13569) and human ABCC4 (UniPt: O15439) were aligned with MAFFT run on the EMBL/EBI server (Madeira et al. 2025). Identical amino acids are colored green. * in the first column indicates strict conservation of the two amino acids of the pairs between CFTR and ABCC4. To the right of each amino acid in each pair, is indicated CF-causing missense variants or missense variants with varying clinical consequences (VCC, enclosed by parentheses) (legacy nomenclature, CFTR2 database, 15 September 2024, https://cftr2.org/). When available, the refined classification proposed by Veit and colleagues (2016), accounting for complex phenotypes of major CFTR cellular defects, is given to the right of the mutation (classes I to VI)). Lines are colored according to the regions in the CFTR 3D structure, as depicted in Figure 1. Additional columns (right) report the nature of the interaction (S= stable/T= transient), the involvement of the charged or polar amino acids in anion or lipid interaction and the inclusion of amino acids in irregularities relative to the alpha-helical pattern. *ND* indicates lack of obvious homologous amino acids (gaps in the alignment), *NA* stands for *Not Applicable.* Homologous positions in TM7/TM8 should be treated with caution given the structure divergence in this TM hairpin, even though the sequences can be well aligned.

Madeira F, Madhusoodanan N, Lee J, et al. The EMBL-EBI Job Dispatcher sequence analysis tools framework in 2024. Nucleic Acids Research. 2024 Jul;52(W1):W521-W525. DOI: 10.1093/nar/gkae241.

Veit G, Avramescu RG, Chiang AN, Houck SA, Cai Z, Peters KW, Hong JS, Pollard HB, Guggino WB, Balch WE, Skach WR, Cutting GR, Frizzell RA, Sheppard DN, Cyr DM, Sorscher EJ, Brodsky JL, Lukacs GL. From CFTR biology toward combinatorial pharmacotherapy: expanded classification of cystic fibrosis mutations. Mol Biol Cell. 2016 Feb 1;27(3):424-33. doi: 10.1091/mbc.E14-04-0935.

| | CFTR (A) | CF-causing (VCC) | CFTR (B) | CF-causing (VCC) | ABCC4 (A) | ABCC4 (B) | S/T | anion | lipid | Irreg |
|---|---|---|---|---|---|---|---|---|---|---|
| | R3$_{\text{lasso-Nter}}$ | | E7$_{\text{lasso-Nter}}$<br>Q39$_{\text{lasso}}$<br>S42$_{\text{lasso}}$ | | P10 | Q14<br>S46<br>P49 | T<br>T<br>T | -<br>-<br>- | -<br>Yes<br>Yes | *NA* |
| | E7$_{\text{lasso-Nter}}$ | | R3$_{\text{lasso-Nter}}$<br>S42$_{\text{lasso}}$ | | Q14 | P10<br>P49 | T<br>T | -<br>- | Yes<br>Yes | *NA* |
| | K26$_{\text{lh1}}$ | | E33$_{\text{lasso}}$<br>**D36$_{\text{lasso}}$** | | I33 | E40<br>D43 | T<br>**S** | -<br>- | -<br>- | *NA* |
| | R31$_{\text{lasso}}$ | | Q1035$_{\text{TM10}}$<br>Q1038$_{\text{TM10}}$<br>Q1039$_{\text{TM10}}$ | | R38 | E884<br>R887<br>D888 | T<br>T<br>T | -<br>-<br>- | Yes<br>Yes<br>- | *NA* |
| | Q39$_{\text{lasso}}$ | | R3$_{\text{lasso-Nter}}$<br>N1083$_{\text{TM11}}$ | | S46 | P10<br>D932 | T<br>T | -<br>- | -<br>- | *NA* |
| | S42$_{\text{lasso}}$ | | R3$_{\text{lasso-Nter}}$<br>E7$_{\text{lasso-Nter}}$ | | P49 | P10<br>Q14 | T<br>T | -<br>- | Yes<br>Yes | *NA* |
| | D44$_{\text{lasso}}$ | | H147$_{\text{TM2}}$<br>Q151$_{\text{TM2}}$ | | D51 | C161<br>R165 | T<br>T | Yes<br>Yes | Yes<br>Yes | *NA* |
| | S45$_{\text{lasso}}$ | | D47$_{\text{lh2}}$<br>N1083$_{\text{TM11}}$ | | R52 | Q54<br>D932 | T<br>T | -<br>- | -<br>- | *NA* |
| | D47$_{\text{lh2}}$ | | S45$_{\text{lasso}}$<br>H1079$_{\text{TM11}}$<br>N1083$_{\text{TM11}}$ | | Q54 | R52<br>D928<br>D932 | T<br>T<br>T | -<br>-<br>- | -<br>-<br>- | *NA* |
| | S50$_{\text{lh2}}$ | | S158$_{\text{TM2}}$<br>K162$_{\text{TM1}}$<br>H1079$_{\text{TM11}}$ | | G57 | H172<br>R176<br>D928 | T<br>T<br>T | -<br>-<br>- | -<br>-<br>- | *NA* |
| | E54$_{\text{lh2}}$ | | K162$_{\text{CH1}}$<br>K166$_{\text{CH1}}$ | | Q61 | R176<br>R180 | T<br>T | -<br>- | -<br>- | *NA* |
| | E60$_{\text{lh2}}$ | E60K | **K64$_{\text{lasso}}$**<br>**N71$_{\text{elbow1}}$**<br>R74$_{\text{elbow1}}$<br>**R75$_{\text{elbow1}}$** | (R74W)$^{\text{II-III}}$ | E67 | *ND*<br>R82<br>IU85<br>K86 | **S**<br>**S**<br>T<br>**S** | Yes<br>-<br>Yes<br>Yes | Yes<br>Yes<br>Yes<br>Yes | *NA* |
| | K64$_{\text{lasso}}$ | | **E60$_{\text{lh2}}$**<br>N71$_{\text{elbow1}}$ | E60K | A71 | E67<br>R82 | **S**<br>T | Yes<br>- | Yes<br>Yes | *NA* |
| | N66$_{\text{lasso}}$ | | Q372$_{\text{TM6}}$<br>Q376$_{\text{TM6}}$ | | K77 | Q387<br>L391 | T<br>T | -<br>- | Yes<br>Yes | *NA* |
| | N71$_{\text{elbow1}}$ | | **E60$_{\text{lh2}}$**<br>K64$_{\text{lasso}}$ | E60K | R82 | E67<br>A71 | **S**<br>T | -<br>- | Yes<br>Yes | *NA* |
| | R75$_{\text{elbow1}}$ | | **E56$_{\text{lh2}}$**<br>**E60$_{\text{lh2}}$**<br>S63$_{\text{lh2}}$ | E56K$^{\text{II-III}}$<br>E60K | K87 | F63<br>E67<br>R70 | **S**<br>**S**<br>T | Yes<br>Yes<br>Yes | Yes<br>Yes<br>Yes | *NA* |
| *<br>* | E92$_{\text{TM1}}$ | E92K$^{\text{II-III}}$ | **K95$_{\text{TM1}}$**<br>Q207$_{\text{TM3}}$<br>Q353$_{\text{TM6}}$ | | E103 | K106<br>Q221<br>F368 | **S**<br>T<br>T | Yes<br>-<br>Yes | -<br>-<br>- | - |
| | Y109$_{\text{TM1}}$ | | S1118$_{\text{TM11}}$<br>T1134$_{\text{TM12}}$ | S1118F | F120 | S967<br>S984 | T<br>T | -<br>- | -<br>- | - |
| | D110$_{\text{ECL1}}$ | D110H$^{\text{II-III}}$<br>(D110E)$^{\text{-III}}$ | R334$_{\text{ECL3}}$<br><br>T910$_{\text{ECL4}}$<br>Y914$_{\text{TM8}}$ | | D124 | S349<br><br>L759<br>W763 | T<br><br>T<br>T | Yes<br><br>-<br>- | Yes<br><br>-<br>- | *NA* |
| | R117$_{\text{ECL1}}$ | (R117H)$^{\text{II-III}}$<br>(R117L)<br>(R117G)$^{\text{II-III}}$<br>R117P<br>R117C | T1122$_{\text{ECL6}}$ | | L131 | K972 | T | - | Yes | *NA* |
| | C128$_{\text{TM2}}$ | | T1115$_{\text{TM11}}$ | | T142 | A964 | T | - | - | - |
| | R134$_{\text{TM2}}$ | | Q98$_{\text{TM1}}$<br>E1104$_{\text{TM11}}$ | Q98R | L148 | Q109<br>D953 | T<br>T | Yes<br>Yes | -<br>- | - |
| | H139$_{\text{TM2}}$ | H139R | Q1100$_{\text{TM11}}$<br>E1104$_{\text{TM11}}$ | | H153 | A949<br>D953 | T<br>T | -<br>Yes | -<br>- | - |
| | H147$_{\text{TM2}}$ | | D44$_{\text{lasso}}$<br>Y84$_{\text{TM1}}$ | | C616 | D51<br>L95 | T<br>T | Yes<br>Yes | Yes<br>Yes | - |

|  | CFTR (A) | CF-causing (VCC) | CFTR (B) | CF-causing (VCC) | ABCC4 (A) | ABCC4 (B) | S/T | anion | lipid | Irreg |
|---|---|---|---|---|---|---|---|---|---|---|
| | Q151$_{TM2}$ | | D44$_{lasso}$ | | R165 | D51 | T | Yes | Yes | - |
| | | | R80$_{TM1}$ | | | S91 | T | Yes | Yes | |
| | R153$_{TM2}$ | | N187$_{TM3}$ | | R167 | D201 | T | - | - | - |
| * | | | N189$_{TM3}$ | | | N203 | T | Yes | - | |
| * | | | **D192$_{TM3}$** | D192G | | D206 | **S** | - | - | |
| * | | | E1044$_{TM10}$ | | | E893 | T | - | - | |
| | | | T1086$_{TM11}$ | | | S935 | T | - | - | |
| | S158$_{TM2}$ | | S50$_{lh2}$ | | H172 | G57 | T | - | - | - |
| | | | H1079$_{TM11}$ | | | D928 | T | - | - | |
| * | Y161$_{TM2}$ | Y161D | **S1058$_{TM10}$** | | Y175 | S907 | **S** | - | - | - |
| * | | | H1054$_{TM10}$ | H1054D $^{II-III}$ | | H903 | T | - | - | |
| | K162$_{TM2}$ | | S50$_{lh2}$ | | R177 | G57 | T | - | - | - |
| | | | E54$_{lh2}$ | | | Q61 | T | - | - | |
| | | | E1075$_{TM11}$ | | | Q924 | T | - | - | |
| | K166$_{CH1}$ | | E54$_{lh2}$ | | R181 | Q61 | T | - | - | - |
| | | | E379$_{TMD1-NBD1}$ | | | E394 | T | - | - | |
| | S169$_{CH1}$ | | **E474$_{NBD1}$** | E474K | N183 | E461 | **S** | - | - | - |
| | | | W1063$_{CH4}$ | | | W912 | T | - | - | |
| | | | R1066$_{CH4}$ | R1066C $^{II}$<br>R1066H $^{II-III}$ | | R915 | T | - | - | |
| | R170$_{CH1}$ | | E402$_{NBD1}$ | | M184 | D420 | T | Yes | - | - |
| | | | E403$_{NBD1}$ | | | *ND* | T | Yes | - | |
| | | | E476$_{NBD1}$ | | | A463 | T | Yes | - | |
| | D173$_{CH1}$ | | W401$_{NBD1}$ | | G187 | W419 | T | - | - | - |
| | | | C1344$_{NBD2}$ | | | S1175 | T | - | - | |
| | K174$_{CH1}$ | | E402$_{NBD1}$ | | K188 | D420 | T | - | - | - |
| | | | E403$_{NBD1}$ | | | *ND* | T | - | - | |
| | | | D1341$_{NBD2}$ | | | E1172 | T | - | - | |
| | S182$_{TM3}$ | | N186$_{TM3}$ | | N196 | N200 | T | Yes | - | - |
| | | | S256$_{TM4}$ | | | D270 | T | - | - | |
| * | S185$_{TM3}$ | | N974$_{TM9}$ | | S199 | N823 | T | Yes | - | - |
| * | | | S977$_{TM9}$ | (S977F) $^{II-III}$ | | S826 | T | - | - | |
| * | N189$_{TM3}$ | | R153$_{TM2}$ | | N203 | R167 | T | Yes | - | Yes |
| * | | | K978$_{TM9}$ | | | K827 | T | Yes | - | |
| * | | | K1041$_{TM10}$ | | | K890 | T | Yes | - | |
| * | | | E1044$_{TM10}$ | | | E893 | T | Yes | - | |
| * | | | R1048$_{TM10}$ | H1085P | | R897 | T | Yes | - | |
| * | | | H1085$_{TM11}$ | | | H930 | T | Yes | - | |
| * | | | W1089$_{TM11}$ | | | W938 | T | Yes | - | |
| * | D192$_{TM3}$ | D192G | **R153$_{TM2}$** | | D206 | R167 | **S** | - | - | Yes |
| * | | | **W1089$_{TM11}$** | | | W938 | **S** | - | - | |
| | Q207$_{TM3}$ | | Y89$_{TM1}$ | | Q221 | L100 | T | - | - | - |
| * | | | E92$_{TM1}$ | G92K $^{II-III}$ | | E103 | T | - | - | |
| | Q220$_{ECL2}$ | | Y325$_{ECL3}$ | | G234 | Y339 | T | - | Yes | *NA* |
| | | | K329$_{TM5}$ | | | G353 | T | - | Yes | |
| * | Q237$_{TM4}$ | (Q237E) | **N306$_{TM5}$** | | Q251 | N320 | **S** | Yes | - | - |
| | | | S313$_{TM5}$ | | | A327 | T | - | - | |
| | | | **N396$_{NBD1}$** | | | D414 | **S** | Yes | - | |
| | R242$_{TM4}$ | | Y362$_{TM6}$ | | K256 | S377 | T | - | Yes | - |
| | R248$_{TM4}$ | | Q359$_{TM6}$ | Q359R | R262 | E374 | T | Yes | Yes | - |
| | | | D363$_{TM6}$ | | | E376 | T | Yes | - | |
| * | | | D985$_{TM9}$ | | | D834 | T | Yes | - | |
| | R251$_{TM4}$ | | **T296$_{TM5}$** | | T265 | I310 | **S** | Yes | - | Yes |
| | | | D984$_{TM9}$ | | | D833 | T | Yes | - | |
| | | | **D985$_{TM9}$** | | | D834 | **S** | Yes | - | |
| | S256$_{TM4}$ | | S176$_{CH1}$ | | D270 | T190 | T | - | - | - |
| | | | Q179$_{TM3}$ | | | Q193 | T | - | - | |
| | | | S182$_{TM3}$ | | | N196 | T | - | - | |
| * | R258$_{TM4}$ | (R258G) | **E292$_{TM5}$** | | R272 | E306 | **S** | Yes | - | - |
| * | | | D984$_{TM9}$ | | | D833 | T | - | - | |

| | CFTR (A) | CF-causing (VCC) | CFTR (B) | CF-causing (VCC) | ABCC4 (A) | ABCC4 (B) | S/T | anion | lipid | Irreg |
|---|---|---|---|---|---|---|---|---|---|---|
| | T262$_{TM4}$ | | H949$_{TM8}$ | | M276 | H798 | T | - | - | - |
| | N268$_{CH2}$ | | K1292$_{NBD2}$ | | G282 | E1123 | T | - | - | *NA* |
| | Q270$_{CH2}$ | | D965$_{CH3}$<br>S1255$_{NBD2}$ | S1255P [III] | R284 | D834<br>S1086 | T<br>T | -<br>- | -<br>- | *NA* |
| <br>* | K273$_{CH2}$ | | Q958$_{CH3}$<br>**E1172**$_{\mathbf{TMD2\text{-}NBD2}}$ | | K287 | K807<br>E1022 | T<br>**S** | -<br>- | -<br>- | *NA* |
| *<br>* | Y275$_{CH2}$ | | **C1355**$_{\mathbf{NBD2}}$<br>R1358$_{NBD2}$ | | Y289 | C1186<br>R1189 | **S**<br>T | -<br>- | -<br>- | *NA* |
| <br><br>* | E278$_{TM5}$ | | Q958$_{CH3}$<br>Q1280$_{NBD2}$<br>R1283$_{NBD2}$ | R1283M | E291 | K807<br>H1111<br>R1104 | T<br>T<br>T | -<br>-<br>- | -<br>-<br>- | - |
| | E279$_{TM5}$ | | Y1307$_{NBD2}$ | | K292 | F1138 | T | - | - | - |
| | E282$_{TM5}$ | | H950$_{TM9}$<br>**H954**$_{\mathbf{TM9}}$<br>Q958$_{CH3}$ | | S294 | N799<br>G803<br>K807 | T<br>**S**<br>T | -<br>-<br>- | -<br>-<br>- | - |
| *<br>*<br>* | R289$_{TM5}$ | | **E292**$_{\mathbf{TM5}}$<br>**D984**$_{\mathbf{TM9}}$<br>D985$_{TM9}$ | | R303 | E306<br>D833<br>D834 | **S**<br>**S**<br>T | -<br>-<br>- | -<br>-<br>- | - |
| | Q290$_{TM5}$ | | K946$_{TM8}$<br>H950$_{TM8}$ | | K304 | Q795<br>N799 | T<br>T | Yes<br>Yes | Yes<br>Yes | - |
| *<br>* | E292$_{TM5}$ | | **R258**$_{\mathbf{TM4}}$<br>**R289**$_{\mathbf{TM5}}$ | (R258G) | E306 | R272<br>R303 | **S**<br>**S** | Yes<br>- | -<br>- | - |
| | R297$_{TM5}$ | | T943$_{TM8}$ | | L311 | N792 | T | Yes | Yes | - |
| | Y301$_{TM5}$ | | **H939**$_{\mathbf{TM8}}$ | | C315 | Y788 | **S** | - | Yes | - |
| <br><br>*<br>* | R303$_{TM5}$ | | S307$_{TM5}$<br>Q359$_{TM6}$<br>T990$_{TM9}$<br>D993$_{TM9}$ | Q359R | R317 | L231<br>E374<br>T839<br>D842 | T<br>T<br>T<br>T | Yes<br>Yes<br>Yes<br>Yes | -<br>Yes<br>-<br>- | - |
| * | N306$_{TM5}$ | | **Q359**$_{\mathbf{TM6}}$<br>**Q237**$_{\mathbf{TM4}}$ | Q359R<br>(Q237E) | N320 | Q251<br>E374 | **S**<br>**S** | Yes<br>Yes | -<br>Yes | - |
| | S307$_{TM5}$ | | R303$_{TM5}$<br>R352$_{TM6}$<br><br>D993$_{TM9}$ | R352Q [III]<br>(R352W) | L321 | R317<br>L367<br><br>D842 | T<br>T<br><br>T | Yes<br>Yes<br><br>- | -<br>-<br><br>- | - |
| | S313$_{TM5}$ | | Q237$_{TM4}$<br>R347$_{TM6}$<br><br>**T351**$_{\mathbf{TM6}}$ | (Q237E)<br>R347P [II-III-VI]<br>R347H [III] | A327 | Q251<br>R362<br><br>T366 | T<br>T<br><br>**S** | -<br>-<br><br>- | -<br>-<br><br>- | Yes |
| | S321$_{TM5}$ | | **C343**$_{\mathbf{TM6}}$ | | T335 | Y358 | **S** | - | - | - |
| | R334$_{ECL3}$ | R334W [II-III]<br>R334L<br>(R334Q) | D110$_{ECL1}$<br>D112$_{ECL1}$<br>E116$_{ECL1}$<br>Y914$_{TM8}$ | E116K | S349 | D124<br>M126<br>A130<br>W763 | T<br>T<br>T<br>T | Yes<br>Yes<br>Yes<br>- | Yes<br>Yes<br>Yes<br>- | *NA* |
| | R347$_{TM6}$ | R347P<br>R347H | S313$_{TM5}$<br>**D924**$_{\mathbf{TM8}}$<br>T925$_{TM8}$ | | R362 | A327<br>V773<br>A774 | T<br>**S**<br>T | -<br>-<br>- | -<br>-<br>Yes | - |
| | R352$_{TM6}$ | R352Q<br>(R352W) | D993$_{TM9}$<br>S307$_{TM5}$ | | L367 | D842<br>L321 | T<br>T | Yes<br>Yes | -<br>- | Yes |
| | Q359$_{TM6}$ | Q359R | R248$_{TM4}$<br>R303$_{TM5}$<br>**N306**$_{\mathbf{TM5}}$ | | E374 | R262<br>R317<br>N320 | T<br>T<br>**S** | Yes<br>Yes<br>Yes | Yes<br>Yes<br>Yes | - |
| | D363$_{TM6}$ | | K190$_{TM3}$<br>R248$_{TM4}$ | | E378 | K204<br>R262 | T<br>T | Yes<br>Yes | -<br>- | - |
| | K370$_{TM6}$ | | D249$_{TM4}$ | | R385 | S263 | T | Yes | - | - |
| | K377$_{TMD1\text{-}NBD1}$ | | E407$_{NBD1}$ | | L392 | *ND* | T | - | - | *NA* |
| * | E379$_{TMD1\text{-}NBD1}$ | | **K163**$_{\mathbf{TM2}}$<br>K166$_{CH1}$ | | E394 | K178<br>R180 | **S**<br>T | -<br>- | -<br>- | *NA* |

| | CFTR (A) | CF-causing (VCC) | CFTR (B) | CF-causing (VCC) | ABCC4 (A) | ABCC4 (B) | S/T | anion | lipid | Irreg |
|---|---|---|---|---|---|---|---|---|---|---|
| | E402$_{NBD1}$ | | R170$_{CH1}$<br>K174$_{CH1}$ | | D420 | M184<br>K188 | T<br>T | Yes<br>- | -<br>- | *NA* |
| | E403$_{NBD1}$ | | R170$_{CH1}$<br>K174$_{CH1}$ | | *ND* | M184<br>K188 | T<br>T | Yes<br>- | -<br>- | *NA* |
| | S459$_{NBD1}$ | | D1377$_{NBD2}$<br>T1380$_{NBD2}$ | | P446 | D1208<br>T1211 | T<br>T | -<br>- | -<br>- | *NA* |
| | T460$_{NBD1}$ | | S1347$_{NBD2}$<br>H1348$_{NBD2}$<br>H1350$_{NBD2}$<br>H1375$_{NBD2}$ | H1375P | V446 | S1178<br>V1179<br>Q1181<br>N1206 | T<br>T<br>T<br>T | -<br>-<br>-<br>- | -<br>-<br>-<br>- | *NA* |
| <br>*<br>* | E474$_{NBD1}$ | E474K | S169$_{CH1}$<br>W1063$_{CH4}$<br>**R1066**$_{CH4}$ | | E461 | N183<br>W912<br>R915 | **S**<br>T<br>**S** | -<br>-<br>- | -<br>-<br>- | *NA* |
| | E476$_{NBD1}$ | | K381$_{TMD1-NBD1}$<br>R170$_{CH1}$ | | S465 | S396<br>M184 | T<br>T | -<br>- | -<br>- | *NA* |
| | Q493$_{NBD1}$ | | H1348$_{NBD2}$<br>H1375$_{NBD2}$ | H1375P | Q480 | V1179<br>N1206 | T<br>T | -<br>- | -<br>- | *NA* |
| | E543$_{NBD1}$ | | K968$_{CH3}$<br>T1053$_{TM10}$<br>T1057$_{TM10}$ | | D530 | P817<br>S902<br>S906 | T<br>T<br>T | -<br>-<br>- | -<br>-<br>- | *NA* |
| | Y577$_{NBD1}$ | | Q1291$_{NBD2}$<br>H1375$_{NBD2}$ | (Q1291H)<br>(Q1291R)<br>H1375P | A564 | Q1122<br>N1206 | T<br>T | Yes<br>Yes | -<br>- | *NA* |
| | E583$_{NBD1}$ | | R1403$_{NBD2}$ | | S570 | R1234 | T | Yes | - | *NA* |
| | T845$_{elbow2}$ | | R1030$_{TM10}$<br>D1154$_{TM12}$ | | G697 | R879<br>E1004 | T<br>T | -<br>- | Yes<br>Yes | *NA* |
| | | | R1158$_{TM12}$ | | | I1008 | T | Yes | Yes | |
| | T848$_{elbow2}$ | | S1161$_{TM12}$<br>R1158$_{TM12}$ | | A700 | E1011<br>I1008 | T<br>T | Yes<br>Yes | -<br>Yes | *NA* |
| * | R851$_{elbow2}$ | | **D1168**$_{TMD2-NBD2}$ | | R706 | D1118 | **S** | - | Yes | *NA* |
| | S909$_{ECL4}$ | | E1126$_{ECL6}$<br>R1128$_{ECL6}$ | | K758 | A976<br>Q978 | T<br>T | -<br>- | -<br>Yes | *NA* |
| | T910$_{ECL4}$ | | D110$_{ECL1}$<br>E1126$_{ECL6}$ | D110H$^{II-III}$<br>(D110E)$^{III}$ | L759 | D124<br>A976 | T<br>T | -<br>- | -<br>- | *NA* |
| | Y913$_{TM8}$ | | Q1012$_{ECL5}$<br>E1126$_{ECL6}$<br>R1128$_{ECL6}$ | | N762 | I861<br>A976<br>Q978 | T<br>T<br>T | -<br>-<br>- | Yes<br>-<br>- | *NA* |
| | Y914$_{TM8}$ | | D110$_{ECL1}$<br>R334$_{TM6}$ | D110H$^{II-III}$<br>(D110E)$^{III}$<br>R334W$^{II-III}$<br>R334L<br>(R334Q) | W763 | D124<br>S349 | T<br>T | - | - | *NA* |
| | Y917$_{TM8}$ | | S341$_{TM6}$<br>T1134$_{TM12}$<br>N1138$_{TM12}$ | S341P | G766 | T356<br>S984<br>T988 | T<br>T<br>T | -<br>-<br>Yes | -<br>-<br>- | - |
| | Y919$_{TM8}$ | | **S877**$_{TM7}$ | | Y768 | V729 | **S** | - | - | - |
| | D924$_{TM8}$ | (D924N) | **R347**$_{TM6}$ | R347P$^{II-III-VI}$<br>R347H$^{III}$ | V773 | R362 | **S** | - | - | - |
| | R933$_{TM8}$ | | **E873**$_{TM7}$<br>C866$_{TM7}$ | | R782 | Q725<br>I718 | **S**<br>T | -<br>- | -<br>- | Yes |
| | H939$_{TM8}$ | | **Y301**$_{TM5}$ | | Y788 | C315 | **S** | - | Yes | - |
| *<br>* | S945$_{TM8}$ | S945L$^{II-III}$ | Y852$_{elbow2}$<br>D984$_{TM9}$ | | S794 | Y704<br>D833 | T<br>T | -<br>Yes | -<br>- | - |
| | H950$_{TM8}$ | | E282$_{TM5}$<br>E286$_{TM5}$<br>Q290$_{TM5}$ | | N799 | E306<br>T300<br>K304 | T<br>T<br>T | -<br>-<br>Yes | -<br>-<br>Yes | - |

| | CFTR (A) | CF-causing (VCC) | CFTR (B) | CF-causing (VCC) | ABCC4 (A) | ABCC4 (B) | S/T | anion | lipid | Irreg |
|---|---|---|---|---|---|---|---|---|---|---|
| | Q958$_{CH3}$ | | K273$_{CH2}$ | | K807 | K287 | T | - | - | *NA* |
| | | | E278$_{TM5}$ | | | E291 | T | - | - | |
| | | | E282$_{TM5}$ | | | S294 | T | - | - | |
| | | | T1171$_{TM11}$ | | | K1021 | T | - | - | |
| | | | Q1280$_{NBD2}$ | | | H1111 | T | - | - | |
| | K968$_{CH3}$ | | E543$_{NBD1}$ | | P817 | D530 | T | - | - | *NA* |
| | | | Q1280$_{NBD2}$ | | | H1111 | T | - | - | |
| * | N974$_{TM9}$ | | S185$_{TM3}$ | | N823 | S199 | T | Yes | - | - |
| * | | | R975$_{TM9}$ | | | R824 | T | Yes | - | |
| * | | | S977$_{TM9}$ | (S977F) $^{II-III}$ | | S826 | T | Yes | - | |
| * | | | D979$_{TM9}$ | D979V | | D828 | T | Yes | - | |
| | | | E1046$_{TM10}$ | | | T895 | T | Yes | - | |
| * | | | R1048$_{TM10}$ | | | R897 | T | Yes | - | |
| * | | | S1049$_{TM10}$ | | | S898 | T | Yes | - | |
| * | | | R1162$_{TM12}$ | | | R1012 | T | Yes | - | |
| * | R975$_{TM9}$ | | N974$_{TM9}$ | | R824 | N823 | T | Yes | - | - |
| * | | | D979$_{TM9}$ | D979V | | D828 | T | Yes | - | |
| * | | | S1045 $_{TM10}$ | | | S894 | T | Yes | - | |
| | | | E1046 $_{TM10}$ | | | T895 | T | Yes | - | |
| * | | | S1049 $_{TM10}$ | | | S898 | T | Yes | | |
| * | D979$_{TM9}$ | D979V | N974$_{TM9}$ | | D828 | N823 | T | Yes | - | Yes |
| * | | | R975$_{TM9}$ | | | R824 | T | Yes | - | |
| * | | | R1048$_{TM10}$ | | | R897 | T | Yes | - | |
| * | | | S1159$_{TM12}$ | S1159F | | S1009 | T | Yes | - | |
| | | | | S1159P | | | | | - | |
| * | | | R1162$_{TM12}$ | | | R1012 | T | Yes | | |
| * | D984$_{TM9}$ | | R248$_{TM4}$ | | D833 | R262 | T | - | - | Yes |
| | | | R251$_{TM4}$ | | | T265 | T | Yes | - | |
| * | | | R258$_{TM4}$ | | | R272 | T | - | - | |
| * | | | **R289$_{TM5}$** | | | R303 | **S** | - | - | |
| | | | T296$_{TM5}$ | | | I310 | T | - | - | |
| * | | | S945$_{TM8}$ | S945L $^{II-III}$ | | S794 | T | Yes | - | |
| * | D985$_{TM9}$ | | R248$_{TM4}$ | | D834 | R262 | T | Yes | - | Yes |
| * | | | **R251$_{TM4}$** | | | R265 | **S** | Yes | - | - |
| | | | Q270$_{CH2}$ | | | R284 | T | - | - | |
| * | D993$_{TM9}$ | | R303$_{TM5}$ | | D842 | R317 | T | Yes | - | - |
| | | | S307$_{TM5}$ | | | L321 | T | - | - | |
| | | | R352$_{TM6}$ | R352Q $^{III}$ | | L367 | T | Yes | - | |
| | | | | (R352W) | | | | | | |
| | R1030$_{TM10}$ | | T845$_{elbow2}$ | | R879 | G697 | T | - | Yes | - |
| * | | | S1150$_{TM12}$ | | | S1000 | T | - | Yes | |
| | | | **D1154$_{TM12}$** | | | E1004 | **S** | - | Yes | |
| | Y1032$_{TM10}$ | (Y1032C) | R21$_{lh1}$ | | Y881 | N28 | T | - | Yes | - |
| | | | R25$_{lh1}$ | | | K32 | T | - | Yes | |
| | S1037$_{TM10}$ | | S1155$_{TM12}$ | | S886 | N1005 | T | Yes | - | - |
| | | | Y1092$_{TM11}$ | | | F941 | T | - | - | |
| | Q1038$_{TM10}$ | | R31$_{lasso}$ | | R887 | R38 | T | - | Yes | - |
| | | | T845$_{elbow2}$ | | | G697 | T | - | Yes | |
| | | | **R1158$_{TM12}$** | | | I1008 | **S** | - | Yes | |
| | Q1039$_{TM10}$ | | R31$_{lasso}$ | | D888 | R38 | T | - | - | - |
| | | | R1158$_{TM12}$ | | | I1008 | T | - | - | |
| * | K1041$_{TM10}$ | | N189$_{TM3}$ | | K890 | N203 | T | Yes | - | - |
| | | | S1155$_{TM12}$ | | | N1005 | T | Yes | - | |
| * | | | S1159$_{TM12}$ | S1159F | | S1009 | T | Yes | - | |
| | | | | S1159P | | | | | | |
| | Q1042$_{TM10}$ | | R1162 $_{TM12}$ | | R891 | R1012 | T | Yes | - | - |
| | | | R1158$_{TM12}$ | | | I1008 | T | Yes | - | |
| | | | S1161$_{TM12}$ | | | E1011 | T | Yes | - | |

| | CFTR (A) | CF-causing (VCC) | CFTR (B) | CF-causing (VCC) | ABCC4 (A) | ABCC4 (B) | S/T | anion | lipid | Irreg |
|---|---|---|---|---|---|---|---|---|---|---|
| *<br>*<br>*<br>* | E1044$_{TM10}$ | | R153$_{TM2}$<br>N189$_{TM3}$<br>H1085$_{TM11}$<br>S1159$_{TM12}$ | H1085P<br>S1159F<br>S1159P | E893 | R167<br>N203<br>H934<br>S1009 | T<br>T<br>T<br>T | -<br>Yes<br>-<br>Yes | -<br>-<br>-<br>- | - |
| *<br>* | S1045$_{TM10}$ | | R975$_{TM9}$<br>R1162$_{TM12}$ | | S894 | R824<br>R1012 | T<br>T | Yes<br>Yes | -<br>- | - |
| | E1046$_{TM10}$ | | N974$_{TM9}$<br>R975$_{TM9}$<br>R1162$_{TM12}$ | | T895 | N823<br>R824<br>R1012 | T<br>T<br>T | Yes<br>Yes<br>Yes | -<br>- | Yes |
| *<br>*<br>*<br>* | R1048$_{TM10}$ | | N189$_{TM3}$<br>N974$_{TM9}$<br>D979$_{TM9}$<br>S1159$_{TM12}$ | D979V<br>S1159F<br>S1159P | R897 | N203<br>N823<br>D828<br>S1009 | T<br>T<br>T<br>T | Yes<br>Yes<br>Yes<br>Yes | -<br>-<br><br>- | Yes |
| *<br>*<br>* | S1049$_{TM10}$ | | N974$_{TM9}$<br>R975$_{TM9}$<br>D979$_{TM9}$ | | S898 | N823<br>R824<br>D828 | T<br>T<br>T | Yes<br>Yes<br>Yes | -<br>-<br>- | - |
| | T1057$_{TM10}$ | | E543$_{NBD1}$<br>W496$_{NBD1}$ | | S906 | D530<br>W483 | T<br>T | -<br>- | -<br>- | - |
| | K1060$_{TM10}$ | | E267$_{TM4}$<br>D1341$_{NBD2}$ | | Q909 | T281<br>E1172 | T<br>T | Yes<br>Yes | -<br>- | - |
| <br>* | W1063$_{CH4}$ | | S169$_{CH1}$<br>E474 $_{NBD1}$ | <br>E474K | W912 | N183<br>E461 | T<br>T | -<br>- | -<br>- | *NA* |
| *<br><br><br>* | T1064$_{CH4}$ | | R560$_{NBD1}$<br><br><br>S492 $_{NBD1}$ | R560T$^{II}$<br>R560K$^{II}$<br>R560S$^{II}$<br>S492F$^{II-VI}$ | T913 | R547<br><br><br>S479 | T<br><br><br>T | -<br><br><br>- | -<br><br><br>- | *NA* |
| <br>* | R1066$_{CH4}$ | R1066C$^{II}$<br>R1066H$^{II-III}$ | S169$_{CH1}$<br>**E474$_{NBD1}$** | <br>E474K | R915 | N182<br>E461 | T<br>**S** | -<br>- | -<br>- | *NA* |
| | Q1071$_{TM11}$ | | Y380$_{TMD1-NBD1}$ | | E920 | I395 | T | - | - | - |
| | Y1073$_{TM11}$ | | E504$_{NBD1}$ | | R922 | S491 | T | - | - | - |
| | H1079$_{TM11}$ | | D47$_{lh2}$<br>S50$_{lh2}$<br>S158$_{TM2}$ | | D928 | Q54<br>G57<br>H172 | T<br>T<br>T | -<br>-<br>- | -<br>-<br>- | - |
| | N1083$_{TM11}$ | | Q39$_{lasso}$<br>S45$_{lasso}$<br>D47$_{lh2}$ | | D932 | S46<br>R52<br>Q54 | T<br>T<br>T | -<br>-<br>- | -<br>-<br>- | - |
| *<br>* | H1085$_{TM11}$ | H1085P | N189$_{TM3}$<br>E1044$_{TM10}$ | | H934 | N203<br>E893 | T<br>T | Yes<br>- | -<br>- | - |
| *<br>* | W1089$_{TM11}$ | | N189$_{TM3}$<br>**D192$_{TM3}$** | <br>D192G | W938 | N203<br>D206 | T<br>**S** | Yes<br>- | -<br>- | - |
| | Y1092$_{TM11}$ | | S1037$_{TM10}$<br>D1152$_{TM12}$<br>S1155$_{TM12}$ | (D1152H)$^{III}$ | F941 | S886<br>E1002<br>N1005 | T<br>T<br>**S** | -<br>-<br>- | -<br>-<br>- | - |
| | W1098$_{TM11}$ | W1098R<br>W1098C | T20$_{lh1}$ | | W947 | D65 | T | - | Yes | - |
| | Q1100$_{TM11}$ | | H139$_{TM2}$<br>**N1148$_{TM12}$** | | A949 | H153<br>R998 | T<br>**S** | Yes<br>Yes | -<br>- | - |
| | R1102$_{TM11}$ | | S18$_{lasso}$ | | R951 | W63 | T | - | Yes | - |
| | E1104$_{TM11}$ | | R134$_{TM2}$<br>T135$_{TM2}$<br>H139$_{TM2}$<br>Q1144$_{TM12}$ | | D953 | L148<br>A149<br>H153<br>Q994 | T<br>T<br>T<br>T | Yes<br>-<br>-<br>Yes | -<br>-<br>-<br>- | - |
| | S1118$_{TM11}$ | S1118F | Y109$_{ECL1}$<br>T1134$_{TM12}$ | | S967 | F120<br>S984 | T<br>T | -<br>- | -<br>- | - |
| | E1126$_{ECL6}$ | | S909$_{ECL4}$<br>T910 $_{ECL4}$<br>Y913 $_{TM8}$ | | A976 | K758<br>L759<br>N762 | T<br>T<br>T | -<br>-<br>- | -<br>-<br>Yes | *NA* |
| | R1128$_{ECL6}$ | | S909$_{ECL4}$<br>Y913$_{TM8}$ | | Q978 | K758<br>N762 | T<br>T | -<br>- | Yes<br>Yes | *NA* |

| | CFTR (A) | CF-causing (VCC) | CFTR (B) | CF-causing (VCC) | ABCC4 (A) | ABCC4 (B) | S/T | anion | lipid | Irreg |
|---|---|---|---|---|---|---|---|---|---|---|
| | T1134$_{TM12}$ | | Y109$_{ECL1}$<br>Y917$_{TM8}$<br>S1118$_{TM11}$ | S1118F | S984 | F120<br>G766<br>S967 | T<br>T<br>T | -<br>-<br>- | -<br>-<br>- | - |
| | D1154$_{TM12}$ | | T845$_{elbow2}$<br>**Y849$_{elbow2}$**<br>**R1030$_{TM10}$** | | E1004 | G697<br>Y701<br>R879 | T<br>**S**<br>**S** | -<br>-<br>- | Yes<br>Yes<br>Yes | - |
| | S1155$_{TM12}$ | | S1037$_{TM10}$<br>K1041$_{TM10}$<br>**Y1092$_{TM11}$** | | N1005 | S886<br>K890<br>F941 | T<br>T<br>**S** | Yes<br>Yes<br>- | -<br>-<br>- | - |
| | R1158$_{TM12}$ | | T845$_{elbow2}$<br>T848$_{elbow2}$<br>**Q1038$_{TM10}$**<br>Q1039$_{TM10}$<br>Q1042$_{TM10}$ | | I1008 | G697<br>A700<br>R887<br>D888<br>R891 | T<br>T<br>**S**<br>T<br>T | Yes<br>Yes<br>-<br>-<br>Yes | Yes<br>Yes<br>Yes<br>-<br>- | - |
| *<br>*<br>*<br>* | S1159$_{TM12}$ | S1159F<br>S1159P | D979$_{TM9}$<br>K1041$_{TM10}$<br>E1044$_{TM10}$<br>R1048$_{TM10}$ | D979V | S1009 | D828<br>K890<br>E893<br>R897 | T<br>T<br>T<br>T | Yes<br>Yes<br>Yes<br>Yes | -<br>-<br>-<br>- | - |
| | S1161$_{TM12}$ | | T848$_{elbow2}$<br>Q1042$_{TM10}$ | | E1011 | A700<br>R891 | T<br>T | Yes<br>Yes | -<br>- | Yes |
| *<br>*<br><br>* | R1162$_{TM12}$ | | N974$_{TM9}$<br>D979$_{TM9}$<br>Q1042$_{TM10}$<br>S1045$_{TM10}$<br>E1046$_{TM10}$ | | R1012 | N823<br>D928<br>R891<br>S894<br>T895 | T<br>T<br>T<br>T<br>T | Yes<br>Yes<br>Yes<br>Yes<br>Yes | -<br>-<br>-<br>-<br>- | Yes |
| <br>*<br>* | R1245$_{NBD2}$ | | E528$_{NBD1}$<br>D529$_{NBD1}$<br>D579$_{NBD1}$<br>T582 $_{NBD1}$ | (D579G) [II-III] | R1076 | K515<br>D516<br>D566<br>V569 | T<br>T<br>T<br>T | -<br>-<br>-<br>- | -<br>-<br>-<br>- | *NA* |
| *<br>* | T1246$_{NBD2}$ | (T1246I) | S549$_{NBD1}$<br><br>R555$_{NBD1}$ | S549N [III]<br>S549R [II-III] | T1077 | S536<br><br>R542 | T<br><br>T | -<br><br>- | -<br><br>- | *NA* |
| | N1262$_{NBD2}$ | | S962$_{CH3}$ | | E1093 | L811 | T | - | - | *NA* |
| | Q1280$_{NBD2}$ | | C276 $_{CH2}$<br>E278$_{TM5}$<br>Q958$_{CH3}$ | | H1111 | A290<br>E292<br>K897 | T<br>T<br>T | -<br>-<br>- | -<br>-<br>- | *NA* |
| *<br>* | R1283$_{NBD2}$ | R1283M | E278$_{CH2}$<br>**E1172$_{TMD2-NBD2}$** | | R1104 | E292<br>E1022 | T<br>**S** | -<br>- | -<br>- | *NA* |
| | Q1291$_{NBD2}$ | (Q1291H)<br>(Q1291R) | R553$_{NBD1}$<br>Y577$_{NBD1}$ | | Q1122 | K540<br>A564 | T<br>T | Yes<br>Yes | -<br>- | *NA* |
| | S1297$_{NBD2}$ | | E264$_{TM4}$ | | T1128 | E278 | T | - | - | *NA* |
| | D1341$_{NBD2}$ | | K174$_{CH1}$<br>K1060$_{TM10}$ | | E1172 | K188<br>Q909 | T<br>T | -<br>Yes | -<br>- | *NA* |
| | H1348$_{NBD2}$ | | T460$_{NBD1}$<br>Q493$_{NBD1}$ | | V1179 | V447<br>Q480 | T<br>T | -<br>- | -<br>- | *NA* |
| *<br>* | R1358$_{NBD2}$ | | Y275$_{CH2}$<br>W277$_{CH2}$ | | R1189 | Y289<br>W291 | T<br>T | -<br>- | -<br>- | *NA* |
| | H1375$_{NBD2}$ | H1375P | T460$_{NBD1}$<br>Q493$_{NBD1}$<br>S573$_{NBD1}$<br>Y577$_{NBD1}$ | | N1206 | V447<br>Q480<br>D560<br>A564 | T<br>T<br>T<br>T | -<br>-<br>Yes<br>Yes | -<br>-<br>-<br>- | *NA* |
